# ‘Dispersification’ of *Agalinis* (Orobanchaceae) into South America is associated with hummingbird pollination and perennial life history shifts

**DOI:** 10.1101/2025.11.21.689784

**Authors:** Pedro H. Pezzi, Pamela S. Soltis, Douglas E. Soltis, Maribeth Latvis

## Abstract

**Aim:** Several mechanisms contribute to the plant biodiversity of the Neotropics, with the highlands of South America serving as important hotspots of diversity. In particular, the Brazilian highlands exhibit high biodiversity due to complex diversification dynamics and a mixture of contributions from different biomes. In this study, we reconstruct the timing and potential triggers of diversification of *Agalinis*, hemiparasitic plants that inhabit open grassland habitats, to investigate their biogeographic history and migration patterns across the Americas.

**Location:** North, Central, and South America

**Taxon:** Agalinis.

**Methods:** We reconstructed dated phylogenies of *Agalinis* using a secondary calibration approach, sampling 73% of the known species, including multiple species from the Andes and Brazilian highlands. We inferred ancestral distributions to understand migration patterns between North and South America and within South America. Additionally, we investigated shifts in diversification rates within the genus and reconstructed ancestral pollination syndrome and life history strategy states. All analyses were performed across a distribution of trees to account for phylogenetic uncertainty.

**Results:** *Agalinis* likely originated in southeastern North America during the early Miocene and rapidly diversified, followed by movement into South America in the Late Miocene or Early Pliocene. We propose two possible routes for *Agalinis* movement into South America: either through the Andes or via the South American lowland grasslands (e.g., Chaco, Pampas, Cerrado, Caatinga, Llanos), using grassland corridors for dispersal within the continent. After its arrival in South American highlands, the clade underwent rapid diversification. State reconstructions indicated that the genus had a bee-pollinated ancestor and that hummingbird pollination evolved only once, with many transitions back to bee pollination. In contrast, the perennial life strategy evolved multiple times within the genus, including at least once in the ancestor of all South American species and twice in the North American species.

**Main Conclusions:** *Agalinis* likely originated in North America and later migrated to South America, followed by rapid diversification (i.e., ‘dispersification’). Thus, *Agalinis* is a clade that refutes the tropical conservatism hypothesis and the out-of-the-tropics model. Moreover, the dispersal to the Brazilian highlands was from nearby species pools and not from other highland habitats such as the Andes, followed by several *in situ* speciation events. These high diversification rates were partly associated with Quaternary climatic oscillations, and perennial and hummingbird-pollinated species exhibited higher turnover rates.

## 1. INTRODUCTION

The Neotropics harbor more than 100,000 species of seed plants, making it the most diverse region in the world (Gentry, 1982). This ecozone extends from Central Mexico to Argentina (Morrone, 2014), spanning several major biomes and encompassing important topographic and habitat heterogeneity, including many of the world’s biodiversity hotspots (Myers et al., 2000). Many theories have been proposed to explain the high biodiversity of the Neotropics (Antonelli & Sanmartín, 2011; Rull, 2011; Antonelli et al., 2018), involving a variety of factors such as adaptation to soil, temperature, pollinator shifts, dispersal ability, phylogenetic niche conservatism, and geological events. However, the relative importance of these factors has long been debated (reviewed in Antonelli & Sanmartín, 2011; Hughes et al., 2013; Antonelli et al., 2018), and evidence suggests that the staggering biodiversity in the region results from a mosaic of distinct, local-scale histories and processes that collectively contribute to large-scale patterns (e.g., Antonelli & Sanmartín, 2011; Drummond et al., 2012; Hughes et al., 2013; Antonelli et al., 2018).

One major pattern that has been repeatedly associated with Neotropical biodiversity is the uplift of the Andes, which served as an avenue for temperate-adapted lineages to migrate into the Neotropics and diversify via allopatric speciation and ecological shifts (Hughes & Eastwood, 2006; Drummond et al., 2012; Hughes et al., 2013). In addition to facilitating the movement of new lineages, the Andes created an elevational gradient with distinct microclimates and also affected the climate of the entire continent (Gregory-Wodzicki, 2000). Thus, the Andes acted as a major ‘species pump’, triggering diversification shifts in biomes elsewhere in the Neotropics and contributing to the overall biodiversity of the region (Antonelli et al., 2009; Hoorn et al., 2010; Sedano & Burns, 2010).

Moreover, several other geographical and climatic factors have contributed to the high biodiversity of the Neotropics, including the closure of the Panama Isthmus (Gentry, 1982), the Miocene marine incursions (Hoorn et al., 2022), and the Pleistocene glacial-interglacial cycles (Hewitt, 2000). Additionally, other montane regions, such as the Guiana Shield and formations on the Brazilian Shield—particularly the Serra do Espinhaço, Serra da Mantiqueira, and Serra do Mar—have played key roles in shaping Neotropical biodiversity. Despite their importance, these other highland systems in South America have received considerably less attention, and phylogenetic and biogeographic studies remain sparse (but see Givnish et al., 2011; Alcantara et al., 2018; Vasconcelos et al., 2020; Miola et al., 2021; Jantzen et al., 2022; Magri et al., 2025).

The Brazilian and Guiana Shields have different geologic histories compared to the Andes. They are much older, with formations dating back around 600 million and nearly 2 billion years, respectively. Due to continuous erosion, their elevations are also considerably lower than those of the Andes, reaching up to 2,000 and 2,900 meters, respectively. Within Brazil’s Serra do Espinhaço range, the *campos rupestres* (open rocky fields) stand out as a species-rich vegetation complex and have been recognized as a UNESCO World Heritage Site due to the high degree of endemism (Karez et al., 2012; Silveira et al., 2016). The vegetation of the *campos rupestres* is isolated from surrounding communities due to elevation (Alves & Kolbek, 1994) and edaphic differences, which may lead to ecophysiological adaptations to the soil (Alves et al., 2008). Such uniqueness has led scientists to propose the *campo rupestres* as a new and distinct biome of Brazil (Miola et al., 2021), separate from the surrounding biomes of the Cerrado, Atlantic Forest, and Caatinga. Despite its small geographical range (covering less than 1% of Brazilian territory), it contains about 15% of all Brazilian angiosperm species (Silveira et al., 2016).

The biodiversity of the Brazilian highlands investigated to date is likely shaped by complex diversification dynamics (Martinelli, 2007; Rapini et al., 2021). While *campos de altitude* and *campos rupestres* share similar landscapes and many plant genera and species, the former shows strong floristic affinities with other geographically distant highland systems such as the Andes, whereas the latter is more similar to Planalto Central, the Guiana Shield, and the lowland coastal restinga vegetation (Alves & Kolbek, 1994, 2010; DeForest Safford, 2007; Vasconcelos, 2011). Thus, the assembly of the Brazilian highlands may represent a mixture of contributions from nearby diverse species pools, as has been reported for lowland Cerrado (Simon et al., 2009). Another possibility is that it represents contributions from distant highlands, supported by the hypothesis that the Brazilian highlands were connected to the Andes through similar climatic and vegetative conditions during the Pleistocene glaciations, when cooler climates facilitated the formation of an open habitat corridor between the two (Behling, 1998; Ledru et al., 1998; Ledru & Araújo, 2023). Following the paradigm proposed by Donoghue (2008), “it is easier [for lineages] to move than to evolve (unless it isn’t)”, the latter scenario would be a more likely explanation for historical community assembly in the Brazilian highlands.

Most endemic lineages of the *campos rupestres* have diversified within the past 5 million years (Vasconcelos et al., 2020), with much of this diversification attributed to Plio-Pleistocene climatic oscillations, which created highland interglacial refugia, fragmented populations, and hindered gene flow (e.g., Ribeiro et al., 2014; Barres et al., 2019; Dantas-Queiroz et al., 2021, 2023). Thus, the accumulation of lineages was likely driven by *in situ* diversification in response to this changing climate, rather than from migration of pre-adapted lineages from other areas, which had been suggested as the dominant paradigm due to a combination of dispersal ability and a high degree of niche conservatism over evolutionary time (Crisp et al., 2009). Thus, a deeper understanding of possible historical movement between Neotropical mountains and, consequently, how the floristic assembly of such regions was formed requires knowledge of the historical biogeography and diversification rates within diverse and densely sampled clades (Edwards & Donoghue, 2013).

The genus *Agalinis* Raf. (Orobanchaceae) is a New World genus of about 70 species with a distribution extending from Nova Scotia, Canada, south through Argentina and Chile. Following the recommendations of Latvis et al. (2024) based on phylogenetic study of *Agalinis* and closely allied genera, *Esterhazya* was transferred to *Agalinis* (Souza et al., 2025). All species in the genus are hemiparasitic (Elias et al., 2001), meaning they are capable of photosynthesis but depend on host plants for water and some nutrients. Although little is known about the identity or specificity of their host plants, species of *Agalinis* have been found growing on a variety of host species within their respective communities (P. H. Pezzi and M. Latvis, personal observation). Approximately 35 *Agalinis* species occur in Central and South America, with centers of diversity in the puna communities of the Peruvian and Bolivian Andes, and also southeastern Brazil, with many species restricted to the fragmented Brazilian highlands in the state of Minas Gerais (Canne-Hilliker, 1988; Souza et al., 2001). *Agalinis* occurs disjunctly between North and South America and the two South American centers of diversity. A densely sampled phylogeny of *Agalinis* (Latvis et al., 2024), encompassing species from all its distribution, provides an excellent opportunity to examine patterns of historical movement and diversification between North and South America, and within the Neotropics. Moreover, this group allows us to evaluate how shifts in pollination and life history strategies may influence biodiversity patterns in the Neotropics. South American species exhibit a diversity of pollinators, with flowers adapted to pollination by bees and hummingbirds and display both perennial and annual life history strategies (Canne-Hilliker, 1988; Latvis et al., 2024). The hummingbird pollination syndrome is believed to have evolved at least twice in South America (Latvis et al., 2024). The group also includes annual and perennial species in both South and North America, making it ideal for exploring transitions between life history strategies and how they can influence diversification.

In this study, we placed *Agalinis* lineages in geographical space and evolutionary time to reconstruct their biogeographic history and better understand the migration routes used to reach the South American highlands. The clade also presents an opportunity to investigate how shifts in pollination syndrome and life history may be coupled with movement into these novel environments, contributing to diversification. We also addressed two hypotheses: (i) *Agalinis* species from the Brazilian highlands originated from distantly located highlands with similar climate and ecological conditions, such as the Andes, rather than from nearby biomes such as the Cerrado and Atlantic Forest; and (ii) the South American species experienced higher diversification rates than North American species, a dynamic associated with new pollination syndromes and life history strategies.

## 2. MATERIAL AND METHODS

### 2.1 Dated phylogeny

We used data from Latvis et al. (2024) and employed three methodological approaches to construct a time-calibrated phylogeny. Due to the lack of Orobanchaceae fossils, we calibrated our target tree with the divergence times estimated by Fonseca (2021) for the Lamiales phylogenetic tree, which included 842 genera, constructed using 10 genetic markers and calibrated with 12 fossils. To account for phylogenetic uncertainty, we employed distinct calibration strategies in each of the three approaches used to date our target tree, as detailed below.

As our first strategy, we applied the congruification method (Eastman et al., 2013) to map a reference tree, the Orobanchaceae dated phylogeny (pruned from Fonseca, 2021), onto our target, the uncalibrated *Agalinis* tree, which comprised 52 *Agalinis* taxa and three outgroup species. We used the PATHd8 algorithm (Britton et al., 2007) to scale branch lengths to time. For our second approach, we used a penalized likelihood (PL; Sanderson, 2002) approach implemented in r8s 1.81 (Sanderson, 2003) to date the Bayesian topology resulting from the concatenated plastid + nuclear pruned *Agalinis* dataset (Latvis et al., 2024). We fixed the age of the Most Recent Common Ancestor (MRCA) of *Agalinis* and *Aureolaria pedicularia* to 24.07 million years ago (Mya), the MRCA of *Aureolaria pedicularia* and *Brachystigma wrightii* to 8.39 Mya, and the MRCA of *Aureolaria pedicularia* and *Dasistoma macrophylla* to 3.22 Mya.

For our third approach, we employed BEAST 2.7.5 (Bouckaert et al., 2019) to reconstruct a dated phylogeny using the alignments from Latvis et al. (2024). We generated phylogenetic trees for a combined dataset that included six plastid DNA (cpDNA) regions (*rbcL*, *matK*, *trnT*(UGU)-*trnF*(GAA), *rps2*, *rpoB*, and *psbA*-*trnH*) and four nuclear regions (ITS, *PPR-AT1G09680*, *PPR-AT3G09060*, and *PPR-AT5G39980*). We also constructed separate trees for the nuclear and plastid datasets as Latvis et al. (2024) reported cytonuclear discordance. We selected the best substitution model available in BEAST for each partition using bModelTest (Bouckaert & Drummond, 2017). We used the Yule tree prior and the Optimised Relaxed Clock model. The crown age of *Agalinis* was constrained to 20.25 Mya (Fonseca, 2021), with normal distribution and a sigma of 1.0 to allow uncertainty in time estimation. We ran BEAST for three independent MCMC chains, each consisting of 100 million generations, sampling every 1,000 steps. We combined the runs with LogCombiner (Bouckaert et al., 2019), setting a burn-in of 25%, and assessed parameter convergence with Tracer version 1.7.2 (Rambaut et al., 2018). We generated Maximum Clade Credibility (MCC) trees for the three datasets using TreeAnnotator (Bouckaert et al., 2019).

### 2.2. Phylogenetic uncertainty

The importance of incorporating phylogenetic uncertainty (i.e., variations in topology and branch lengths) into biogeographic analyses has been demonstrated by Ceccarelli et al. (2018) and Magalhaes et al. (2021). One strategy to handle these uncertainties is by creating a small set of median trees from clusters of similar trees, which effectively condenses numerous trees that are difficult to visualize into a smaller, more manageable set. For this purpose, we randomly selected 10% of the trees of the postburn-in posterior trees to identify 10 clusters of similar trees using treespace (Jombart et al., 2017), retaining the first three principal components. We then calculated the median tree of each cluster using the treeVec algorithm (Kendall & Colijn, 2016).

However, the clustering of trees approach has faced some criticism as it may fail to represent the full distribution of trees (Smith, 2022). Thus, an alternative approach involves randomly sampling a larger number of trees from the posterior trees and summarizing the results. For this approach, we selected 100 random trees from the 50% MCMC post-burn-in posterior trees that were at least 1000 apart from each other. This approach has also faced criticism, primarily regarding the reconstruction of ancestral biogeographic regions (Matzke, 2019), as it summarizes the results of several trees into one, similar to what an MCC tree does. For this reason, we decided to continue with both strategies. All subsequent analyses were run on 15 trees, derived from the congruification method, the r8s method, the three full MCC trees (concatenated, plastid, and nuclear datasets), and the 10 median trees estimated using treespace, as well as on the 100 randomly selected trees.

### 2.3 Ancestral range estimation

To address our question regarding the origin of the Brazilian highlands floristic assembly, we considered six geographic areas for *Agalinis*: A) Central and Eastern North America; B) arid southwestern North America; C) Caribbean; D) Andes; E) lowland Cerrado, Chaco, and Pampas; F) *campos rupestres* and *campo de altitude sensu latu* (Brazilian highlands). These areas were coded as discrete, multistate characters for each species of *Agalinis*, *Esterhazya* (now included in *Agalinis*), and outgroup taxa *Aureolaria pedicularia*, *Brachystigma wrightii*, and *Dasistoma macrophylla* based on a combination of species distributions in Global Biodiversity Information Facility (GBIF.org, 2024), knowledge of collaborators, and fieldwork experience. The major geographic areas were delimited based on vegetational zones and geologic history while minimizing the total number of areas and polymorphic distributions for coded species but allowing for ranges spanning more than one of the six geographic areas when necessary. Lowland ‘open vegetation’ biomes (i.e., Cerrado, Chaco, Pampas; see Fiaschi & Pirani, 2009; Werneck, 2011) were coded together for this study, although there are notable floristic and climatic differences between them (Furley, 1999; Cardoso Da Silva & Bates, 2002; Werneck, 2011; Jaramillo, 2023). Although considered subtypes of Cerrado vegetation in most classifications, *campos rupestres* and *campos de altitude*, communities that are both restricted to high elevations (rocky outcrops and grasslands, respectively), were coded together and separately from the Cerrado biome, given the goals of our study.

We used BioGeoBEARS 1.1.3 package (Matzke, 2013) in R 4.3.3 (R Core Team, 2024) using the Dispersal-Extinction-Cladogenesis (DEC; Ree & Smith, 2008), Dispersal-Vicariance Analysis (DIVAlike; Ronquist, 1997), and Bayesian Inference of Historical Biogeography for Discrete Areas (BAYAREAlike; Landis et al., 2013) models. We set the maximum possible range size to three areas. Additionally, we ran these same models allowing for ‘jump’ dispersal (+j parameter), wherein cladogenesis events can be associated with range dispersal and expansion. Ree & Sanmartín (2018) reported statistical issues with models that allow for ‘jump dispersal’ events. However, Matzke (2022) compared these models and concluded that they are valid and may perform better than models without the +j parameter. Therefore, we chose to keep these models. We selected the best model based on the Akaike Information Criterion (AIC; Akaike, 1973). We pruned *A. acuta* and *A. paupercula* from the tree because they are synonyms of *A. decemloba* and *A. purpurea*, respectively. Although *A. albida* is recognized as a synonym of *A. harperi*, we chose not to remove it from the tree because we did not recover them as sister taxa (see Results below). For the dataset of 100 random trees, we ran the DEC+j model, which was the best model for the MCC tree (see Results below), using the ‘run_bears_optim_on_multiple_trees’ function (Matzke, 2019). We summarized the results by averaging the probabilities across all trees for nodes also present in the MCC concatenated tree.

### 2.4 Diversification rates

We used BAMM 2.5.0 (Rabosky, 2014) to estimate tip diversification rates and identify rate shifts in *Agalinis*. We pruned the outgroup and synonym species from the tree and set the sampling fraction to 0.71 for the Andean clade, 0.58 for the Brazilian clade, 0.84 for the North American clade, and 0.66 for the sister clade of the South American species (*A. heterophylla* + *A. calycina* + *A. densiflora*, the latter of which was not sampled). Priors were set to recommendations from the BAMMtools 2.1.11 package (Rabosky et al., 2014), and we ran four chains for 50 million generations, sampling every 5000 generations. Chain convergence was assessed by examining the effective sample sizes (ESS) with the coda 0.19-4.1 package (Plummer et al., 2006) after discarding 20% of the MCMC runs as burn-in. We used the ‘getBestShiftConfiguration’ function to determine the maximum a posteriori probability of shift scenario and ‘getRateThroughTimeMatrix’ to calculate the diversification rate of *Agalinis* over time.

We also investigated *Agalinis* diversification rates using MiSSE (Vasconcelos et al., 2022) implemented with the hisse 2.1.12 package (Beaulieu & O’Meara, 2016). MiSSE is a State-dependent Speciation and Extinction (SSE) model that does not depend on traits. We set the overall sampling fraction to 0.72 and identified the set of models to run with the function ‘MiSSEGreedy’, allowing for both turnover and extinction fraction to vary. We tested a maximum of 52 parameters and stopped the search when deltaAICc > 10. We pruned the redundant models with the function ‘PruneRedundantModels’ and averaged the tip diversification rates of the models with similar AIC using the ‘GetModelAveRates’ function. For the dataset with 100 random trees, we calculated the average tip net diversification (λ - μ) and turnover (λ + μ) for BAMM and MiSSE, respectively.

### 2.5 Trait reconstruction

We used corHMM 2.8 (Beaulieu et al., 2013; Boyko & Beaulieu, 2021) to estimate pollination syndromes and life history strategies (i.e., annual versus perennial) at the nodes of *Agalinis* across the phylogenetic trees. Because effective pollinators are not known for many species, we classified them as bee- or hummingbird-pollinated by examining flower shape, tube length, corolla color, and stamen position, as a combination of traits provides a more reliable prediction of pollinators than a single trait alone (Dellinger, 2020). More specifically, we coded flowers with a combination of red color, exserted stamens, a tubular shape with a long corolla tube, and the absence of a landing platform as indicative of hummingbird pollination syndrome, and flowers with pink or purple color, inserted stamens, a bell-shaped form with a short corolla tube, and the presence of a landing platform as indicative of bee pollination syndrome. This classification aligned with established floral syndromes and known pollinators documented for both North American (Dieringer & Cabrera Rodriguez, 2024) and South American *Agalinis* species (Canne-Hilliker, 1988; Rodrigues & Rodrigues, 2014; Ferreira et al., 2016). We coded species as annual or perennial for life history strategies based on their descriptions, observations, and occurrence records.

Ancestral trait reconstruction methods assume that the phylogenetic tree accurately reflects the diversity of the group, and any bias in the tree can affect the results (Duchêne & Lanfear, 2015). Because hummingbird-pollinated species were overrepresented in the Andean clades (see Results), we adjusted the proportion of pollination syndromes in the phylogeny as in Emberts & Wiens (2021). For the pollination syndrome reconstruction analysis, we pruned *A. bangii*, *A. stenantha*, and *A. reflexidensis* to maintain the proportion of hummingbird-pollinated species at approximately 50%, while also preserving the number of pollination transitions in the phylogeny.

We ran corHMM models using simple Markov models (SMMs) and hidden Markov models (HMMs). We applied the Yang rooting method for all models, and for HMM models, we used two hidden states. We created models allowing transition rates to be either equal (SYM) or different (ARD). Similar to Sazatornil et al. (2023), we calculated the model-averaged marginal probabilities using corrected Akaike Information Criterion (AICc) weights from the eight corHMM models for each of the 15 trees. For the dataset with 100 random trees, we used the same approach to average the likelihood of bee and hummingbird pollination as well as annual and perennial strategies for the ancestral node of the South American clade in each tree and reported the percentage of trees supporting each type of pollination and life history strategy for that node.

### 2.6 Pollination syndrome, life history strategies, and turnover rates

We used the hisse R package to explore whether pollination mode (i.e., bee versus hummingbird pollination) and life history strategy (i.e., annual versus perennial) were associated with distinct turnover rates. We compared models with constant diversification rates to a Binary State Speciation and Extinction (BiSSE; Maddison et al., 2007) model. Additionally, we ran three Hidden State Speciation and Extinction (HiSSE; Beaulieu & O’Meara, 2016) models to account for hidden factors potentially influencing diversification rates in addition to pollination syndromes and life history strategies. We also ran two character-independent models (CID), which do not consider these traits, with two (CID-2) and four (CID-4) hidden states. We kept extinction parameters linked for all models and set the sampling fractions to 0.71 and 0.58 for bee-pollinated and hummingbird-pollinated species, respectively, and to 0.77 and 0.68 for annual and perennial species, respectively. Because SSE models assume random sampling (FitzJohn et al., 2009), we pruned the tree for the SSE analysis of pollination syndrome similarly to the ancestral trait reconstruction. The best model was selected based on AICc.

We also investigated whether turnover rates associated with different pollination modes are influenced by life history strategy using the multistate HiSSE model (MuHiSSE; Nakov et al., 2019). We used the pruned phylogeny for pollination mode to avoid overrepresenting hummingbird-pollinated species, consisting of 49 taxa. Our character state combinations included: bee-pollinated and annual (00), bee-pollinated and perennial (01), hummingbird-pollinated and annual (10; no species were present in this category, as all hummingbird-pollinated species are perennials), and hummingbird-pollinated and perennial (11), along with the hidden states A and B. We tested eight MuHiSSE models under different assumptions: (1) constant turnover rates across all categories, without hidden states (Constant rate); (2) turnover varying between annual and perennial species (NoHidden1); (3) turnover varying between bee-and hummingbird-pollinated species (NoHidden2); (4) turnover varying among all combinations of life history and pollination mode (NoHidden3); (5) turnover varying between annual and perennial species and hidden states, but constrained to pollination syndromes (MuHiSSE1); (6) turnover varying between bee- and hummingbird-pollinated species and hidden states, but constrained to life history strategies (MuHiSSE2); (7) turnover varying among all categories, with hidden states A and B (MuHiSSE3); and (8) a null model in which turnover rates are associated only with hidden states, independent of life history strategies and pollination modes (MuCID2). We set the sampling fractions to 0.77 for bee-pollinated annual species, 0.56 for bee-pollinated perennial species, 1.0 for hummingbird-pollinated annual species, and 0.58 for hummingbird-pollinated perennial species. As in the standard SSE models, we linked extinction parameters across all models and selected the best-fitting model using AICc.

## 3. RESULTS

### 3.1 Divergence Time Estimation within *Agalinis*

Because we used the same sequences and phylogenies as Latvis et al. (2024), the relationships among species and clades followed the patterns discussed in that study. As we applied a secondary calibration approach to our tree, fixing the age of the *Agalinis* clade at 20.25 Mya based on the results of Fonseca (2021), the origin of *Agalinis* was consistent across methods; however, the ages of clades within the genus varied (Figure 1; Table S1). The MCC BEAST concatenated tree indicated that the origin of the group in North America occurred at 18.57 Mya (95% highest posterior density [HPD]: 16.16–20.93 Mya), with the major subclades within North America diverging over a short span of approximately 3 million years. However, the other clades were much younger: the crown age of the *A. heterophylla* + *A. calycina* + South American clade was estimated at 6.18 Mya (95% HPD: 4.01–8.63 Mya), the South American clade at 4.92 Mya (95% HPD: 3.31–6.68 Mya), the Brazilian clade at 4.29 Mya (95% HPD: 2.85–5.85 Mya), and the Andean clade at 2.51 Mya (95% HPD: 1.49–3.66 Mya). The congruification and r8s methods estimated similar ages for the North American clade, 18.28 and 19.55 Mya, respectively, but both methods indicated an older divergence time compared to the MCC BEAST concatenated tree for the *A. heterophylla* + *A. calycina* + South American clade (8.60 and 12.78 Mya for congruification and r8s, respectively), the South American clade (6.08 and 9.40 Mya), and the Brazilian clade (5.02 and 8.52 Mya).

**Figure 1.**
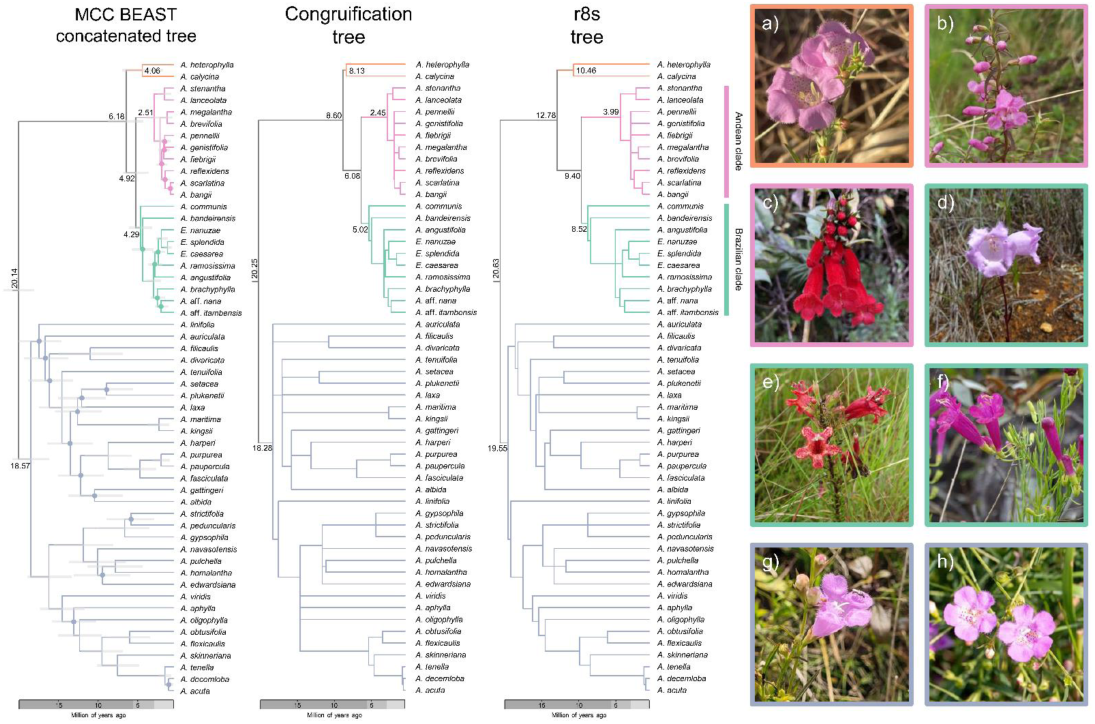
Dated phylogenies of *Agalinis* based on three calibration methods: BEAST secondary calibration (left), congruification (center), and r8s penalized likelihood (right). The BEAST Maximum Clade Credibility (MCC) tree was built using concatenated nuclear and plastid markers. The circles on nodes of the BEAST MCC concatenated tree indicate posterior probability of < 90%. The congruification and r8s trees were calibrated using the *Agalinis* phylogeny from Latvis et al. (2024) and a pruned Lamiales fossil-calibrated tree from Fonseca (2021). Ages of important nodes for our discussion are shown in the trees. Floral diversity within *Agalinis* is illustrated in panels a–h, with borders colored to match the clades in the phylogenies: a) *A. heterophylla* (iNaturalist 246468527, photo by jankolk), b) *A. megalantha* (iNaturalist 194505173, photo by J. R. Kuethe [@jrkuethe]), c) *A. fiebregii* (iNaturalist 260449073, photo by Gonzalo Martinez [@gonza_martinez35]), d) *A. bandeirensis* (photo by Pedro H. Pezzi), e) *Esterhazya splendida* (photo by Pedro H. Pezzi), f) *A. angustifolia* (photo by Pedro H. Pezzi), g) *A. fasciculata* (photo courtesy of Leonardo T. Gonçalves), and h) *A. maritima* (iNaturalist 254840475, photo by Holly Greening [@hsgzoe]). Photos a–c and h are sourced from iNaturalist.com and used under Creative Commons licenses, with observation numbers and contributor usernames in parentheses.

### 3.2 Biogeographic inference

DEC models with and without the +j parameter were the best fit for all trees (Table S2). The MCC BEAST concatenated tree inferred the origin of *Agalinis* in region AB (Figure 2a), while all trees inferred region A or a combination of areas including A as the most likely area of origin (Figures 2b–d, S1). However, the ancestral biogeographic region of the clade comprising the South American species + *A. heterophylla* + *A. calycina* showed greater uncertainty, suggesting multiple regions as possible ancestral areas and variation between trees. The MCC BEAST concatenated tree indicated an origin as Central and Eastern North America (Figure 2a); however, other trees suggested different combinations of regions as the possible ancestral region for the group (Figures 2b–d, S1). Regarding the ancestor of the South American clade, which includes the Brazilian and Andean clades, the models also suggest uncertainty across the different trees tested: the MCC concatenated BEAST tree indicated the Andean region as the ancestral area of the clade, a result supported by 12 other trees (those in which region D, or a combination of areas including D, had a probability greater than 20%). The second most likely region was E, or a combination of areas including E, supported by 10 trees. For the Brazilian clade, the trees suggest an ancestral range in the Cerrado, Pampas, Chaco regions, and the Brazilian highlands, and if the lowlands were occupied first, migration to the highlands occurred soon after the lowlands.

**Figure 2.**
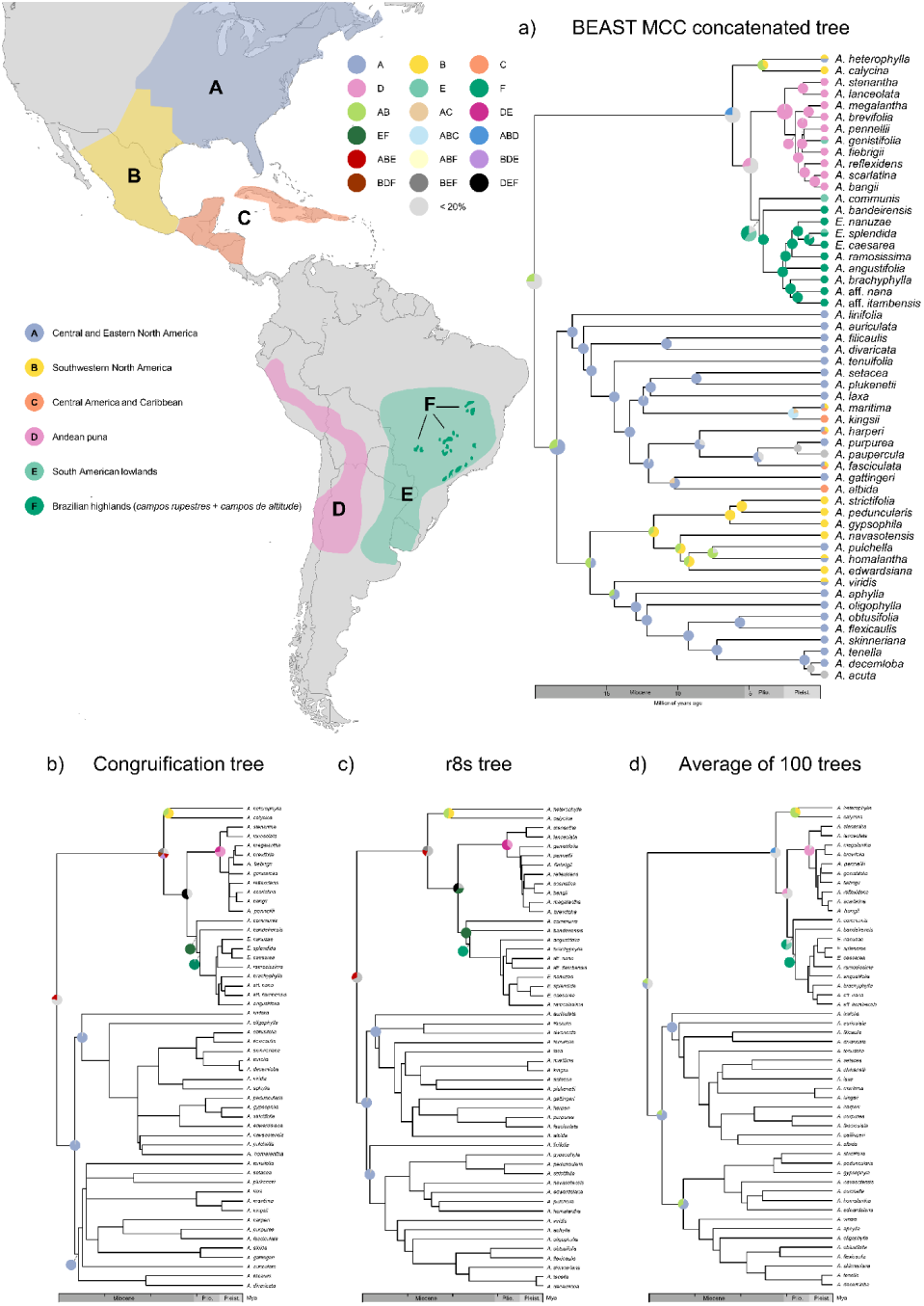
Biogeographic ancestral range reconstruction for *Agalinis* using BioGeoBEARS. Ancestral range estimations are shown for: a) the BEAST Maximum Clade Credibility (MCC) concatenated tree under the DEC+J model, b) the congruification tree under the DEC model, c) the r8s penalized likelihood tree under the DEC model, and d) averaged probabilities across 100 randomly sampled post–burn-in MCMC trees under the DEC+J model, plotted onto the BEAST MCC tree. All analyses included the outgroups *Aureolaria pedicularia*, *Dasistoma macrophylla*, and *Brachystigma wrightii*, which were pruned here for clarity. Circles at the tips of the BEAST MCC tree indicate the current geographic ranges of extant species. Pie charts at internal nodes represent ancestral area probabilities or combinations of areas, showing only those with probabilities >20%; if no area exceeds this threshold, areas with probabilities >15% are shown. Gray circles on the BEAST MCC tree indicate pruned taxa considered synonymous with their sister species. Model log-likelihood values for each tree are summarized in Table S1.

### 3.3 Diversification shift and ancestral pollination syndromes and life strategies

The BAMM analysis on the MCC BEAST concatenated tree suggested a diversification rate shift at the stem of the South American species + *A. heterophylla* + *A. calycina*, with tip net diversification rates higher in these species compared to the North American species (Figure 3A). This result was also observed in almost all trees as 10 of the other 14 trees indicated a rate shift in the same clade, and 79 of the 100 trees supported this finding (Figure S2). The net diversification through time followed a consistent pattern across all trees: it was higher near the origin of the genus, decreased until the origin of the South American clade, and then increased, although it never reached the same level as at the genus origin (Figure S3). MiSSE found a similar pattern to BAMM, suggesting higher turnover rates in the South American species (Figures 3A and S2).

**Figure 3.**
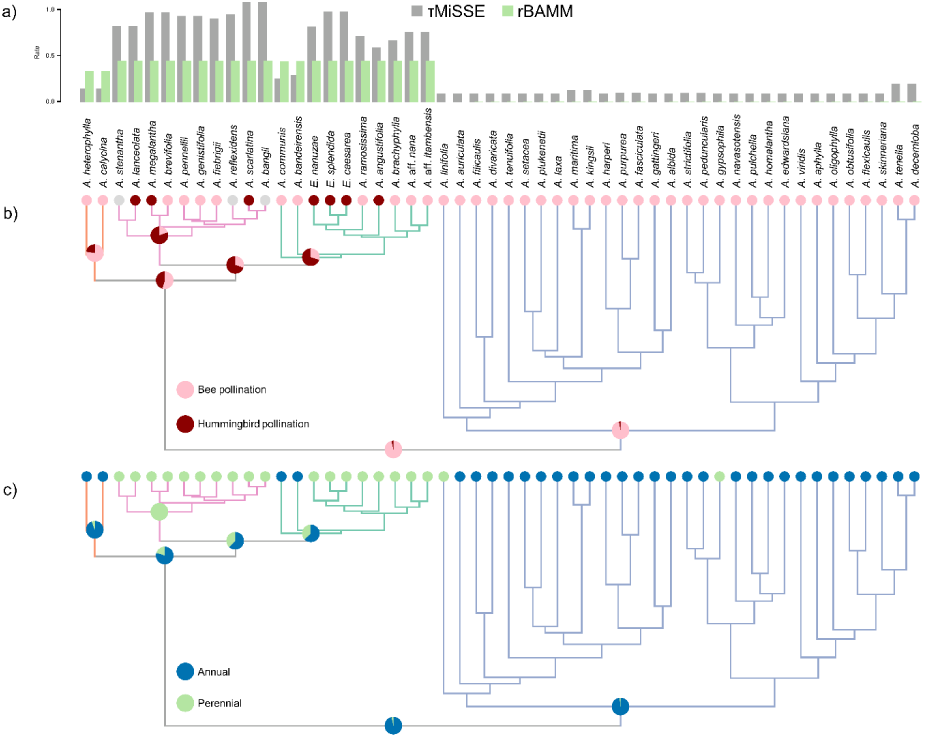
a) Tip diversification rates in *Agalinis* estimated using BAMM (rBAMM; green) and tip turnover rates estimated using MiSSE (τMiSSE; gray). b) Ancestral reconstruction of pollination syndrome using corHMM at key nodes, with marginal probabilities averaged across eight AICc-weighted models. Gray tip circles indicate species pruned from the analysis to maintain a proportion of sampled bee- and hummingbird-pollinated species that reflects their natural frequencies in the Andean clade. For a reconstruction including results for all nodes, see Figure S4. (c) Ancestral reconstruction of life history strategy using corHMM at key nodes, with marginal probabilities averaged across eight AICc-weighted models. For a reconstruction including results for all nodes, see Figure S5.

The reconstruction of pollination syndrome for all trees indicated that the ancestor of the genus had a bee-pollination syndrome (Figures 3B and S4), as did all nodes within the North American clade. Most trees suggested that the ancestor of the South American clade + *A. heterophylla* + *A. calycina* had a bee-pollination syndrome, but the ancestral nodes of the South American, Andean, and Brazilian clades were reconstructed as hummingbird-pollinated. Among the 100 trees, 84% reconstructed the node of the South American clade as having the highest probability of being hummingbird-pollinated. This suggests that the origin of hummingbird pollination occurred only once in the evolutionary history of *Agalinis* but with several transitions back to bee pollination. Regarding life history strategies, the ancestor of *Agalinis* was annual (Figures 3C and S5). However, in contrast to the pattern observed in the reconstruction of pollination syndromes, the perennial life history strategy evolved multiple times: twice within the North American clade and twice in the South American clade, with independent origins in the Andean and Brazilian clades. Among the 100 trees, 73 indicated that the ancestor of the South American clade was annual.

Our SSE analysis revealed that both pollination type and life history strategy are strongly associated with turnover rates. BiSSE consistently showed the lowest AIC across all trees, with a delta AICc > 10 compared to the second-best model in the MCC BEAST concatenated tree (Table 1), as well as in most of the trees (Tables S3–S6). Additionally, HiSSE was the second-best model in all trees. The results showed that hummingbird-pollinated and perennial species have turnover rates multiple times higher than that of bee-pollinated and annual species, respectively (Figure 4A–B). When pollination syndrome and life history strategy were combined to assess diversification, the model in which turnover rates were associated only with pollination syndrome (NoHidden2) showed the lowest AICc for the MCC BEAST concatenated tree (Table 3). Most trees corroborated this result (Tables S7–S8), and hummingbird-pollinated species consistently showed higher turnover rates than bee-pollinated species (Figure 4C). However, the delta AICc between this model and the model in which each category without hidden states had its own turnover rate (NoHidden3) was very small for the MCC BEAST concatenated tree, and NoHidden3 was identified as the best-fitting model in 15 of the 100 randomly sampled trees (Table S8). In the NoHidden3 model, the lowest turnover rates were observed in bee-pollinated annuals, while the other three categories showed similar turnover rates (Figure 4D).

**Figure 4.**
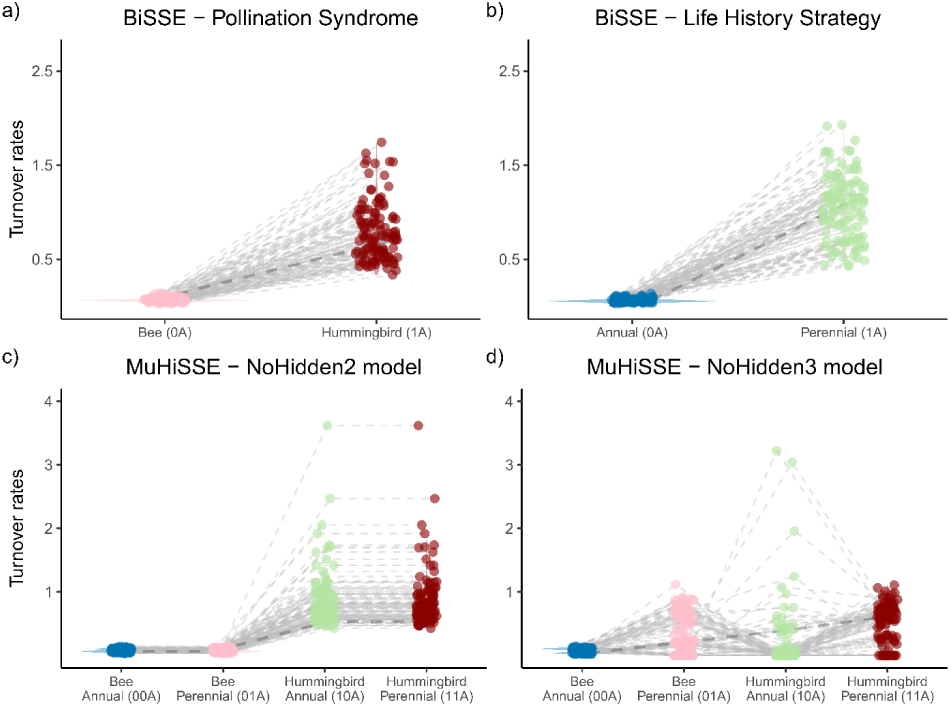
Turnover rates in *Agalinis* estimated using State Speciation and Extinction (SSE) models across all trees. a) Turnover rates by pollination syndrome and b) by life history strategy, estimated under the Binary SSE (BiSSE) model for each trait. BiSSE results indicated that hummingbird-pollinated species exhibit higher turnover rates than bee-pollinated species, and that perennial species have higher turnover rates than annuals. Turnover rates combining pollination syndrome and life history strategy under MuHiSSE for models c) NoHidden2 and d) NoHidden3. Both models are shown due to a low delta AIC between them. The NoHidden2 model associates turnover only with pollination syndrome, not life history strategy, and shows higher turnover rates for hummingbird-pollinated species. The NoHidden3 model estimates distinct turnover rates for each trait combination and indicates overall lower turnover for bee-pollinated annual species. Dashed lines in all graphs indicate turnover rates estimated from the BEAST MCC concatenated tree.

**Table 1.** Log-likelihood values (lnL), Akaike Information Criterion (AIC), and corrected AIC (AICc) for different State Speciation and Extinction (SSE) diversification models associating pollination syndrome and life history strategy in *Agalinis*, based on the BEAST MCC concatenated tree. The Binary SSE (BiSSE) and Hidden SSE (HiSSE) models link diversification rates to the traits, whereas the constant rate model and CID2 and CID4 models assume diversification is independent of these traits. For pollination syndrome, 0 indicates bee-pollinated species and 1 indicates hummingbird-pollinated species. For life history strategy, 0 indicates annual species and 1 indicates perennial species. In both analyses, A–D denote hidden states.

|  | Model | lnL | AIC | AICc | turnover<br>0A | turnover<br>1A | turnover<br>0B | turnover<br>1B | turnover<br>0C | turnover<br>1C | turnover<br>0D | turnover<br>1D |
| --- | --- | --- | --- | --- | --- | --- | --- | --- | --- | --- | --- | --- |
| Pollination<br>Syndrome | Constant rate | -163.23 | 334.47 | 335.38 | 0.14 | 0.14 | NA | NA | NA | NA | NA | NA |
|  | **BiSSE** | **-146.84** | **303.68** | **305.08** | **0.07** | **0.71** | **NA** | **NA** | **NA** | **NA** | **NA** | **NA** |
|  | HiSSE | -146.23 | 312.46 | 318.25 | 0.00 | 0.00 | 0.10 | 0.52 | NA | NA | NA | NA |
|  | CID2 | -155.85 | 323.70 | 325.70 | 0.00 | 0.00 | 0.95 | 0.95 | NA | NA | NA | NA |
|  | CID4 | -152.68 | 321.36 | 324.96 | 1.00 | 1.00 | 0.03 | 0.03 | 0.03 | 0.03 | 0.03 | 0.03 |
| Life History<br>Strategy | Constant rate | -172.36 | 352.73 | 353.58 | 0.15 | 0.15 | NA | NA | NA | NA | NA | NA |
|  | **BiSSE** | **-155.81** | **321.62** | **322.92** | **0.06** | **1.10** | **NA** | **NA** | **NA** | **NA** | **NA** | **NA** |
|  | HiSSE | -153.20 | 326.40 | 331.77 | 0.04 | 0.27 | 0.00 | 0.57 | NA | NA | NA | NA |
|  | CID2 | -162.51 | 337.02 | 338.88 | 0.00 | 0.00 | 1.28 | 1.28 | NA | NA | NA | NA |
|  | CID4 | -159.32 | 334.63 | 337.98 | 1.17 | 1.17 | 0.04 | 0.04 | 0.04 | 0.04 | 0.04 | 0.04 |

## 4. DISCUSSION

In this study, we investigated continental and local-scale processes contributing to biodiversity patterns in the Neotropical highlands, particularly the Brazilian highlands in southeastern Brazil, by estimating divergence times for *Agalinis* (Orobanchaceae) and reconstructing its historical biogeography and diversification. We used data from a densely sampled *Agalinis* phylogeny (Latvis et al., 2024) and a dated Lamiales phylogeny (Fonseca, 2021), employing biogeographic, diversification, and trait reconstruction methods to evaluate scenarios of lineage movement of *Agalinis* in space and time, while accounting for phylogenetic uncertainty, i.e., the distinct evolutionary relationships (topology) and divergence times (branch lengths) among trees. Our analyses suggest that the present-day distribution of *Agalinis* results from recent dispersal from North America into South America via southwestern North America. However, we found conflicting results regarding the ancestral area of South American lineages, suggesting that dispersal may have occurred either using the Andes as a corridor or across South American lowland grassland regions, followed by separate subsequent dispersals and radiations into Neotropical highlands (Brazilian highlands and the Andes). Our results also suggest that *Agalinis* arrived in the Brazilian highlands through a dispersal event from nearby species pools rather than the Andes, rapidly followed by repeated *in situ* speciation events. The dispersal of *Agalinis* into South America was accompanied by an increase in turnover rates, which were associated with shifts from bee to hummingbird pollination and from an annual to a perennial life strategy.

### The Origin of *Agalinis* and Movement into South America

We found that *Agalinis* originated 18 to 22 Mya (Table S1), during the early Miocene, in Central-Eastern North America. As we used a secondary calibration approach, the placement of the origin of *Agalinis* in the early Miocene was consistent across all trees. The early Miocene was characterized by the slow expansion of grasslands in the continental interior of North America (reviewed in Steinthorsdottir et al., 2021). The climate started to warm around 20 Mya, followed by the Middle Miocene Climatic Optimum (MMCO) between 17 and 14 Mya, a period of rapid global cooling and aridification (Zachos et al., 2001). This time frame was marked by the beginning of aridification in North America (Eronen et al., 2012), the expansion of forests along the East and West coasts, and the dominance of grasslands in the central regions of the continent (Steinthorsdottir et al., 2021). The North American clade is estimated to have originated just before the MMCO, around 18 Mya, in Central-Eastern North America. The clade then rapidly diversified during the MMCO, as indicated by the shorter branches near its crown. Neel & Cummings (2004) and Pettengill & Neel (2008) previously suggested a rapid radiation scenario in North America due to poorly supported nodes along the backbone of this clade and short internal branch lengths. Given that *Agalinis* is commonly found in open areas, grasslands, and savannas throughout its range, mostly occurring in well-drained soil, this radiation could be associated with the fragmentation of grasslands due to forest expansion and/or the dispersal to new grass-dominated areas across the continent. A study focused on North American *Agalinis* species supports a Central-Eastern North America ancestral area, specifically the southeastern USA, although it estimated a much younger crown age for the group, ranging from 4.9 to 12.24 Mya (Roy et al., 2021). The discrepancy in divergence time estimates for *Agalinis* (Roy et al., 2021; Yu et al., 2018) and related groups (Xu et al., 2022) compared to our study can be attributed to the lack of fossils in Orobanchaceae and the dependence on secondary calibrations.

Roy et al. (2021) adopted divergence times from Yu et al. (2018), who calibrated their phylogeny using four fossils. Similarly, Xu et al. (2022) dated their tree based on estimates from Xu et al. (2019), which incorporated three fossil calibrations within Lamiales. Thus, our study represents improved taxon sampling of *Agalinis* and fossil constraints in Lamiales (Fonseca, 2021).

Our dated phylogenies indicated a broad range for the crown age of the South American clade + *A. heterophylla* + *A. calycina*, estimated between 12 and 6 Mya, with a widespread ancestor occupying more than one geographical area. Regarding the divergence of the crown clade of South American *Agalinis* from its sister lineage, most phylogenetic trees support a divergence time of 5–6 Mya, during the Late Miocene and Early Pliocene. During this time, the vegetation in South America was characterized by the expansion of drier, grass-dominated habitats, primarily driven by the rise of C4 grasses (Edwards et al., 2010) and increased aridity (Amidon et al., 2017). Because *Agalinis* species are hemiparasites and likely parasitize graminoids (P. H. Pezzi and M. Latvis, personal observation), the movement into South America and their higher diversification rates in the region could also be linked to the expansion of such grassland habitats. Although no studies have properly identified their hosts or examined host specificity in *Agalinis*, one can expect the biogeographic history of parasitic plants to be associated with the biogeography and diversification of their hosts. Range expansion and establishment in new areas may occur through co-diversification with their hosts (Santiago-Rosario et al., 2024) or through adaptation to new hosts (Martos et al., 2023). Including information on host-parasite relationships in future biogeographic studies will provide valuable insights into the biogeographic patterns and diversification history of *Agalinis* and other parasitic clades.

If we consider younger estimates of the closure of the Isthmus of Panama, which suggest it was completed around 3 Mya (O’Dea et al., 2016), the migration from North to South America took place before its completion. However, this scenario changes if we consider the earlier estimates of the closure of the Isthmus of Panama (Bacon et al., 2015; Jaramillo, 2018), in which expansion from North to South America would have taken place after its closure. In either case, this North to South America migration pattern contrasts with the scenario envisioned by Pennell (1928, 1929), who suggested that the major lineages in North America represented recent invasions from the Neotropics. The closure of the Isthmus of Panama was a defining event in Neotropical biogeography as it facilitated the migration of lineages between the Americas, a process known as the Great American Biotic Interchange (Stehli & Webb, 1985; Webb, 1991).

However, many dated plant phylogenies suggest earlier divergence time estimates for historical migration events across the Isthmus of Panama, before the Pliocene (e.g., Fritsch et al., 2015; Freitas et al., 2016; Leal et al., 2022), indicating that the lack of a land bridge was a weaker barrier to plant dispersal compared to animal dispersal (Cody et al., 2010), or that the closure of the Isthmus took place before 3 Mya and facilitated migration of plants, including *Agalinis*, between North and South America.

While most of our trees agreed on the divergence time of the South American clade around the Late Miocene to Early Pliocene, reconstruction of the biogeographic ancestral area indicated uncertainty about the route *Agalinis* took to reach South America. This uncertainty was due to disagreements among different phylogenies and widespread ancestral distribution across multiple areas. In the first possible scenario, the Andes, a well-known North-South corridor (e.g., Luebert et al., 2009; Luebert & Weigend, 2014), may have facilitated the movement of *Agalinis* into South America. After arriving in the Andes, *Agalinis* rapidly diversified within the mountains and dispersed to other regions of South America, including the southeastern highlands of Brazil, where it experienced a second radiation. Although the Andes is an ancient mountain range dating back to approximately 200 Mya, it underwent major uplifts during the Late Miocene and Early Pliocene, around 8–5 Mya (reviewed in Pérez-Escobar et al., 2022), attaining its current heights within the last 10 million years. This uplift facilitated the diversification of high-elevation species (Madriñán et al., 2013; Luebert & Weigend, 2014; Hughes, 2016) and the origin of the páramo ecosystem (Garzione et al., 2008; Graham, 2011) and may have contributed to diversification within *Agalinis*. Climatic fluctuations during the Pleistocene were associated with evolutionary radiations in the region through a process known as flickering connectivity (Flantua et al., 2019), in which populations experienced repeated cycles of geographical isolation and connectivity. Flantua et al. (2019) also hypothesized that species experienced varying levels of flickering connectivity, which may now be reflected in the genetic diversity of *Agalinis*.

All of our dated phylogenies indicated a slightly older crown age for the Brazilian clade compared to the Andean clade, with concomitant rapid and recent radiations in both clades. Considering this, along with the results of the biogeographic ancestral area reconstruction, a second scenario for the expansion into South America is through either the lowland grassland biomes or the southeastern highlands of Brazil, rather than via the Andean corridor. Given that the ancestors of the South American + *A. calycina* + *A. heterophylla* clade likely resembled extant North American species inhabiting open grasslands, and considering that the Brazilian highlands represent a small geographic area within the South American open grassland biomes (biogeographic area E), it is reasonable to propose that the lowland grassland biomes—such as the Caatinga, Cerrado, Pampas, Llanos, and Chaco—offered a more plausible route for *Agalinis* than the Brazilian highlands, followed shortly after by a rapid movement into the *campos rupestres*, *campos de altitude*, and the Andes. Although we present strong alternative scenarios for the dispersal to and within South America, we unfortunately did not sample *A. glandulosa*, which occupies the Central-West region of Brazil. Its placement in the phylogeny could help clarify the migration history with greater resolution, as it currently occupies the intermediate region between two major biodiversity hotspots, the Andes and the Brazilian highlands.

Although the migration route *Agalinis* used to reach South America remains uncertain, the consistent phylogenetic placement of *A. communis*, a widespread species from lowland grasslands in southern Brazil, as sister to the rest of the Brazilian clade (Figure 1), along with the reconstructed ancestral biogeographic area of the clade (Figure S1), suggests that lineage accumulation of *Agalinis* in the Brazilian highlands started with the migration of a lineage from surrounding lowland biomes rather than the Andes, followed by multiple *in situ* speciation events. The timing of these *in situ* speciation events coincides with the Late Pliocene-Pleistocene climatic oscillations, which fragmented populations and hindered gene flow by flickering connectivity (Vasconcelos et al., 2020; Dantas-Queiroz et al., 2021; Jantzen et al., 2022; Magri et al., 2025), a similar mechanism found in the Andes (Flantua et al., 2019). Additionally, a slight uplift of the Precambrian Brazilian Plateau along the Serra do Espinhaço, Serra da Mantiqueira, and Serra do Mar 2–4 Mya has been proposed to influence diversification in Bromeliaceae (Givnish et al., 2011). A combination of these climatic and geologic factors likely contributed to the diversification of *Agalinis* during this time. Moreover, other plant species have shown patterns of dispersal to highlands followed by rapid radiation, accompanied by edaphic (Scarano, 2007; Alves et al., 2008; Alcantara et al., 2018) and reproductive (Drummond et al., 2012) adaptations. Although the hummingbird pollination syndrome is estimated to have originated before its establishment in the Brazilian highlands (Figure 3), improved sampling and the use of high-throughput sequencing data could help determine whether these adaptations evolved once or multiple times within the genus and clarify their role in the successful occupation of the highlands in southeastern Brazil.

The migration history of *Agalinis* and its subsequent dispersal between the Andes and Brazilian highlands aligns with the well-documented grassland expansion during the Miocene. This relatively recent movement, along with the simultaneous expansions into two distinct highland regions, suggests the existence of corridors of suitable habitat during that period. The assembly of biodiversity in the Andes is often attributed to exchanges with the Amazon, but with important contributions from other biomes, including the Cerrado (Antonelli et al., 2018). There is evidence of a connection between the Andes and the Atlantic Forest, as well as the *campos rupestres* in the Cerrado biome, through Bolivia, where grasslands reached the foothills of the Andes (reviewed in Ledru & Araújo, 2023), dating back to the Pliocene (Batalha-Filho et al., 2013). Additionally, the crown ages of Andean and Brazilian *Agalinis* clades, which coincide with the spread of grasses in the Andean highlands and the establishment of the Brazilian Cerrado (Kirschner & Hoorn, 2020), suggest that these habitat corridors facilitated the expansion of *Agalinis* across South America.

### Diversification of *Agalinis* in South America - ‘Dispersification’, Pollinator Shifts, and New Life History Strategies

BAMM analysis indicated a diversification rate shift in the South American clade + *A. heterophylla* + *A. calycina* (Figure 3A), with tip net diversification rates higher than North American species. These results were corroborated by the high tip turnover rates inferred by MiSSE for the South American clade (Figure 3A). This increase in net diversification and turnover coincides with movement into a new biogeographic realm, providing evidence of a recent ‘dispersification’ event (Moore & Donoghue, 2007) into the Neotropics, a pattern already observed in other plant groups (e.g., Hughes & Eastwood, 2006; Uribe-Convers & Tank, 2015; Vieu et al., 2022). The high diversification of Neotropical *Agalinis* is likely associated with adaptations to abiotic factors, such as different climate conditions due to the elevational gradient they occupy, although it requires further investigation. Moreover, *Agalinis* likely represents yet another clade refuting the tropical conservatism hypothesis, which attributes high tropical biodiversity to origin and persistence in the tropics for longer periods due to phylogenetic niche conservatism and predicts that temperate lineages will more often be younger, less diverse, and derived from tropical ones (Wiens & Donoghue, 2004; Mittelbach et al., 2007; Moore & Donoghue, 2007; Donoghue, 2008). It also refutes the out-of-the-tropics model (Jablonski et al., 2006), which states that most temperate taxa have a tropical origin, as we observe the opposite pattern in *Agalinis*. Net diversification over time revealed similar patterns across all trees, peaking at the origin of the genus around 20 Mya and coinciding with the North American radiation, then declining until about 5 Mya, marking the ‘dispersification’ event and the rapid radiations of the Andean and Brazilian clades. However, it is crucial to interpret these diversification estimates with caution, as we could not use fossils, which could potentially mislead these estimates (Matos-Maraví, 2016).

Dispersal to new environments is not the only factor associated with higher diversification in *Agalinis*. Our results indicated that hummingbird-pollinated and perennial species exhibit an increase in turnover rates compared to bee-pollinated and annual species, respectively (Figure 4A–B). The relationship between hummingbird pollination and diversification rates is complex, with no universal pattern (reviewed in Barreto et al., 2024). As observed here, some studies have shown higher speciation rates in clades that evolved hummingbird pollination (e.g., Lagomarsino et al., 2016; Serrano-Serrano et al., 2017), while in other clades, the hummingbird pollination syndrome was considered a dead-end evolutionary transition, leading to decreased diversification rates (e.g., Wessinger et al., 2019; Siniscalchi et al., 2023). As vertebrate pollinators are more efficient in mountainous regions (Cruden, 1972), especially in tropical areas (Dellinger et al., 2023), and hummingbirds, in particular, perform better than bees in tropical high-elevation habitats, which are often cold, rainy, and foggy (Stiles, 1978), this pattern may partially explain why the hummingbird pollination syndrome is associated with higher turnover rates in *Agalinis*, and why it has evolved in the South American clade but not in North American species. Moreover, hummingbirds are more common in the Brazilian highlands than in the surrounding biomes (Carstensen et al., 2014), but their distribution across the mountains is not uniform (Rodrigues & Rodrigues, 2014; Monteiro et al., 2021). This suggests that *Agalinis* diversification could be a result of local pollinator communities, with different pollination syndromes affecting population connectivity and genetic diversity, making *Agalinis* a good model for exploring microevolutionary processes across the landscape in association with pollination syndromes.

The hummingbird pollination syndrome has evolved multiple times across at least 22 families, with bee pollination being the most common ancestral state (Barreto et al., 2024). Ancestral pollination syndrome reconstruction for *Agalinis* suggests that the genus was originally bee-pollinated, which aligns with the distribution and timeline of hummingbird evolution, as modern hummingbirds only reached North America after the origin and radiation of *Agalinis* in the region, around 12 Mya (McGuire et al., 2014; Licona-Vera & Ornelas, 2017). In contrast, corHMM analysis of the South American species indicated that the ancestor of the South American clade was already hummingbird-pollinated, implying that the hummingbird pollination syndrome evolved only once in the genus, followed by several reversions to bee pollination. Transitions back to bee pollination were once considered rare (Barrett, 2013); however, a recent review showed that they occur with similar frequency to the transition from bee to hummingbird pollination (Barreto et al., 2024). When *Agalinis* arrived in South America, modern hummingbirds were already established in the region, having originated there approximately 22 Mya (McGuire et al., 2014). This suggests that the hummingbird pollination syndrome evolved rapidly following the movement of *Agalinis* into South America. However, these findings should be interpreted with caution, as confirming how many times the hummingbird pollination syndrome evolved will require a more comprehensive sampling, ideally including all South American species, as well as genomic studies targeting genes associated with hummingbird-adapted floral traits and phylogenetic analyses that account for potential adaptive introgression.

High diversification rates are often associated with multiple factors. For example, in Bromeliaceae, diversification rates were linked to traits such as an epiphytic lifestyle, tank formation, and adaptation to high elevations (Kessler et al., 2020), while in Campanulaceae, they were associated with temperature and the Andean uplift (Lagomarsino et al., 2016). In *Agalinis*, life history strategies were also associated with differences in turnover rates, with perennial species showing higher turnover than annual species (Figure 4B). Annuality occurs in more than 120 plant families and has evolved multiple times (Friedman, 2020). This trait is typically derived from a perennial lifestyle, which is considered the ancestral condition in plants. Although reversals from annual to perennial have been reported (Bena et al., 1998; Soltis et al., 2013), including in another genus of Orobanchaceae, *Castilleja* (Tank & Olmstead, 2008), such transitions are considered rare. However, in *Agalinis*, whose ancestral state is annual, the perennial life strategy has evolved multiple times within the group (Figure 3), twice in South America and twice in North America. The ancestral state of *Agalinis* is unlikely to change with further sampling, although including more South American species could affect estimates of how often life history strategies have shifted. The genetic mechanisms underlying life history strategy shifts are not fully understood, but they probably involve a large gene regulatory network (Hjertaas et al., 2023). This complexity challenges our finding that the Brazilian and Andean clades evolved the perennial lifestyle independently (Figure 3). One possibility is that perenniality evolved independently in the Brazilian and Andean clades due to pre-adaptations in the ancestor of South American *Agalinis*, which would have facilitated the transition from annual to perennial in both radiations.

Higher turnover in perennial and hummingbird-pollinated species is supported by the MuHiSSE results (Table 2; Figures 4C–D). The low delta AICc values among models NoHidden2 (turnover rates associated with pollination syndromes), NoHidden3 (unique turnover rates for each trait combination), and in some cases NoHidden1 (turnover rates associated with life history strategy) indicate low support for selecting a single best model (Tables S7–S8). This suggests that both perenniality and hummingbird pollination contribute to high turnover rates in *Agalinis*. Additionally, hummingbird pollination is found only in perennial species, a pattern also reported in *Castilleja* (Tank & Olmstead, 2008), indicating that these traits may be linked and that perenniality likely evolved before the hummingbird pollination syndrome, challenging our trait reconstruction findings. Therefore, *Agalinis* represents a valuable opportunity to investigate the genetic mechanisms underlying transitions in life history strategies and pollination syndromes and their association with diversification rates.

**Table 2.** Log-likelihood values (lnL), Akaike Information Criterion (AIC), and corrected AIC (AICc) for Multistate Hidden State Speciation and Extinction (MuHiSSE) models combining pollination syndrome and life history strategy with diversification in *Agalinis*, based on the BEAST MCC concatenated tree. Categories are bee-pollinated annual species (00), bee-pollinated perennial species (01), hummingbird-pollinated annual species (10), and hummingbird-pollinated perennial species (11). The constant rate and MuCID2 models assume diversification is independent of these traits. Models NoHidden1, NoHidden2, and NoHidden3 do not include hidden states, while MuHiSSE1, MuHiSSE2, and MuHiSSE3 include hidden states A and B. Model NoHidden1 associates diversification with life history strategy, NoHidden2 associates diversification with pollination syndrome, and NoHidden3 assigns unique turnover rates to each category. The corresponding MuHiSSE models include hidden states for each of these trait combinations.

| Model | lnL | AIC | AICc | turnover<br>00A | turnover<br>01A | turnover<br>10A | turnover<br>11A | turnover<br>00B | turnover<br>01B | turnover<br>10B | turnover<br>11B |
| --- | --- | --- | --- | --- | --- | --- | --- | --- | --- | --- | --- |
| Constant rate | -175.86 | 371.73 | 377.52 | 0.14 | 0.14 | 0.14 | 0.14 | NA | NA | NA | NA |
| NoHidden1 | -164.39 | 350.78 | 357.91 | 0.10 | 0.54 | 0.10 | 0.54 | NA | NA | NA | NA |
| **NoHidden2** | **-159.06** | **340.11** | **347.25** | **0.06** | **0.06** | **0.53** | **0.53** | **NA** | **NA** | **NA** | **NA** |
| NoHidden3 | -157.08 | 340.16 | 350.56 | 0.03 | 0.20 | 0.41 | 0.61 | NA | NA | NA | NA |
| MuHiSSE1 | -159.17 | 346.34 | 358.69 | 0.15 | 0.15 | 0.83 | 0.83 | 0.00 | 0.00 | 0.00 | 0.00 |
| MuHiSSE2 | -160.01 | 348.02 | 360.37 | 0.14 | 0.00 | 0.14 | 0.00 | 0.00 | 0.57 | 0.00 | 0.57 |
| MuHiSSE3 | -155.69 | 363.38 | 427.20 | 0.04 | 0.26 | 0.00 | 0.35 | 0.00 | 0.00 | 0.00 | 0.73 |
| MuCID2 | -168.20 | 360.40 | 369.07 | 0.10 | 0.10 | 0.10 | 0.10 | 0.48 | 0.48 | 0.48 | 0.48 |

## 5. CONCLUSIONS

Here, we inferred the historical biogeographic and diversification history of *Agalinis* using a densely sampled, time-calibrated phylogeny, accounting for phylogenetic uncertainty. Our results indicate that *Agalinis* originated in southeastern North America during the Early Miocene and dispersed into South America in the Late Miocene to Early Pliocene. We propose two potential migration routes that *Agalinis* may have used to reach South America, using grassland corridors to disperse between mountainous regions within the continent. We also observed an increase in diversification rates in *Agalinis* following movement from North America into the Neotropics, thus providing another example that refutes the tropical conservatism hypothesis and the out-of-the-tropics model. We further suggest that very recent radiation within the Brazilian highlands and the Andes was driven, in part, by expansion and contractions of open habitats during the Quaternary. In addition to the effects of ‘dispersification’, we found that the transition from bee to hummingbird pollination syndromes and from annual to perennial life history strategy also played an important role in the diversification of South American species.

Altogether, these findings show the importance of multiple local and regional-scale processes working in concert to contribute to the overall biodiversity of the Neotropics, particularly in the understudied Brazilian highlands region.

## ACKNOWLEDGMENTS

This research is supported by the Arkansas High Performance Computing Center which is funded through multiple National Science Foundation grants and the Arkansas Economic Development Commission. This research was funded by an NSF DDIG grant (DEB-1310863) awarded to D. Soltis and M. Latvis, an NSF Graduate Research Fellowship to M. Latvis, a Lewis and Clark grant for field research, and graduate research awards from The Botanical Society of America, American Society of Plant Taxonomists, Society for Systematic Biologists, and Explorers’ Club.

## CRediT AUTHOR CONTRIBUTION

PHP: Data curation, Formal Analysis, Investigation, Software, Visualization, Writing – original draft, Writing – review & editing; PSS: Conceptualization, Funding acquisition, Investigation, Supervision, Writing – review & editing; DES: Conceptualization, Funding acquisition, Investigation, Supervision, Writing – review & editing; ML: Conceptualization, Data curation, Formal Analysis, Funding acquisition, Investigation, Supervision, Writing – original draft, Writing – review & editing.

## DATA AVAILABILITY STATEMENT

All code used for the analyses and the resulting data are available at GitHub https://github.com/pedrohpezzi/AgalinisBiogeography.git and Zenodo https://zenodo.org/records/15758305?preview=1&token=eyJhbGciOiJIUzUxMiJ9.eyJpZCI6ImIzNzBhN2EwLWExM2YtNGRlMS1hN2UzLTM5YzA0MjM4MjE1OCIsImRhdGEiOnt9LCJyYW5kb20iOiJiMjJlNjA4ZjBhNWQ2NDliNzE0ZGY4ODg2ZjM4YTlmZiJ9.PutODN3aTHzf4qktjhSIPU0zXxXVgHFjg1iVSW3sbST6Vez8ry8LAwH40wUZ5zatLnkii37qVtmnl2u8G92mgg.

## LIST OF TABLES AND FIGURES

**Figure S1.** Biogeographic ancestral range reconstruction for key nodes of *Agalinis* using BioGeoBEARS, based on the BEAST Maximum Clade Credibility (MCC) trees for nuclear and plastid datasets, as well as Cluster 1–10 trees. The best-fitting biogeographic model and associated likelihood scores for each tree are summarized in Table S1. Outgroups (*Aureolaria pedicularia*, *Dasistoma macrophylla*, and *Brachystigma wrightii*) were included in all analyses but are pruned from the figure for clarity. Pie charts at internal nodes represent ancestral area probabilities or combinations of areas, showing only those with probabilities >20%.

**Figure S2.** Tip diversification rates in *Agalinis* estimated using BAMM (rBAMM; green) and tip turnover rates estimated using MiSSE (τMiSSE; gray) across the congruification tree, r8s tree, BEAST MCC nuclear and plastid trees, the average of 100 randomly sampled trees, and Cluster 1–10 trees.

**Figure S3.** *Agalinis* diversification through time estimated using BAMM for the BEAST MCC concatenated tree, the congruification tree, the r8s tree, the BEAST MCC nuclear and plastid trees, and Cluster 1–10 trees.

**Figure S4.** Ancestral reconstruction of *Agalinis* pollination syndrome using corHMM, with marginal probabilities averaged across eight AICc-weighted models for the BEAST MCC concatenated tree, the congruification tree, the r8s tree, the BEAST MCC nuclear and plastid trees, and Cluster 1–10 trees.

**Figure S5.** Ancestral reconstruction of *Agalinis* life history strategy using corHMM, with marginal probabilities averaged across eight AICc-weighted models for the BEAST MCC concatenated tree, the congruification tree, the r8s tree, the BEAST MCC nuclear and plastid trees, and Cluster 1–10 trees.

**Table S1.** Divergence time estimates for major *Agalinis* clades across different phylogenetic analyses.

**Table S2.** Corrected Akaike Information Criterion (AICc) values for biogeographic models tested with BioGeoBEARS across multiple phylogenetic trees of *Agalinis*: the BEAST MCC concatenated tree, congruification tree, r8s tree, BEAST MCC nuclear and plastid trees, and Cluster 1–10 trees. The models evaluated include DEC, DIVALIKE, and BAYAREALIKE, also including the variation of each model allowing for ‘jump’ dispersal (+j). For each tree, the best-fitting model is indicated in bold.

**Table S3.** Log-likelihood values (lnL), Akaike Information Criterion (AIC), and corrected AIC (AICc) for State Speciation and Extinction (SSE) models associating pollination syndrome and diversification in *Agalinis*. Results are presented for the congruification tree, r8s tree, BEAST MCC nuclear and plastid trees, and Cluster 1–10 trees. The Binary SSE (BiSSE) and Hidden SSE (HiSSE) models link diversification rates to pollination syndrome, while the constant rate, CID2, and CID4 models assume diversification is independent of this trait. In the analyses, 0 denotes bee-pollinated species, 1 denotes hummingbird-pollinated species, and letters A–D indicate hidden states.

**Table S4.** Log-likelihood values (lnL), Akaike Information Criterion (AIC), and corrected AIC (AICc) for State Speciation and Extinction (SSE) models associating life history strategy and diversification in *Agalinis*. Results are presented for the congruification tree, r8s tree, BEAST MCC nuclear and plastid trees, and Cluster 1–10 trees. The Binary SSE (BiSSE) and Hidden State SSE (HiSSE) models link diversification rates to life history strategies, while the constant rate, CID2, and CID4 models assume diversification is independent of this trait. In the analyses, 0 denotes annual species, 1 denotes perennial species, and letters A–D indicate hidden states.

**Table S5.** Log-likelihood values (lnL), Akaike Information Criterion (AIC), and corrected AIC (AICc) for State Speciation and Extinction (SSE) models associating pollination syndrome and diversification in *Agalinis* for the 100 randomly sampled post–burn-in MCMC trees. The Binary SSE (BiSSE) and Hidden State SSE (HiSSE) models link diversification rates to pollination syndrome, while the constant rate, CID2, and CID4 models assume diversification is independent of this trait.

**Table S6.** Log-likelihood values (lnL), Akaike Information Criterion (AIC), and corrected AIC (AICc) for State Speciation and Extinction (SSE) models associating life history strategy and diversification in *Agalinis* for the 100 randomly sampled post–burn-in MCMC trees. The Binary SSE (BiSSE) and Hidden State SSE (HiSSE) models link diversification rates to life history strategy, while the constant rate, CID2, and CID4 models assume diversification is independent of this trait.

**Table S7.** Log-likelihood values (lnL), Akaike Information Criterion (AIC), and corrected AIC (AICc) for Multistate Hidden State Speciation and Extinction (MuHiSSE) models combining pollination syndrome and life history strategy with diversification in *Agalinis* for the congruification tree, r8s tree, BEAST MCC nuclear and plastid trees, and Cluster 1–10 trees. Categories are bee-pollinated annual species (00), bee-pollinated perennial species (01), hummingbird-pollinated annual species (10), and hummingbird-pollinated perennial species (11). The constant rate and MuCID2 models assume diversification is independent of these traits. Models NoHidden1, NoHidden2, and NoHidden3 do not include hidden states, while MuHiSSE1, MuHiSSE2, and MuHiSSE3 include hidden states A and B. Model NoHidden1 associates diversification with life history strategy, NoHidden2 associates diversification with pollination syndrome, and NoHidden3 assigns unique turnover rates to each category. The corresponding MuHiSSE models include hidden states for each of these trait associations.

**Table S8.** Log-likelihood values (lnL), Akaike Information Criterion (AIC), and corrected AIC (AICc) for Multistate Hidden State Speciation and Extinction (MuHiSSE) models combining pollination syndrome and life history strategy with diversification in *Agalinis* for the 100 randomly sampled post–burn-in MCMC trees. The constant rate and MuCID2 models assume diversification is independent of these traits. Models NoHidden1, NoHidden2, and NoHidden3 do not include hidden states, while MuHiSSE1, MuHiSSE2, and MuHiSSE3 include hidden states A and B. Model NoHidden1 associates diversification with life history strategy, NoHidden2 associates diversification with pollination syndrome, and NoHidden3 assigns unique turnover rates to each category. The corresponding MuHiSSE models include hidden states for each of these trait associations.

