## Supplementary figures and images for "‘Dispersification’ of *Agalinis* (Orobanchaceae) into South America is associated with hummingbird pollination and perennial life history shifts"

### Figure S1.

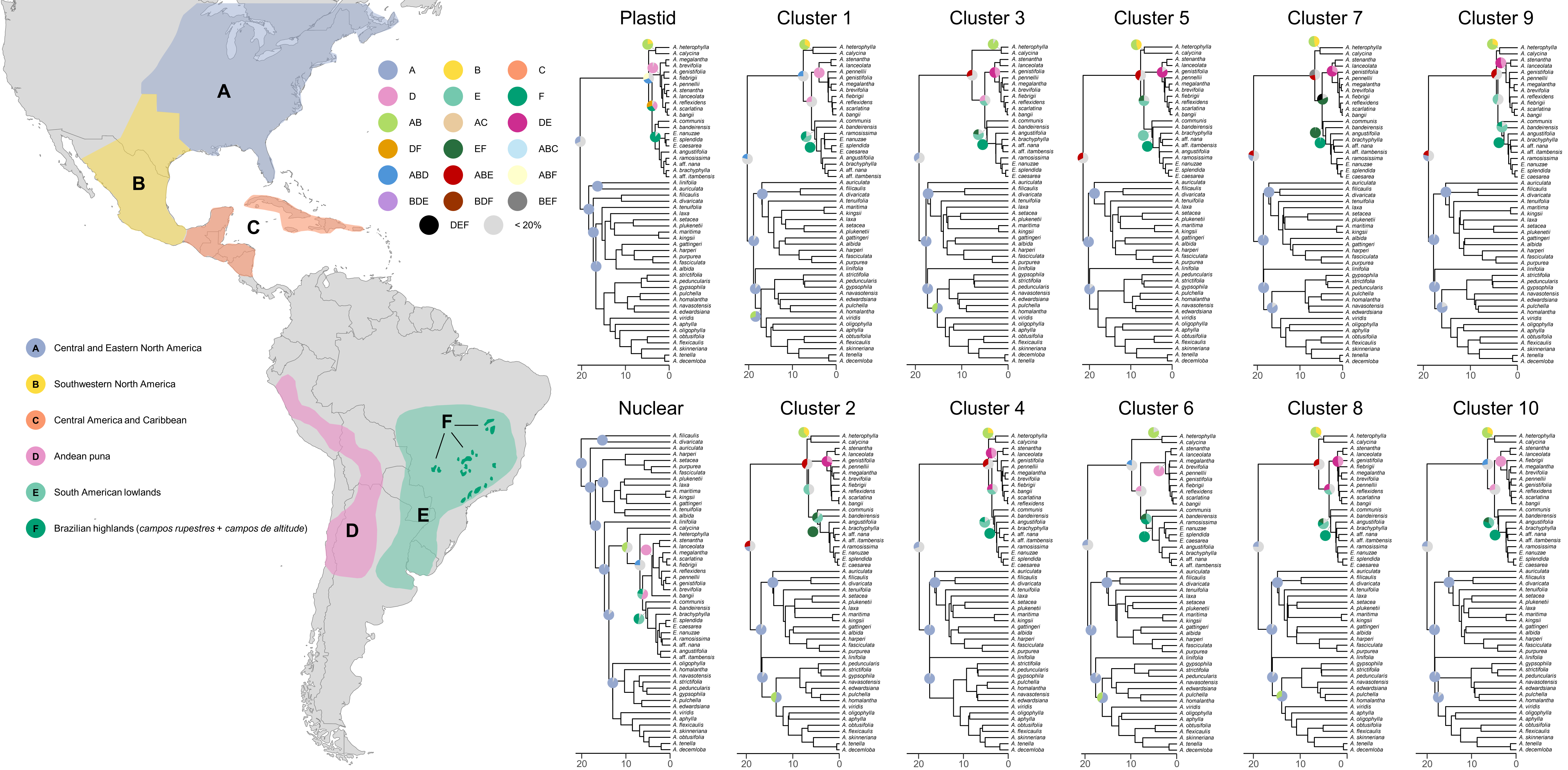

### Figure S2.

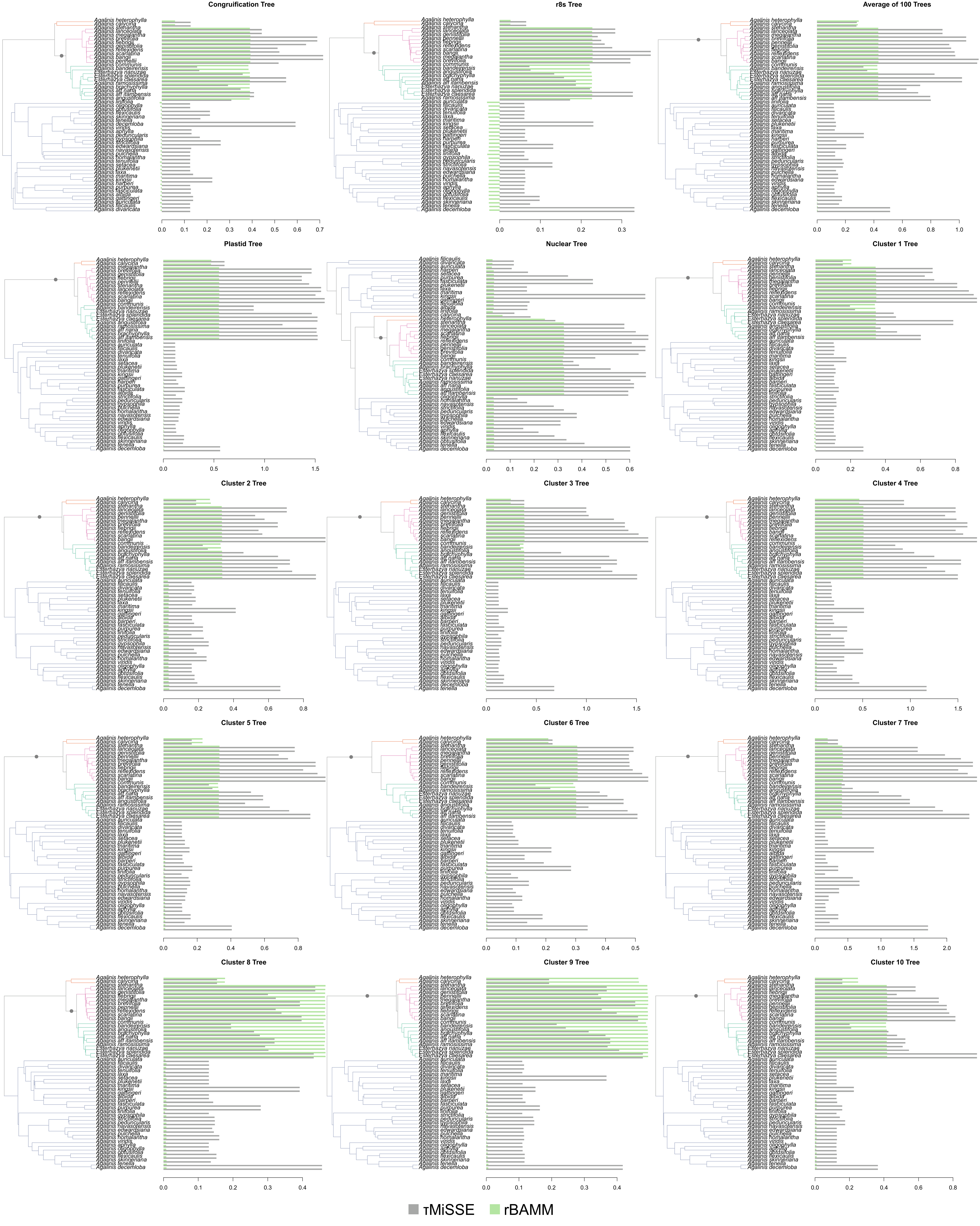

### Figure S3.

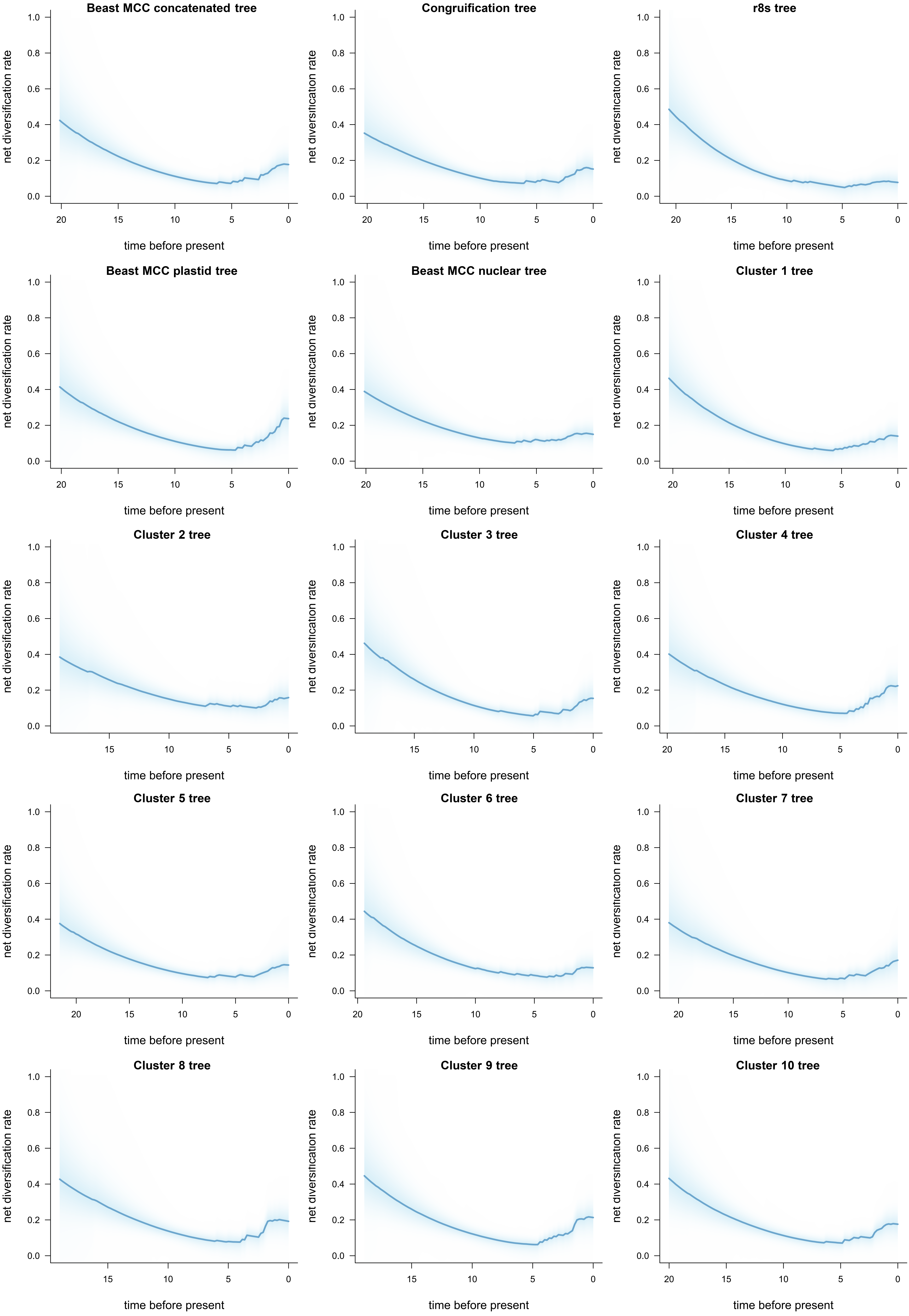

### Figure S4.

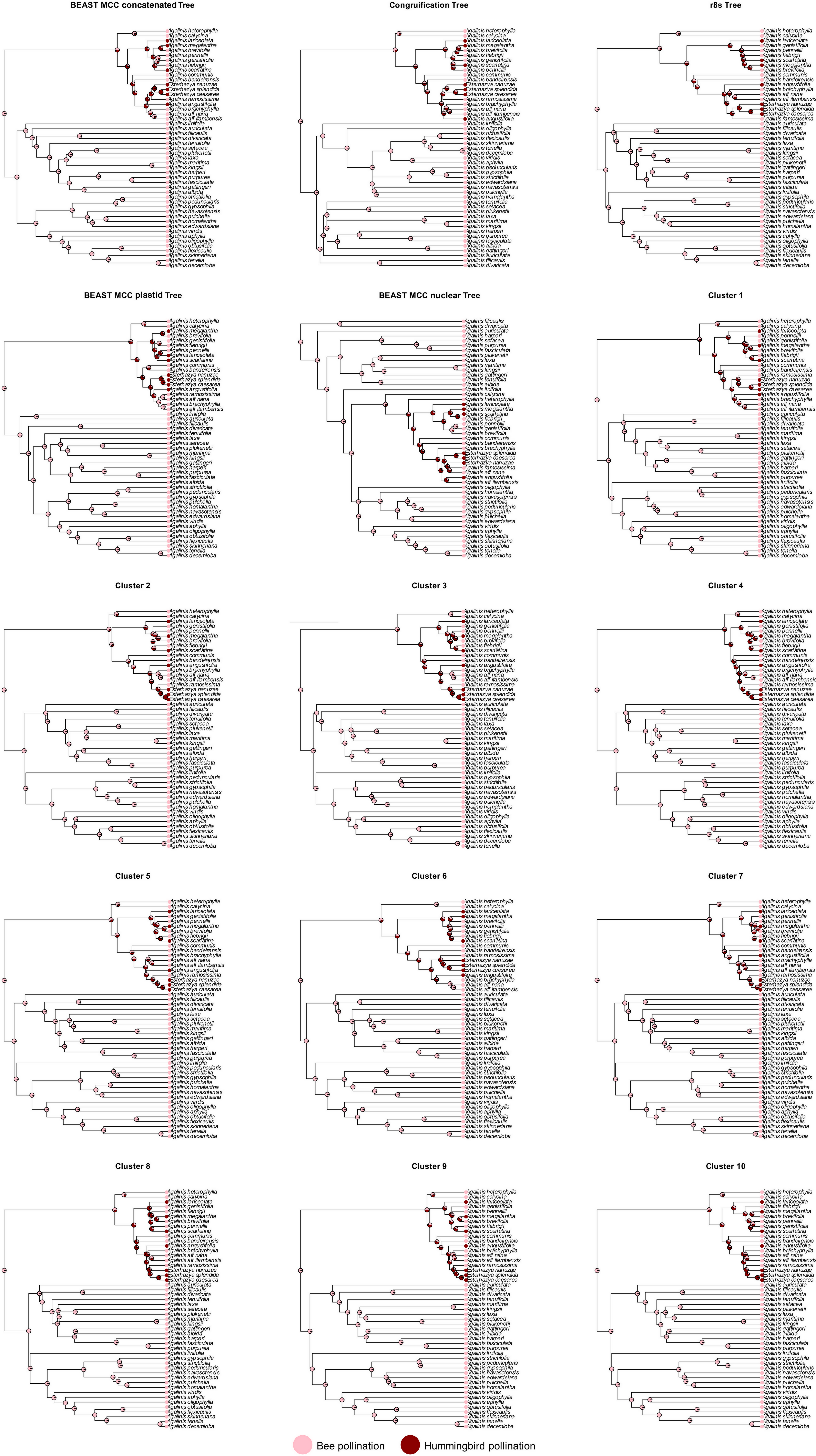

### Figure S5.

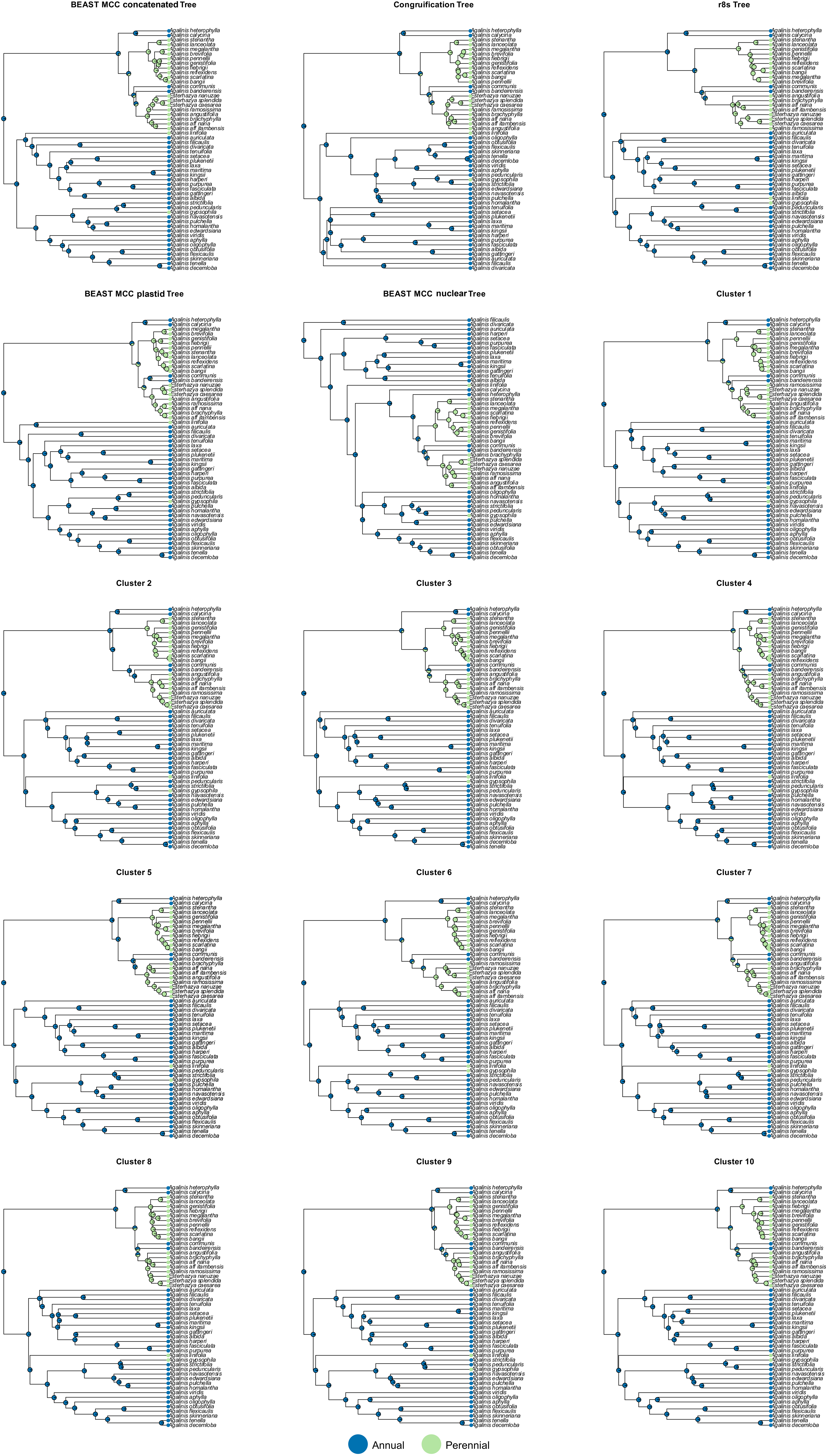
