## Supplementary material for "‘Dispersification’ of *Agalinis* (Orobanchaceae) into South America is associated with hummingbird pollination and perennial life history shifts": Table S1.

**Table S1.** Divergence time estimates for major *Agalinis* clades across different phylogenetic analyses.

|  | **Tree** | **Clade** | **Mean Age (Ma)** | **Lower 95% HPD** | **Upper 95% HPD** |
| --- | --- | --- | --- | --- | --- |
|  | BEAST MCC concatenated | *Agalinis* | 20.14 | 18.18 | 22.10 |
|  |  | North American clade | 18.56 | 16.16 | 20.93 |
|  |  | South American clade + *A. heterophylla* + *A. calycina* | 6.18 | 4.01 | 8.63 |
|  |  | *A. heterophylla* + *A. calycina* | 4.06 | 1.42 | 6.76 |
|  |  | South American clade | 4.92 | 3.31 | 6.68 |
|  |  | Andean clade | 2.51 | 1.49 | 3.66 |
|  |  | Brazilian clade | 4.29 | 2.85 | 5.85 |
| Average of 100 trees | | *Agalinis* | 20.01 | 17.36 | 21.60 |
|  |  | North American clade | 18.39 | 16.17 | 20.36 |
|  |  | South American clade + *A. heterophylla* + *A. calycina* | 6.26 | 3.86 | 8.62 |
|  |  | *A. heterophylla* + *A. calycina* | 4.25 | 2.21 | 6.89 |
|  |  | South American clade | 4.89 | 3.23 | 6.62 |
|  |  | Andean clade | 2.53 | 1.55 | 3.65 |
|  |  | Brazilian clade | 4.26 | 2.66 | 5.65 |

| Congruification | *Agalinis* | 20.25 | NA | NA |
| --- | --- | --- | --- | --- |
|  | North American clade | 18.28 | NA | NA |
|  | South American clade + *A. heterophylla* + *A. calycina* | 8.60 | NA | NA |
|  | *A. heterophylla* + *A. calycina* | 8.13 | NA | NA |
|  | South American clade | 6.08 | NA | NA |
|  | Andean clade | 2.45 | NA | NA |
|  | Brazilian clade | 5.02 | NA | NA |
| r8s | *Agalinis* | 20.63 | NA | NA |
|  | North American clade | 19.54 | NA | NA |
|  | South American clade + *A. heterophylla* + *A. calycina* | 12.78 | NA | NA |
|  | *A. heterophylla* + *A. calycina* | 10.46 | NA | NA |
|  | South American clade | 9.40 | NA | NA |
|  | Andean clade | 3.99 | NA | NA |
|  | Brazilian clade | 8.52 | NA | NA |
| BEAST MCC nuclear | *Agalinis* | 20.14 | 18.16 | 22.09 |
|  | North American clade | NA | NA | NA |
|  | South American clade + *A. heterophylla* + *A. calycina* | 9.69 | 6.61 | 12.91 |
|  | *A. heterophylla* + *A. calycina* | NA | NA | NA |
|  | South American clade | 6.83 | 4.51 | 7.98 |
|  | Andean clade | 4.11 | 2.56 | 5.89 |
|  | Brazilian clade | 5.48 | 3.96 | 7.10 |
| BEAST MCC plastid | *Agalinis* | 20.14 | 18.18 | 22.12 |
|  | North American clade | 18.20 | 15.37 | 20.86 |
|  | South American clade + *A. heterophylla* + *A. calycina* | 4.63 | 2.48 | 7.13 |
|  | *A. heterophylla* + *A. calycina* | 3.03 | 0.85 | 5.41 |
|  | South American clade | 3.87 | 2.15 | 5.84 |
|  | Andean clade | 2.00 | 0.97 | 3.16 |
|  | Brazilian clade | 3.17 | 1.72 | 4.83 |
| BEAST cluster 1 | *Agalinis* | 20.35 | NA | NA |
|  | North American clade | 18.90 | NA | NA |
|  | South American clade + *A. heterophylla* + *A. calycina* | 7.51 | NA | NA |
|  | *A. heterophylla* + *A. calycina* | 5.29 | NA | NA |
|  | South American clade | 5.65 | NA | NA |
|  | Andean clade | 3.92 | NA | NA |
|  | Brazilian clade | 4.92 | NA | NA |
| BEAST cluster 2 | *Agalinis* | 19.15 | NA | NA |
|  | North American clade | 16.82 | NA | NA |
|  | South American clade + *A. heterophylla* + *A. calycina* | 6.94 | NA | NA |
|  | *A. heterophylla* + *A. calycina* | 6.08 | NA | NA |
|  | South American clade | 6.61 | NA | NA |
|  | Andean clade | 2.65 | NA | NA |
|  | Brazilian clade | 4.69 | NA | NA |
| BEAST cluster 3 | *Agalinis* | 19.19 | NA | NA |
|  | North American clade | 17.70 | NA | NA |
|  | South American clade + *A. heterophylla* + *A. calycina* | 7.78 | NA | NA |
|  | *A. heterophylla* + *A. calycina* | 1.48 | NA | NA |
|  | South American clade | 5.01 | NA | NA |
|  | Andean clade | 2.87 | NA | NA |
|  | Brazilian clade | 4.61 | NA | NA |
| BEAST cluster 4 | *Agalinis* | 19.85 | NA | NA |
|  | North American clade | 17.56 | NA | NA |
|  | South American clade + *A. heterophylla* + *A. calycina* | 4.39 | NA | NA |
|  | *A. heterophylla* + *A. calycina* | 2.84 | NA | NA |
|  | South American clade | 3.64 | NA | NA |
|  | Andean clade | 2.41 | NA | NA |
|  | Brazilian clade | 3.28 | NA | NA |
| BEAST cluster 5 | *Agalinis* | 21.56 | NA | NA |
|  | North American clade | 20.25 | NA | NA |
|  | South American clade + *A. heterophylla* + *A. calycina* | 7.61 | NA | NA |
|  | *A. heterophylla* + *A. calycina* | 6.73 | NA | NA |
|  | South American clade | 6.82 | NA | NA |
|  | Andean clade | 2.42 | NA | NA |
|  | Brazilian clade | 4.89 | NA | NA |
| BEAST cluster 6 | *Agalinis* | 19.43 | NA | NA |
|  | North American clade | 18.75 | NA | NA |
|  | South American clade + *A. heterophylla* + *A. calycina* | 9.82 | NA | NA |
|  | *A. heterophylla* + *A. calycina* | 3.21 | NA | NA |
|  | South American clade | 7.86 | NA | NA |
|  | Andean clade | 2.44 | NA | NA |
|  | Brazilian clade | 6.64 | NA | NA |
| BEAST cluster 7 | *Agalinis* | 20.87 | NA | NA |
|  | North American clade | 18.62 | NA | NA |
|  | South American clade + *A. heterophylla* + *A. calycina* | 6.53 | NA | NA |
|  | *A. heterophylla* + *A. calycina* | 5.29 | NA | NA |
|  | South American clade | 4.80 | NA | NA |
|  | Andean clade | 2.45 | NA | NA |
|  | Brazilian clade | 4.59 | NA | NA |
| BEAST cluster 8 | *Agalinis* | 18.99 | NA | NA |
|  | North American clade | 16.12 | NA | NA |
|  | South American clade + *A. heterophylla* + *A. calycina* | 6.07 | NA | NA |
|  | *A. heterophylla* + *A. calycina* | 4.99 | NA | NA |
|  | South American clade | 3.85 | NA | NA |
|  | Andean clade | 2.04 | NA | NA |
|  | Brazilian clade | 3.52 | NA | NA |
| BEAST cluster 9 | *Agalinis* | 18.91 | NA | NA |
|  | North American clade | 17.84 | NA | NA |
|  | South American clade + *A. heterophylla* + *A. calycina* | 4.46 | NA | NA |
|  | *A. heterophylla* + *A. calycina* | 3.79 | NA | NA |
|  | South American clade | 4.15 | NA | NA |
|  | Andean clade | 2.43 | NA | NA |
|  | Brazilian clade | 3.32 | NA | NA |
| BEAST cluster 10 | *Agalinis* | 20.01 | NA | NA |
|  | North American clade | 18.33 | NA | NA |
|  | South American clade + *A. heterophylla* + *A. calycina* | 6.41 | NA | NA |
|  | *A. heterophylla* + *A. calycina* | 4.82 | NA | NA |
|  | South American clade | 4.85 | NA | NA |
|  | Andean clade | 3.43 | NA | NA |
|  | Brazilian clade | 4.06 | NA | NA |

Ma = million years ago; HPD = highest posterior density; NA = not applicable.
