## Supplementary material for "‘Dispersification’ of *Agalinis* (Orobanchaceae) into South America is associated with hummingbird pollination and perennial life history shifts": Table S2.

**Table S2.** Corrected Akaike Information Criterion (AICc) values for biogeographical models tested with BioGeoBEARS across multiple phylogenetic trees of *Agalinis*: the BEAST MCC concatenated tree, congruification tree, r8s tree, BEAST MCC muclear and plastid trees, and Cluster 1–10 trees. The models evaluated include DEC, DIVALIKE, and BAYAREALIKE, also including the variation of each model allowing for ‘jump’ dispersal (+j). For each tree, the best-fitting model is indicated in bold.

| **Tree** | **AICc** | | | | | | | | |
| --- | --- | --- | --- | --- | --- | --- | --- | --- | --- |
|  | **DEC** | **DEC+j** | **DIVALIKE** | | **DIVALIKE+j** | **BAYAREALIKE** | | **BAYAREALIKE+j** | |
| BEAST MCC concatenated | 168.98 | **168.56** | 178.44 | 179.71 | | | 178.44 | | 179.71 |
| Congruification | **163.72** | 165.80 | 172.91 | 173.35 | | | 172.91 | | 173.35 |
| r8s | **167.91** | 168.27 | 182.70 | 182.73 | | | 182.70 | | 182.73 |
| BEAST MCC nuclear | 182.21 | **177.58** | 188.65 | 188.12 | | | 188.65 | | 188.12 |
| BEAST MCC plastid | 167.97 | **167.89** | 178.65 | 180.18 | | | 178.65 | | 180.18 |
| BEAST cluster 1 | 169.72 | **168.88** | 178.78 | 179.52 | | | 178.78 | | 179.52 |
| BEAST cluster 2 | 167.21 | **166.95** | 177.45 | 178.96 | | | 177.45 | | 178.96 |
| BEAST cluster 3 | 167.78 | **167.60** | 181.49 | 182.73 | | | 181.49 | | 182.73 |
| BEAST cluster 4 | 162.48 | **161.72** | 171.17 | 173.01 | | | 171.17 | | 173.01 |
| BEAST cluster 5 | **163.13** | 164.01 | 174.22 | 176.10 | | | 174.22 | | 176.10 |
| BEAST cluster 6 | 171.42 | **171.34** | 180.76 | 182.43 | | | 180.76 | | 182.43 |
| BEAST cluster 7 | **163.44** | 163.91 | 178.34 | 179.66 | | | 178.34 | | 179.66 |
| BEAST cluster 8 | 169.57 | **167.95** | 185.12 | 183.37 | | | 185.12 | | 183.37 |
| BEAST cluster 9 | 170.93 | **168.23** | 185.57 | 181.87 | | | 185.57 | | 181.87 |
| BEAST cluster 10 | 173.39 | **171.31** | 186.74 | 183.72 | | | 186.74 | | 183.72 |
