## Supplementary material for "‘Dispersification’ of *Agalinis* (Orobanchaceae) into South America is associated with hummingbird pollination and perennial life history shifts": Table S4.

**Table S4.** Log-likelihood values (lnL), Akaike Information Criterion (AIC), and corrected AIC (AICc) for State Speciation and Extinction (SSE) models associating life history strategy and diversification in *Agalinis*. Results are presented for the congruification tree, r8s tree, BEAST MCC nuclear and plastid trees, and Cluster 1–10 trees. The Binary SSE (BiSSE) and Hidden SSE (HiSSE) models link diversification rates to life history strategies, while the constant rate, CID2, and CID4 models assume diversification is independent of this trait. In the analyses, 0 denotes annual species, 1 denotes perennial species, and letters A–D indicate hidden states.

| **Tree** | **Model** | **lnL** | **AIC** | **AICc** | **turnover 0A** | **turnover 1A** | **turnover 0B** | **turnover 1B** | **turnover 0C** | **turnover 1C** | **turnover 0D** | **turnover 1D** |
| --- | --- | --- | --- | --- | --- | --- | --- | --- | --- | --- | --- | --- |
| **Congruification** | Constant rate | -172.75 | 353.51 | 354.36 | 0.15 | 0.15 | NA | NA | NA | NA | NA | NA |
|  | **BiSSE** | **-161.51** | **333.02** | **334.32** | **0.11** | **0.49** | **NA** | **NA** | **NA** | **NA** | **NA** | **NA** |
|  | HiSSE | -155.07 | 330.14 | 335.51 | 0.00 | 0.00 | 0.14 | 1.46 | NA | NA | NA | NA |
|  | CID2 | -165.91 | 343.82 | 345.68 | 0.44 | 0.44 | 0.11 | 0.11 | NA | NA | NA | NA |
|  | CID4 | -159.97 | 335.94 | 339.29 | 0.90 | 0.90 | 0.04 | 0.04 | 0.04 | 0.04 | 0.04 | 0.04 |
| **r8s** | Constant rate | -181.48 | 370.96 | 371.81 | 0.12 | 0.12 | NA | NA | NA | NA | NA | NA |
|  | **BiSSE** | **-168.26** | **346.51** | **347.82** | **0.03** | **0.43** | **NA** | **NA** | **NA** | **NA** | **NA** | **NA** |
|  | HiSSE | -165.56 | 351.12 | 356.49 | 0.00 | 0.35 | 0.26 | 0.26 | NA | NA | NA | NA |
|  | CID2 | -173.03 | 358.06 | 359.93 | 0.00 | 0.00 | 0.69 | 0.69 | NA | NA | NA | NA |
|  | CID4 | -168.45 | 352.90 | 356.24 | 0.54 | 0.54 | 0.00 | 0.00 | 0.02 | 0.02 | 0.00 | 0.00 |
| **BEAST MCC nuclear** | Constant rate | -171.31 | 350.62 | 351.48 | 0.15 | 0.15 | NA | NA | NA | NA | NA | NA |
|  | **BiSSE** | **-157.28** | **324.57** | **325.87** | **0.03** | **0.52** | **NA** | **NA** | **NA** | **NA** | **NA** | **NA** |
|  | HiSSE | -155.87 | 331.74 | 337.11 | 0.00 | 0.00 | 0.04 | 0.35 | NA | NA | NA | NA |
|  | CID2 | -166.89 | 345.77 | 347.64 | 0.36 | 0.36 | 0.11 | 0.11 | NA | NA | NA | NA |
|  | CID4 | -158.82 | 333.64 | 336.99 | 0.75 | 0.75 | 0.00 | 0.00 | 0.00 | 0.00 | 0.00 | 0.00 |
| **BEAST MCC plastid** | Constant rate | -170.07 | 348.14 | 348.99 | 0.21 | 0.21 | NA | NA | NA | NA | NA | NA |
|  | **BiSSE** | **-151.58** | **313.16** | **314.46** | **0.10** | **1.64** | **NA** | **NA** | **NA** | **NA** | **NA** | **NA** |
|  | HiSSE | -148.41 | 316.83 | 322.19 | 0.05 | 0.26 | 0.13 | 0.81 | NA | NA | NA | NA |
|  | CID2 | -158.44 | 328.88 | 330.75 | 0.06 | 0.06 | 2.34 | 2.34 | NA | NA | NA | NA |
|  | CID4 | -155.91 | 327.83 | 331.18 | 0.00 | 0.00 | 0.64 | 0.64 | 0.00 | 0.00 | 0.11 | 0.11 |
| **BEAST cluster 1** | Constant rate | -177.03 | 362.07 | 362.92 | 0.13 | 0.13 | NA | NA | NA | NA | NA | NA |
|  | **BiSSE** | **-160.63** | **331.26** | **332.56** | **0.05** | **0.73** | **NA** | **NA** | **NA** | **NA** | **NA** | **NA** |
|  | HiSSE | -158.97 | 337.95 | 343.31 | 0.00 | 0.00 | 0.07 | 0.41 | NA | NA | NA | NA |
|  | CID2 | -168.06 | 348.12 | 349.98 | 0.00 | 0.00 | 1.01 | 1.01 | NA | NA | NA | NA |
|  | CID4 | -163.76 | 343.53 | 346.88 | 0.04 | 0.04 | 0.04 | 0.04 | 1.11 | 1.11 | 0.04 | 0.04 |
| **BEAST cluster 2** | Constant rate | -164.92 | 337.84 | 338.70 | 0.17 | 0.17 | NA | NA | NA | NA | NA | NA |
|  | **BiSSE** | **-152.45** | **314.91** | **316.21** | **0.07** | **1.17** | **NA** | **NA** | **NA** | **NA** | **NA** | **NA** |
|  | HiSSE | -151.68 | 323.37 | 328.73 | 0.18 | 0.70 | 0.00 | 0.00 | NA | NA | NA | NA |
|  | CID2 | -157.63 | 327.27 | 329.13 | 0.00 | 0.00 | 1.11 | 1.11 | NA | NA | NA | NA |
|  | CID4 | -155.79 | 327.58 | 330.93 | 0.03 | 0.03 | 0.03 | 0.03 | 0.03 | 0.03 | 0.95 | 0.95 |
| **BEAST cluster 3** | Constant rate | -171.99 | 351.99 | 352.84 | 0.14 | 0.14 | NA | NA | NA | NA | NA | NA |
|  | **BiSSE** | **-153.98** | **317.95** | **319.25** | **0.07** | **1.30** | **NA** | **NA** | **NA** | **NA** | **NA** | **NA** |
|  | HiSSE | -151.67 | 323.33 | 328.70 | 0.10 | 0.00 | 0.07 | 1.44 | NA | NA | NA | NA |
|  | CID2 | -160.34 | 332.68 | 334.54 | 2.63 | 2.63 | 0.05 | 0.05 | NA | NA | NA | NA |
|  | CID4 | -156.82 | 329.63 | 332.98 | 1.92 | 1.92 | 0.07 | 0.07 | 0.07 | 0.07 | 0.07 | 0.07 |
| **BEAST cluster 4** | Constant rate | -168.77 | 345.54 | 346.39 | 0.16 | 0.16 | NA | NA | NA | NA | NA | NA |
|  | **BiSSE** | **-149.43** | **308.87** | **310.17** | **0.04** | **1.10** | **NA** | **NA** | **NA** | **NA** | **NA** | **NA** |
|  | HiSSE | -146.38 | 312.76 | 318.12 | 0.02 | 0.32 | 0.00 | 0.65 | NA | NA | NA | NA |
|  | CID2 | -157.97 | 327.95 | 329.81 | 0.11 | 0.11 | 0.55 | 0.55 | NA | NA | NA | NA |
|  | CID4 | -151.35 | 318.70 | 318.70 | 0.02 | 0.02 | 0.02 | 0.02 | 0.02 | 0.02 | 1.63 | 1.63 |
| **BEAST cluster 5** | Constant rate | -175.84 | 359.67 | 360.52 | 0.13 | 0.13 | NA | NA | NA | NA | NA | NA |
|  | **BiSSE** | **-160.83** | **331.67** | **332.97** | **0.06** | **0.95** | **NA** | **NA** | **NA** | **NA** | **NA** | **NA** |
|  | HiSSE | -159.17 | 338.35 | 343.71 | 0.13 | 0.00 | 0.00 | 0.52 | NA | NA | NA | NA |
|  | CID2 | -167.66 | 347.33 | 349.19 | 1.67 | 1.67 | 0.02 | 0.02 | NA | NA | NA | NA |
|  | CID4 | -164.31 | 344.62 | 347.97 | 0.05 | 0.05 | 0.05 | 0.05 | 1.27 | 1.27 | 0.05 | 0.05 |
| **BEAST cluster 6** | Constant rate | -171.90 | 351.80 | 352.65 | 0.14 | 0.14 | NA | NA | NA | NA | NA | NA |
|  | **BiSSE** | **-162.51** | **335.02** | **336.33** | **0.07** | **0.70** | **NA** | **NA** | **NA** | **NA** | **NA** | **NA** |
|  | HiSSE | -159.83 | 339.66 | 345.03 | 0.15 | 0.00 | 0.00 | 0.47 | NA | NA | NA | NA |
|  | CID2 | -165.82 | 343.64 | 345.51 | 0.00 | 0.00 | 0.45 | 0.45 | NA | NA | NA | NA |
|  | CID4 | -163.41 | 342.83 | 346.18 | 0.03 | 0.03 | 0.53 | 0.53 | 0.03 | 0.03 | 0.00 | 0.00 |
| **BEAST cluster 7** | Constant rate | -171.60 | 351.19 | 352.04 | 0.18 | 0.18 | NA | NA | NA | NA | NA | NA |
|  | **BiSSE** | **-151.02** | **312.04** | **313.34** | **0.06** | **1.64** | **NA** | **NA** | **NA** | **NA** | **NA** | **NA** |
|  | HiSSE | -148.52 | 317.04 | 322.41 | 0.12 | 0.03 | 0.00 | 2.14 | NA | NA | NA | NA |
|  | CID2 | -156.66 | 325.33 | 327.19 | 4.23 | 4.23 | 0.04 | 0.04 | NA | NA | NA | NA |
|  | CID4 | -154.12 | 324.23 | 327.58 | 0.01 | 0.01 | 2.34 | 2.34 | 0.07 | 0.07 | 0.10 | 0.10 |
| **BEAST cluster 8** | Constant rate | -167.28 | 342.55 | 343.40 | 0.16 | 0.16 | NA | NA | NA | NA | NA | NA |
|  | **BiSSE** | **-152.52** | **315.04** | **316.35** | **0.05** | **0.72** | **NA** | **NA** | **NA** | **NA** | **NA** | **NA** |
|  | HiSSE | -150.45 | 320.90 | 326.27 | 0.00 | 0.00 | 0.17 | 0.59 | NA | NA | NA | NA |
|  | CID2 | -156.85 | 325.70 | 327.57 | 1.11 | 1.11 | 0.00 | 0.00 | NA | NA | NA | NA |
|  | CID4 | -152.57 | 321.14 | 324.49 | 0.02 | 0.02 | 1.03 | 1.03 | 0.00 | 0.00 | 0.02 | 0.02 |
| **BEAST cluster 9** | Constant rate | -169.91 | 347.82 | 348.67 | 0.15 | 0.15 | NA | NA | NA | NA | NA | NA |
|  | **BiSSE** | **-150.21** | **310.41** | **311.72** | **0.05** | **1.16** | **NA** | **NA** | **NA** | **NA** | **NA** | **NA** |
|  | HiSSE | -147.44 | 314.88 | 320.24 | 0.04 | 1.52 | 0.08 | 0.00 | NA | NA | NA | NA |
|  | CID2 | -157.47 | 326.94 | 328.81 | 0.00 | 0.00 | 1.87 | 1.87 | NA | NA | NA | NA |
|  | CID4 | -153.21 | 322.42 | 325.77 | 1.80 | 1.80 | 0.04 | 0.04 | 0.04 | 0.04 | 0.04 | 0.04 |
| **BEAST cluster 10** | Constant rate | -172.32 | 352.64 | 353.49 | 0.14 | 0.14 | NA | NA | NA | NA | NA | NA |
|  | **BiSSE** | **-154.85** | **319.70** | **321.00** | **0.05** | **0.90** | **NA** | **NA** | **NA** | **NA** | **NA** | **NA** |
|  | HiSSE | -151.96 | 323.91 | 329.28 | 0.00 | 0.55 | 0.00 | 0.00 | NA | NA | NA | NA |
|  | CID2 | -161.64 | 335.27 | 337.14 | 0.00 | 0.00 | 1.32 | 1.32 | NA | NA | NA | NA |
|  | CID4 | -157.45 | 330.91 | 334.26 | 0.03 | 0.03 | 1.17 | 1.17 | 0.03 | 0.03 | 0.03 | 0.03 |
