## Supplementary material for "‘Dispersification’ of *Agalinis* (Orobanchaceae) into South America is associated with hummingbird pollination and perennial life history shifts": Table S5.

**Table S5.** Log-likelihood values (lnL), Akaike Information Criterion (AIC), and corrected AIC (AICc) for State Speciation and Extinction (SSE) models associating life history strategy and diversification in *Agalinis* for the 100 randomly sampled post–burn-in MCMC trees. The Binary SSE (BiSSE) and Hidden State SSE (HiSSE) models link diversification rates to life history strategy, while the Constant rate rate, CID2, and CID4 models assume diversification is independent of this trait.

| File | Model | lnL | AIC | AICc |
|---|---|---|---|---|
| Tree1 | Constant rate | -175.44 | 358.88 | 359.73 |
| | **BiSSE** | **-157.10** | **324.20** | **325.50** |
| | HiSSE | -155.99 | 331.97 | 337.34 |
| | CID2 | -163.33 | 338.66 | 340.53 |
| | CID4 | -164.52 | 345.03 | 348.38 |
| Tree2 | Constant rate | -170.49 | 348.98 | 349.83 |
| | **BiSSE** | **-152.02** | **314.03** | **315.33** |
| | HiSSE | -149.58 | 319.15 | 324.52 |
| | CID2 | -162.59 | 337.19 | 339.05 |
| | CID4 | -153.39 | 322.79 | 326.14 |
| Tree3 | Constant rate | -173.09 | 354.18 | 355.03 |
| | **BiSSE** | **-152.71** | **315.42** | **316.73** |
| | HiSSE | -151.67 | 323.35 | 328.71 |
| | CID2 | -159.70 | 331.41 | 333.28 |
| | CID4 | -160.89 | 337.78 | 341.13 |
| Tree4 | Constant rate | -174.99 | 357.98 | 358.83 |
| | **BiSSE** | **-158.61** | **327.22** | **328.53** |
| | HiSSE | -157.13 | 334.25 | 339.62 |
| | CID2 | -164.56 | 341.12 | 342.98 |
| | CID4 | -160.85 | 337.71 | 341.06 |
| Tree5 | Constant rate | -163.24 | 334.48 | 335.33 |
| | **BiSSE** | **-148.02** | **306.04** | **307.35** |
| | HiSSE | -146.29 | 312.59 | 317.95 |
| | CID2 | -152.41 | 316.82 | 318.68 |
| | CID4 | -151.75 | 319.50 | 322.85 |
| Tree6 | Constant rate | -181.53 | 371.06 | 371.91 |
| | **BiSSE** | **-164.04** | **338.07** | **339.38** |
| | HiSSE | -163.15 | 346.30 | 351.66 |
| | CID2 | -171.18 | 354.35 | 356.22 |
| | CID4 | -166.82 | 349.65 | 352.99 |
| Tree7 | Constant rate | -173.82 | 355.65 | 356.50 |
| | **BiSSE** | **-151.95** | **313.90** | **315.20** |
| | HiSSE | -148.96 | 317.92 | 323.28 |
| | CID2 | -158.53 | 329.05 | 330.92 |
| | CID4 | -154.47 | 324.94 | 328.29 |
| Tree8 | Constant rate | -179.47 | 366.94 | 367.80 |
| | **BiSSE** | **-163.11** | **336.21** | **337.52** |
| | HiSSE | -160.12 | 340.25 | 345.61 |
| | CID2 | -170.89 | 353.78 | 355.64 |

| | | | | |
|---|---|---|---|---|
| | CID4 | -165.23 | 346.46 | 349.81 |
| Tree9 | Constant rate | -168.95 | 345.90 | 346.75 |
| | **BiSSE** | **-151.49** | **312.98** | **314.29** |
| | HiSSE | -149.95 | 319.90 | 325.26 |
| | CID2 | -159.08 | 330.17 | 332.03 |
| | CID4 | -155.25 | 326.50 | 329.85 |
| Tree10 | Constant rate | -177.09 | 362.19 | 363.04 |
| | **BiSSE** | **-157.56** | **325.12** | **326.42** |
| | HiSSE | -156.32 | 332.65 | 338.01 |
| | CID2 | -164.38 | 340.75 | 342.62 |
| | CID4 | -162.13 | 340.26 | 343.61 |
| Tree11 | Constant rate | -174.18 | 356.37 | 357.22 |
| | **BiSSE** | **-154.79** | **319.59** | **320.89** |
| | HiSSE | -153.69 | 327.38 | 332.75 |
| | CID2 | -163.38 | 338.77 | 340.63 |
| | CID4 | -156.94 | 329.89 | 333.24 |
| Tree12 | Constant rate | -176.44 | 360.88 | 361.73 |
| | **BiSSE** | **-157.21** | **324.43** | **325.73** |
| | HiSSE | -154.27 | 328.54 | 333.91 |
| | CID2 | -164.33 | 340.65 | 342.52 |
| | CID4 | -160.11 | 336.21 | 339.56 |
| Tree13 | Constant rate | -175.39 | 358.78 | 359.63 |
| | **BiSSE** | **-158.33** | **326.65** | **327.96** |
| | HiSSE | -155.71 | 331.42 | 336.78 |
| | CID2 | -165.81 | 343.62 | 345.48 |
| | CID4 | -161.92 | 339.84 | 343.19 |
| Tree14 | Constant rate | -173.27 | 354.54 | 355.39 |
| | **BiSSE** | **-156.89** | **323.79** | **325.09** |
| | HiSSE | -155.12 | 330.25 | 335.62 |
| | CID2 | -162.06 | 336.12 | 337.98 |
| | CID4 | -158.80 | 333.61 | 336.95 |
| Tree15 | Constant rate | -174.52 | 357.04 | 357.89 |
| | **BiSSE** | **-153.11** | **316.22** | **317.53** |
| | HiSSE | -153.60 | 327.19 | 332.56 |
| | CID2 | -159.05 | 330.09 | 331.96 |
| | CID4 | -155.06 | 326.13 | 329.47 |
| Tree16 | Constant rate | -174.99 | 357.99 | 358.84 |
| | **BiSSE** | **-157.01** | **324.03** | **325.33** |
| | HiSSE | -154.08 | 328.16 | 333.53 |
| | CID2 | -166.12 | 344.25 | 346.11 |
| | CID4 | -159.49 | 334.99 | 338.33 |
| Tree17 | Constant rate | -170.55 | 349.10 | 349.95 |
| | **BiSSE** | **-155.03** | **320.06** | **321.36** |
| | HiSSE | -152.91 | 325.81 | 331.18 |
| | CID2 | -161.22 | 334.43 | 336.30 |
| | CID4 | -157.50 | 331.01 | 334.36 |

| | | | | |
|---|---|---|---|---|
| Tree18 | Constant rate | -167.18 | 342.36 | 343.21 |
| | **BiSSE** | **-152.51** | **315.02** | **316.33** |
| | HiSSE | -149.10 | 318.20 | 323.57 |
| | CID2 | -159.41 | 330.82 | 332.69 |
| | CID4 | -154.19 | 324.38 | 327.73 |
| Tree19 | Constant rate | -174.15 | 356.31 | 357.16 |
| | **BiSSE** | **-153.24** | **316.48** | **317.78** |
| | HiSSE | -150.90 | 321.80 | 327.17 |
| | CID2 | -163.50 | 338.99 | 340.86 |
| | CID4 | -157.38 | 330.77 | 334.12 |
| Tree20 | Constant rate | -172.72 | 353.44 | 354.29 |
| | BiSSE | -158.67 | 327.33 | 328.63 |
| | **HiSSE** | **-156.04** | **332.08** | **337.45** |
| | CID2 | -164.76 | 341.51 | 343.38 |
| | CID4 | -161.15 | 338.30 | 341.65 |
| Tree21 | Constant rate | -171.94 | 351.87 | 352.72 |
| | **BiSSE** | **-154.95** | **319.91** | **321.21** |
| | HiSSE | -152.06 | 324.11 | 329.48 |
| | CID2 | -160.36 | 332.72 | 334.58 |
| | CID4 | -157.74 | 331.48 | 334.83 |
| Tree22 | Constant rate | -174.59 | 357.18 | 358.03 |
| | **BiSSE** | **-151.72** | **313.44** | **314.74** |
| | HiSSE | -151.72 | 323.44 | 328.80 |
| | CID2 | -157.45 | 326.89 | 328.76 |
| | CID4 | -154.49 | 324.97 | 328.32 |
| Tree23 | Constant rate | -174.85 | 357.70 | 358.55 |
| | **BiSSE** | **-161.97** | **333.94** | **335.24** |
| | HiSSE | -153.71 | 327.43 | 332.79 |
| | CID2 | -165.13 | 342.27 | 344.13 |
| | CID4 | -158.78 | 333.55 | 336.90 |
| Tree24 | Constant rate | -173.84 | 355.68 | 356.53 |
| | **BiSSE** | **-159.90** | **329.81** | **331.11** |
| | HiSSE | -157.56 | 335.13 | 340.49 |
| | CID2 | -166.68 | 345.36 | 347.22 |
| | CID4 | -163.84 | 343.67 | 347.02 |
| Tree25 | Constant rate | -170.55 | 349.11 | 349.96 |
| | **BiSSE** | **-149.51** | **309.02** | **310.32** |
| | HiSSE | -147.61 | 315.22 | 320.59 |
| | CID2 | -157.35 | 326.70 | 328.56 |
| | CID4 | -156.66 | 329.32 | 332.67 |
| Tree26 | Constant rate | -171.63 | 351.26 | 352.11 |
| | **BiSSE** | **-155.13** | **320.26** | **321.56** |
| | HiSSE | -152.23 | 324.45 | 329.82 |
| | CID2 | -163.33 | 338.65 | 340.52 |
| | CID4 | -158.02 | 332.03 | 335.38 |
| | Constant rate | -172.03 | 352.05 | 352.91 |

| | | | | |
|---|---|---|---|---|
| Tree27 | **BiSSE** | **-155.04** | **320.08** | **321.38** |
| | HiSSE | -154.11 | 328.23 | 333.59 |
| | CID2 | -161.26 | 334.53 | 336.39 |
| | CID4 | -157.96 | 331.92 | 335.27 |
| Tree28 | Constant rate | -171.50 | 351.01 | 351.86 |
| | **BiSSE** | **-152.06** | **314.12** | **315.43** |
| | HiSSE | -152.13 | 324.26 | 329.63 |
| | CID2 | -162.32 | 336.64 | 338.51 |
| | CID4 | -155.31 | 326.62 | 329.97 |
| Tree29 | Constant rate | -173.54 | 355.08 | 355.93 |
| | **BiSSE** | **-161.18** | **332.36** | **333.66** |
| | HiSSE | -159.06 | 338.13 | 343.50 |
| | CID2 | -169.02 | 350.03 | 351.90 |
| | CID4 | -162.52 | 341.05 | 344.40 |
| Tree30 | Constant rate | -178.70 | 365.40 | 366.25 |
| | **BiSSE** | **-161.25** | **332.51** | **333.81** |
| | HiSSE | -158.81 | 337.62 | 342.98 |
| | CID2 | -168.05 | 348.09 | 349.96 |
| | CID4 | -164.97 | 345.95 | 349.30 |
| Tree31 | Constant rate | -183.06 | 374.12 | 374.97 |
| | **BiSSE** | **-163.56** | **337.13** | **338.43** |
| | HiSSE | -163.30 | 346.61 | 351.97 |
| | CID2 | -174.17 | 360.34 | 362.21 |
| | CID4 | -166.08 | 348.17 | 351.51 |
| Tree32 | Constant rate | -177.12 | 362.23 | 363.08 |
| | **BiSSE** | **-160.23** | **330.46** | **331.77** |
| | HiSSE | -157.66 | 335.32 | 340.69 |
| | CID2 | -165.92 | 343.84 | 345.71 |
| | CID4 | -162.63 | 341.25 | 344.60 |
| Tree33 | Constant rate | -177.68 | 363.36 | 364.21 |
| | **BiSSE** | **-156.18** | **322.35** | **323.66** |
| | HiSSE | -153.69 | 327.37 | 332.74 |
| | CID2 | -163.46 | 338.93 | 340.79 |
| | CID4 | -159.52 | 335.04 | 338.38 |
| Tree34 | Constant rate | -168.99 | 345.98 | 346.83 |
| | **BiSSE** | **-147.31** | **304.62** | **305.93** |
| | HiSSE | -145.34 | 310.68 | 316.05 |
| | CID2 | -153.81 | 319.63 | 321.49 |
| | CID4 | -155.12 | 326.25 | 329.60 |
| Tree35 | Constant rate | -173.49 | 354.98 | 355.83 |
| | **BiSSE** | **-153.02** | **316.03** | **317.34** |
| | HiSSE | -152.62 | 325.23 | 330.60 |
| | CID2 | -161.02 | 334.04 | 335.91 |
| | CID4 | -157.86 | 331.72 | 335.07 |
| | Constant rate | -172.04 | 352.09 | 352.94 |
| | **BiSSE** | **-155.34** | **320.68** | **321.99** |

| | | | | |
|---|---|---|---|---|
| Tree36 | HiSSE | -152.32 | 324.64 | 330.00 |
| | CID2 | -162.41 | 336.83 | 338.69 |
| | CID4 | -158.36 | 332.72 | 336.07 |
| Tree37 | Constant rate | -175.77 | 359.54 | 360.39 |
| | **BiSSE** | **-155.71** | **321.42** | **322.72** |
| | HiSSE | -152.85 | 325.70 | 331.07 |
| | CID2 | -166.79 | 345.57 | 347.44 |
| | CID4 | -159.24 | 334.47 | 337.82 |
| Tree38 | Constant rate | -171.87 | 351.73 | 352.58 |
| | **BiSSE** | **-154.25** | **318.51** | **319.81** |
| | HiSSE | -152.85 | 325.70 | 331.07 |
| | CID2 | -158.98 | 329.95 | 331.82 |
| | CID4 | -156.18 | 328.36 | 331.71 |
| Tree39 | Constant rate | -174.07 | 356.13 | 356.99 |
| | **BiSSE** | **-157.75** | **325.51** | **326.81** |
| | HiSSE | -156.49 | 332.97 | 338.34 |
| | CID2 | -163.71 | 339.42 | 341.29 |
| | CID4 | -160.07 | 336.13 | 339.48 |
| Tree40 | Constant rate | -173.29 | 354.58 | 355.43 |
| | **BiSSE** | **-153.17** | **316.34** | **317.65** |
| | HiSSE | -151.08 | 322.15 | 327.52 |
| | CID2 | -163.02 | 338.04 | 339.90 |
| | CID4 | -156.86 | 329.73 | 333.07 |
| Tree41 | Constant rate | -171.45 | 350.89 | 351.74 |
| | **BiSSE** | **-153.05** | **316.10** | **317.41** |
| | HiSSE | -153.05 | 326.10 | 331.47 |
| | CID2 | -161.43 | 334.85 | 336.72 |
| | CID4 | -154.94 | 325.88 | 329.23 |
| Tree42 | Constant rate | -168.80 | 345.60 | 346.45 |
| | **BiSSE** | **-155.26** | **320.52** | **321.83** |
| | HiSSE | -149.82 | 319.64 | 325.00 |
| | CID2 | -156.90 | 325.80 | 327.67 |
| | CID4 | -154.90 | 325.79 | 329.14 |
| Tree43 | Constant rate | -175.39 | 358.77 | 359.63 |
| | **BiSSE** | **-160.82** | **331.63** | **332.94** |
| | HiSSE | -155.94 | 331.89 | 337.25 |
| | CID2 | -163.31 | 338.62 | 340.49 |
| | CID4 | -159.67 | 335.34 | 338.69 |
| Tree44 | Constant rate | -169.54 | 347.09 | 347.94 |
| | **BiSSE** | **-153.09** | **316.18** | **317.49** |
| | HiSSE | -150.82 | 321.64 | 327.00 |
| | CID2 | -161.84 | 335.68 | 337.54 |
| | CID4 | -156.29 | 328.58 | 331.93 |
| Tree45 | Constant rate | -172.27 | 352.55 | 353.40 |
| | **BiSSE** | **-160.84** | **331.68** | **332.98** |
| | HiSSE | -158.42 | 336.83 | 342.20 |

| | | | | |
|---|---|---|---|---|
| | CID2 | -165.22 | 342.44 | 344.31 |
| | CID4 | -162.25 | 340.51 | 343.86 |
| Tree46 | Constant rate | -170.46 | 348.92 | 349.77 |
| | **BiSSE** | **-153.02** | **316.03** | **317.34** |
| | HiSSE | -151.68 | 323.37 | 328.73 |
| | CID2 | -158.38 | 328.75 | 330.62 |
| | CID4 | -155.85 | 327.70 | 331.05 |
| Tree47 | Constant rate | -172.17 | 352.34 | 353.19 |
| | **BiSSE** | **-153.72** | **317.44** | **318.74** |
| | HiSSE | -151.33 | 322.66 | 328.03 |
| | CID2 | -160.56 | 333.12 | 334.98 |
| | CID4 | -156.74 | 329.48 | 332.83 |
| Tree48 | Constant rate | -170.95 | 349.90 | 350.75 |
| | **BiSSE** | **-153.42** | **316.83** | **318.14** |
| | HiSSE | -151.90 | 323.79 | 329.16 |
| | CID2 | -160.40 | 332.80 | 334.67 |
| | CID4 | -157.44 | 330.88 | 334.23 |
| Tree49 | Constant rate | -178.29 | 364.57 | 365.43 |
| | **BiSSE** | **-166.76** | **343.52** | **344.83** |
| | HiSSE | -161.23 | 342.46 | 347.83 |
| | CID2 | -169.86 | 351.73 | 353.59 |
| | CID4 | -163.84 | 343.69 | 347.04 |
| Tree50 | Constant rate | -171.61 | 351.23 | 352.08 |
| | **BiSSE** | **-155.37** | **320.74** | **322.05** |
| | HiSSE | -154.46 | 328.92 | 334.29 |
| | CID2 | -162.05 | 336.10 | 337.97 |
| | CID4 | -158.71 | 333.41 | 336.76 |
| Tree51 | Constant rate | -174.93 | 357.85 | 358.70 |
| | **BiSSE** | **-158.96** | **327.91** | **329.22** |
| | HiSSE | -158.29 | 336.58 | 341.95 |
| | CID2 | -166.67 | 345.33 | 347.20 |
| | CID4 | -162.66 | 341.32 | 344.67 |
| Tree52 | Constant rate | -166.39 | 340.79 | 341.64 |
| | **BiSSE** | **-156.25** | **322.50** | **323.80** |
| | HiSSE | -152.81 | 325.62 | 330.99 |
| | CID2 | -159.65 | 331.31 | 333.17 |
| | CID4 | -158.39 | 332.78 | 336.13 |
| Tree53 | Constant rate | -167.38 | 342.76 | 343.61 |
| | **BiSSE** | **-153.11** | **316.22** | **317.52** |
| | HiSSE | -147.26 | 314.52 | 319.89 |
| | CID2 | -154.95 | 321.91 | 323.78 |
| | CID4 | -155.56 | 327.11 | 330.46 |
| Tree54 | Constant rate | -169.57 | 347.15 | 348.00 |
| | **BiSSE** | **-153.69** | **317.38** | **318.68** |
| | HiSSE | -151.34 | 322.68 | 328.04 |
| | CID2 | -159.58 | 331.17 | 333.03 |

| | | | | |
|---|---|---|---|---|
| | CID4 | -158.27 | 332.54 | 335.89 |
| Tree55 | Constant rate | -174.36 | 356.73 | 357.58 |
| | **BiSSE** | **-155.90** | **321.81** | **323.11** |
| | HiSSE | -154.02 | 328.04 | 333.41 |
| | CID2 | -164.10 | 340.20 | 342.06 |
| | CID4 | -160.62 | 337.23 | 340.58 |
| Tree56 | Constant rate | -175.36 | 358.72 | 359.57 |
| | **BiSSE** | **-152.00** | **314.01** | **315.31** |
| | HiSSE | -149.54 | 319.09 | 324.45 |
| | CID2 | -164.48 | 340.95 | 342.82 |
| | CID4 | -155.94 | 327.89 | 331.24 |
| Tree57 | Constant rate | -168.26 | 344.51 | 345.37 |
| | **BiSSE** | **-151.52** | **313.04** | **314.34** |
| | HiSSE | -151.30 | 322.60 | 327.96 |
| | CID2 | -158.84 | 329.68 | 331.54 |
| | CID4 | -155.62 | 327.24 | 330.59 |
| Tree58 | Constant rate | -174.69 | 357.37 | 358.22 |
| | **BiSSE** | **-153.00** | **316.00** | **317.31** |
| | HiSSE | -152.18 | 324.36 | 329.73 |
| | CID2 | -161.64 | 335.27 | 337.14 |
| | CID4 | -158.01 | 332.02 | 335.37 |
| Tree59 | Constant rate | -175.10 | 358.19 | 359.04 |
| | **BiSSE** | **-152.48** | **314.96** | **316.26** |
| | HiSSE | -149.97 | 319.93 | 325.30 |
| | CID2 | -160.22 | 332.43 | 334.30 |
| | CID4 | -156.45 | 328.89 | 332.24 |
| Tree60 | Constant rate | -177.72 | 363.44 | 364.29 |
| | **BiSSE** | **-166.16** | **342.32** | **343.62** |
| | HiSSE | -160.84 | 341.68 | 347.04 |
| | CID2 | -168.16 | 348.31 | 350.18 |
| | CID4 | -164.61 | 345.21 | 348.56 |
| Tree61 | Constant rate | -171.50 | 351.00 | 351.85 |
| | **BiSSE** | **-154.61** | **319.22** | **320.53** |
| | HiSSE | -152.18 | 324.37 | 329.73 |
| | CID2 | -161.96 | 335.93 | 337.80 |
| | CID4 | -157.98 | 331.97 | 335.31 |
| Tree62 | Constant rate | -173.65 | 355.31 | 356.16 |
| | **BiSSE** | **-157.92** | **325.85** | **327.15** |
| | HiSSE | -157.15 | 334.30 | 339.67 |
| | CID2 | -164.94 | 341.88 | 343.74 |
| | CID4 | -161.13 | 338.26 | 341.61 |
| Tree63 | Constant rate | -165.99 | 339.99 | 340.84 |
| | **BiSSE** | **-150.72** | **311.43** | **312.74** |
| | HiSSE | -147.64 | 315.27 | 320.64 |
| | CID2 | -155.44 | 322.89 | 324.75 |
| | CID4 | -154.46 | 324.92 | 328.27 |

| | | | | |
|---|---|---|---|---|
| Tree64 | Constant rate | -173.42 | 354.83 | 355.68 |
| | **BiSSE** | **-160.13** | **330.26** | **331.57** |
| | HiSSE | -156.81 | 333.62 | 338.98 |
| | CID2 | -164.52 | 341.04 | 342.91 |
| | CID4 | -161.50 | 339.00 | 342.35 |
| Tree65 | Constant rate | -169.72 | 347.43 | 348.28 |
| | **BiSSE** | **-156.42** | **322.85** | **324.15** |
| | HiSSE | -154.18 | 328.36 | 333.72 |
| | CID2 | -161.77 | 335.54 | 337.41 |
| | CID4 | -159.04 | 334.08 | 337.43 |
| Tree66 | Constant rate | -168.04 | 344.08 | 344.93 |
| | **BiSSE** | **-155.02** | **320.03** | **321.34** |
| | HiSSE | -150.18 | 320.35 | 325.72 |
| | CID2 | -158.80 | 329.60 | 331.47 |
| | CID4 | -157.77 | 331.54 | 334.89 |
| Tree67 | Constant rate | -175.52 | 359.04 | 359.89 |
| | **BiSSE** | **-154.37** | **318.74** | **320.04** |
| | HiSSE | -151.89 | 323.78 | 329.15 |
| | CID2 | -161.25 | 334.51 | 336.37 |
| | CID4 | -157.30 | 330.60 | 333.95 |
| Tree68 | Constant rate | -166.36 | 340.73 | 341.58 |
| | **BiSSE** | **-153.83** | **317.67** | **318.97** |
| | HiSSE | -149.45 | 318.91 | 324.28 |
| | CID2 | -156.36 | 324.71 | 326.58 |
| | CID4 | -152.97 | 321.94 | 325.29 |
| Tree69 | Constant rate | -177.00 | 362.00 | 362.85 |
| | **BiSSE** | **-157.83** | **325.66** | **326.97** |
| | HiSSE | -156.15 | 332.30 | 337.66 |
| | CID2 | -165.99 | 343.99 | 345.85 |
| | CID4 | -160.98 | 337.96 | 341.31 |
| Tree70 | Constant rate | -169.33 | 346.66 | 347.51 |
| | **BiSSE** | **-150.89** | **311.78** | **313.09** |
| | HiSSE | -150.44 | 320.89 | 326.25 |
| | CID2 | -158.95 | 329.90 | 331.77 |
| | CID4 | -155.32 | 326.63 | 329.98 |
| Tree71 | Constant rate | -174.80 | 357.60 | 358.45 |
| | **BiSSE** | **-154.36** | **318.72** | **320.02** |
| | HiSSE | -152.34 | 324.69 | 330.05 |
| | CID2 | -162.48 | 336.96 | 338.82 |
| | CID4 | -158.67 | 333.35 | 336.70 |
| Tree72 | Constant rate | -166.24 | 340.47 | 341.32 |
| | **BiSSE** | **-147.20** | **304.40** | **305.70** |
| | HiSSE | -145.30 | 310.60 | 315.97 |
| | CID2 | -153.10 | 318.20 | 320.07 |
| | CID4 | -150.01 | 316.01 | 319.36 |
| | Constant rate | -171.60 | 351.19 | 352.04 |

| | | | | |
|---|---|---|---|---|
| Tree73 | **BiSSE** | **-156.24** | **322.47** | **323.78** |
| | HiSSE | -154.18 | 328.36 | 333.72 |
| | CID2 | -161.61 | 335.23 | 337.10 |
| | CID4 | -161.01 | 338.02 | 341.37 |
| Tree74 | Constant rate | -171.15 | 350.30 | 351.15 |
| | **BiSSE** | **-151.05** | **312.10** | **313.41** |
| | HiSSE | -148.82 | 317.63 | 323.00 |
| | CID2 | -158.12 | 328.24 | 330.10 |
| | CID4 | -160.00 | 335.99 | 339.34 |
| Tree75 | Constant rate | -172.75 | 353.49 | 354.34 |
| | **BiSSE** | **-158.63** | **327.25** | **328.56** |
| | HiSSE | -157.27 | 334.55 | 339.91 |
| | CID2 | -165.98 | 343.97 | 345.84 |
| | CID4 | -161.26 | 338.52 | 341.87 |
| Tree76 | Constant rate | -174.91 | 357.82 | 358.67 |
| | **BiSSE** | **-150.61** | **311.21** | **312.52** |
| | HiSSE | -148.10 | 316.20 | 321.57 |
| | CID2 | -161.46 | 334.91 | 336.78 |
| | CID4 | -154.41 | 324.81 | 328.16 |
| Tree77 | Constant rate | -160.28 | 328.57 | 329.42 |
| | **BiSSE** | **-142.72** | **295.45** | **296.75** |
| | HiSSE | -140.96 | 301.92 | 307.29 |
| | CID2 | -148.86 | 309.73 | 311.59 |
| | CID4 | -146.82 | 309.65 | 313.00 |
| Tree78 | Constant rate | -172.60 | 353.19 | 354.04 |
| | **BiSSE** | **-159.16** | **328.32** | **329.63** |
| | HiSSE | -156.44 | 332.89 | 338.26 |
| | CID2 | -165.34 | 342.67 | 344.54 |
| | CID4 | -160.70 | 337.41 | 340.76 |
| Tree79 | Constant rate | -175.28 | 358.56 | 359.41 |
| | **BiSSE** | **-154.20** | **318.39** | **319.70** |
| | HiSSE | -151.20 | 322.41 | 327.78 |
| | CID2 | -159.75 | 331.49 | 333.36 |
| | CID4 | -158.56 | 333.12 | 336.47 |
| Tree80 | Constant rate | -168.84 | 345.67 | 346.52 |
| | **BiSSE** | **-151.33** | **312.65** | **313.95** |
| | HiSSE | -148.62 | 317.24 | 322.61 |
| | CID2 | -161.66 | 335.32 | 337.19 |
| | CID4 | -153.87 | 323.75 | 327.10 |
| Tree81 | Constant rate | -170.71 | 349.42 | 350.28 |
| | **BiSSE** | **-157.62** | **325.23** | **326.53** |
| | HiSSE | -152.19 | 324.37 | 329.74 |
| | CID2 | -159.92 | 331.83 | 333.70 |
| | CID4 | -160.33 | 336.66 | 340.01 |
| | Constant rate | -173.12 | 354.24 | 355.09 |
| | **BiSSE** | **-150.84** | **311.67** | **312.98** |

| | | | | |
|---|---|---|---|---|
| Tree82 | HiSSE | -151.02 | 322.04 | 327.41 |
| | CID2 | -162.55 | 337.10 | 338.97 |
| | CID4 | -154.71 | 325.43 | 328.78 |
| Tree83 | Constant rate | -171.29 | 350.58 | 351.43 |
| | **BiSSE** | **-157.36** | **324.73** | **326.03** |
| | HiSSE | -154.83 | 329.65 | 335.02 |
| | CID2 | -163.26 | 338.52 | 340.38 |
| | CID4 | -162.43 | 340.85 | 344.20 |
| Tree84 | Constant rate | -166.77 | 341.55 | 342.40 |
| | **BiSSE** | **-143.84** | **297.68** | **298.99** |
| | HiSSE | -141.51 | 303.02 | 308.39 |
| | CID2 | -151.97 | 315.95 | 317.81 |
| | CID4 | -148.17 | 312.33 | 315.68 |
| Tree85 | Constant rate | -167.04 | 342.09 | 342.94 |
| | **BiSSE** | **-153.04** | **316.09** | **317.39** |
| | HiSSE | -147.80 | 315.60 | 320.97 |
| | CID2 | -156.22 | 324.43 | 326.30 |
| | CID4 | -155.94 | 327.88 | 331.23 |
| Tree86 | Constant rate | -164.39 | 336.78 | 337.63 |
| | **BiSSE** | **-152.16** | **314.33** | **315.63** |
| | HiSSE | -149.01 | 318.03 | 323.39 |
| | CID2 | -156.82 | 325.65 | 327.51 |
| | CID4 | -154.55 | 325.11 | 328.46 |
| Tree87 | Constant rate | -174.58 | 357.17 | 358.02 |
| | **BiSSE** | **-159.69** | **329.38** | **330.69** |
| | HiSSE | -159.29 | 338.59 | 343.95 |
| | CID2 | -165.29 | 342.59 | 344.45 |
| | CID4 | -161.54 | 339.09 | 342.44 |
| Tree88 | Constant rate | -161.88 | 331.77 | 332.62 |
| | **BiSSE** | **-144.95** | **299.90** | **301.20** |
| | HiSSE | -144.28 | 308.57 | 313.93 |
| | CID2 | -149.97 | 311.93 | 313.80 |
| | CID4 | -148.17 | 312.34 | 315.68 |
| Tree89 | Constant rate | -177.45 | 362.91 | 363.76 |
| | **BiSSE** | **-162.58** | **335.16** | **336.47** |
| | HiSSE | -159.44 | 338.87 | 344.24 |
| | CID2 | -167.78 | 347.57 | 349.44 |
| | CID4 | -164.04 | 344.08 | 347.43 |
| Tree90 | Constant rate | -172.43 | 352.86 | 353.71 |
| | **BiSSE** | **-155.26** | **320.53** | **321.83** |
| | HiSSE | -154.99 | 329.98 | 335.34 |
| | CID2 | -162.06 | 336.12 | 337.99 |
| | CID4 | -159.25 | 334.50 | 337.85 |
| Tree91 | Constant rate | -166.58 | 341.15 | 342.00 |
| | **BiSSE** | **-149.90** | **309.80** | **311.11** |
| | HiSSE | -147.72 | 315.43 | 320.80 |

| | | | | |
|---|---|---|---|---|
| | CID2 | -155.62 | 323.23 | 325.10 |
| | CID4 | -155.65 | 327.29 | 330.64 |
| Tree92 | Constant rate | -167.60 | 343.21 | 344.06 |
| | **BiSSE** | **-150.81** | **311.63** | **312.93** |
| | HiSSE | -147.09 | 314.17 | 319.54 |
| | CID2 | -160.63 | 333.27 | 335.13 |
| | CID4 | -150.94 | 317.88 | 321.22 |
| Tree93 | Constant rate | -165.16 | 338.33 | 339.18 |
| | **BiSSE** | **-150.96** | **311.93** | **313.23** |
| | HiSSE | -150.04 | 320.08 | 325.45 |
| | CID2 | -157.13 | 326.26 | 328.13 |
| | CID4 | -156.21 | 328.42 | 331.77 |
| Tree94 | Constant rate | -159.80 | 327.60 | 328.45 |
| | **BiSSE** | **-150.02** | **310.05** | **311.35** |
| | HiSSE | -145.33 | 310.65 | 316.02 |
| | CID2 | -152.16 | 316.33 | 318.19 |
| | CID4 | -150.63 | 317.26 | 320.61 |
| Tree95 | Constant rate | -174.19 | 356.38 | 357.23 |
| | **BiSSE** | **-157.33** | **324.66** | **325.96** |
| | HiSSE | -158.16 | 336.32 | 341.69 |
| | CID2 | -163.42 | 338.84 | 340.71 |
| | CID4 | -159.59 | 335.18 | 338.53 |
| Tree96 | Constant rate | -166.71 | 341.42 | 342.27 |
| | **BiSSE** | **-149.16** | **308.32** | **309.62** |
| | HiSSE | -146.33 | 312.66 | 318.03 |
| | CID2 | -155.02 | 322.03 | 323.90 |
| | CID4 | -153.05 | 322.09 | 325.44 |
| Tree97 | Constant rate | -173.56 | 355.12 | 355.97 |
| | **BiSSE** | **-154.18** | **318.35** | **319.66** |
| | HiSSE | -151.49 | 322.98 | 328.34 |
| | CID2 | -161.66 | 335.31 | 337.18 |
| | CID4 | -158.04 | 332.08 | 335.43 |
| Tree98 | Constant rate | -164.94 | 337.88 | 338.73 |
| | **BiSSE** | **-144.81** | **299.62** | **300.92** |
| | HiSSE | -142.48 | 304.96 | 310.33 |
| | CID2 | -150.48 | 312.96 | 314.83 |
| | CID4 | -147.42 | 310.84 | 314.18 |
| Tree99 | Constant rate | -171.67 | 351.34 | 352.19 |
| | **BiSSE** | **-157.21** | **324.41** | **325.72** |
| | HiSSE | -157.21 | 334.41 | 339.78 |
| | CID2 | -163.61 | 339.21 | 341.08 |
| | CID4 | -160.04 | 336.08 | 339.43 |
| Tree100 | Constant rate | -168.01 | 344.01 | 344.87 |
| | **BiSSE** | **-151.82** | **313.64** | **314.95** |
| | HiSSE | -148.28 | 316.56 | 321.92 |
| | CID2 | -157.51 | 327.01 | 328.88 |

| | | | | |
|---|---|---|---|---|
| | CID4 | -156.88 | 329.76 | 333.11 |
