## Supplementary material for "‘Dispersification’ of *Agalinis* (Orobanchaceae) into South America is associated with hummingbird pollination and perennial life history shifts": Table S6.

**Table S6.** Log-likelihood values (lnL), Akaike Information Criterion (AIC), and corrected AIC (AICc) for State Speciation and Extinction (SSE) models associating pollination syndrome and diversification in *Agalinis* for the 100 randomly sampled post–burn-in MCMC trees. The Binary SSE (BiSSE) and Hidden State SSE (HiSSE) models link diversification rates to pollination syndrome, while the constant rate, CID2, and CID4 models assume diversification is independent of this trait.

| File | Model | lnL | AIC | AICc |
|---|---|---|---|---|
| Tree1 | Constant rate | -166.77 | 341.54 | 342.45 |
| | **BiSSE** | **-147.32** | **304.63** | **306.03** |
| | HiSSE | -147.32 | 314.63 | 320.42 |
| | CID2 | -161.51 | 335.02 | 337.02 |
| | CID4 | -153.79 | 323.58 | 327.18 |
| Tree2 | Constant rate | -163.59 | 335.19 | 336.09 |
| | **BiSSE** | **-141.86** | **293.72** | **295.12** |
| | HiSSE | -141.86 | 303.72 | 309.51 |
| | CID2 | -151.97 | 315.93 | 317.93 |
| | CID4 | -149.32 | 314.64 | 318.24 |
| Tree3 | Constant rate | -164.02 | 336.04 | 336.95 |
| | **BiSSE** | **-143.54** | **297.09** | **298.48** |
| | HiSSE | -143.54 | 307.09 | 312.88 |
| | CID2 | -155.48 | 322.96 | 324.96 |
| | CID4 | -150.38 | 316.76 | 320.36 |
| Tree4 | Constant rate | -166.23 | 340.46 | 341.37 |
| | **BiSSE** | **-146.66** | **303.32** | **304.72** |
| | HiSSE | -145.31 | 310.63 | 316.41 |
| | CID2 | -158.20 | 328.40 | 330.40 |
| | CID4 | -154.21 | 324.42 | 328.02 |
| Tree5 | Constant rate | -156.97 | 321.93 | 322.84 |
| | **BiSSE** | **-142.54** | **295.08** | **296.47** |
| | HiSSE | -140.60 | 301.20 | 306.99 |
| | CID2 | -148.91 | 309.82 | 311.82 |
| | CID4 | -147.76 | 311.51 | 315.11 |
| Tree6 | Constant rate | -171.46 | 350.93 | 351.84 |
| | **BiSSE** | **-152.33** | **314.66** | **316.06** |
| | HiSSE | -151.40 | 322.80 | 328.59 |
| | CID2 | -163.21 | 338.43 | 340.43 |
| | CID4 | -158.97 | 333.94 | 337.54 |
| Tree7 | Constant rate | -165.74 | 339.48 | 340.39 |
| | **BiSSE** | **-140.96** | **291.91** | **293.31** |
| | HiSSE | -140.96 | 301.91 | 307.70 |
| | CID2 | -151.43 | 314.85 | 316.85 |
| | CID4 | -149.72 | 315.43 | 319.03 |

| | | | | |
|---|---|---|---|---|
| Tree8 | Constant rate | -170.50 | 349.01 | 349.92 |
| | **BiSSE** | **-150.95** | **311.91** | **313.30** |
| | HiSSE | -150.95 | 321.91 | 327.70 |
| | CID2 | -163.97 | 339.93 | 341.93 |
| | CID4 | -158.38 | 332.76 | 336.36 |
| Tree9 | Constant rate | -160.27 | 328.54 | 329.45 |
| | **BiSSE** | **-144.60** | **299.19** | **300.59** |
| | HiSSE | -143.07 | 306.14 | 311.93 |
| | CID2 | -151.76 | 315.52 | 317.52 |
| | CID4 | -148.72 | 313.43 | 317.03 |
| Tree10 | Constant rate | -169.61 | 347.22 | 348.13 |
| | **BiSSE** | **-147.85** | **305.69** | **307.09** |
| | HiSSE | -147.85 | 315.69 | 321.48 |
| | CID2 | -159.90 | 331.80 | 333.80 |
| | CID4 | -155.61 | 327.23 | 330.83 |
| Tree11 | Constant rate | -166.37 | 340.75 | 341.66 |
| | **BiSSE** | **-145.16** | **300.33** | **301.73** |
| | HiSSE | -143.91 | 307.81 | 313.60 |
| | CID2 | -155.70 | 323.41 | 325.41 |
| | CID4 | -152.22 | 320.43 | 324.03 |
| Tree12 | Constant rate | -167.19 | 342.39 | 343.30 |
| | **BiSSE** | **-147.24** | **304.48** | **305.87** |
| | HiSSE | -147.24 | 314.48 | 320.27 |
| | CID2 | -157.63 | 327.26 | 329.26 |
| | CID4 | -153.44 | 322.87 | 326.47 |
| Tree13 | Constant rate | -166.08 | 340.17 | 341.08 |
| | **BiSSE** | **-146.78** | **303.56** | **304.95** |
| | HiSSE | -146.78 | 313.56 | 319.35 |
| | CID2 | -158.21 | 328.42 | 330.42 |
| | CID4 | -154.21 | 324.43 | 328.03 |
| Tree14 | Constant rate | -165.30 | 338.60 | 339.51 |
| | **BiSSE** | **-145.95** | **301.90** | **303.30** |
| | HiSSE | -145.95 | 311.90 | 317.69 |
| | CID2 | -156.47 | 324.94 | 326.94 |
| | CID4 | -153.00 | 321.99 | 325.59 |
| Tree15 | Constant rate | -166.40 | 340.80 | 341.71 |
| | **BiSSE** | **-142.52** | **295.04** | **296.44** |
| | HiSSE | -142.52 | 305.04 | 310.83 |
| | CID2 | -153.62 | 319.25 | 321.25 |
| | CID4 | -149.65 | 315.29 | 318.89 |
| | Constant rate | -166.85 | 341.71 | 342.62 |

| | | | | |
|---|---|---|---|---|
| Tree16 | **BiSSE** | **-148.53** | **307.06** | **308.46** |
| | HiSSE | -148.17 | 316.33 | 322.12 |
| | CID2 | -160.29 | 332.57 | 334.57 |
| | CID4 | -154.43 | 324.86 | 328.46 |
| Tree17 | Constant rate | -161.83 | 331.66 | 332.57 |
| | **BiSSE** | **-142.29** | **294.57** | **295.97** |
| | HiSSE | -141.72 | 303.44 | 309.23 |
| | CID2 | -154.27 | 320.55 | 322.55 |
| | CID4 | -150.53 | 317.06 | 320.66 |
| Tree18 | Constant rate | -159.14 | 326.29 | 327.20 |
| | **BiSSE** | **-140.89** | **291.77** | **293.17** |
| | HiSSE | -140.76 | 301.52 | 307.31 |
| | CID2 | -151.41 | 314.82 | 316.82 |
| | CID4 | -148.81 | 313.62 | 317.22 |
| Tree19 | Constant rate | -166.79 | 341.57 | 342.48 |
| | **BiSSE** | **-145.94** | **301.88** | **303.28** |
| | HiSSE | -145.94 | 311.88 | 317.67 |
| | CID2 | -156.78 | 325.55 | 327.55 |
| | CID4 | -153.41 | 322.83 | 326.43 |
| Tree20 | Constant rate | -164.58 | 337.15 | 338.06 |
| | **BiSSE** | **-147.28** | **304.57** | **305.96** |
| | HiSSE | -146.85 | 313.69 | 319.48 |
| | CID2 | -158.69 | 329.38 | 331.38 |
| | CID4 | -155.57 | 327.14 | 330.74 |
| Tree21 | Constant rate | -164.48 | 336.95 | 337.86 |
| | **BiSSE** | **-145.10** | **300.20** | **301.59** |
| | HiSSE | -144.75 | 309.49 | 315.28 |
| | CID2 | -155.30 | 322.60 | 324.60 |
| | CID4 | -151.95 | 319.89 | 323.49 |
| Tree22 | Constant rate | -168.01 | 344.02 | 344.93 |
| | **BiSSE** | **-144.71** | **299.41** | **300.81** |
| | HiSSE | -144.71 | 309.41 | 315.20 |
| | CID2 | -153.62 | 319.23 | 321.23 |
| | CID4 | -156.87 | 329.74 | 333.34 |
| Tree23 | Constant rate | -166.32 | 340.64 | 341.55 |
| | **BiSSE** | **-145.32** | **300.63** | **302.03** |
| | HiSSE | -145.32 | 310.63 | 316.42 |
| | CID2 | -157.19 | 326.38 | 328.38 |
| | CID4 | -153.26 | 322.51 | 326.11 |
| | Constant rate | -166.16 | 340.31 | 341.22 |
| | **BiSSE** | **-151.09** | **312.18** | **313.58** |

| | | | | |
|---|---|---|---|---|
| Tree24 | HiSSE | -151.09 | 322.18 | 327.97 |
| | CID2 | -160.64 | 333.27 | 335.27 |
| | CID4 | -157.83 | 331.67 | 335.27 |
| Tree25 | Constant rate | -163.32 | 334.63 | 335.54 |
| | **BiSSE** | **-140.15** | **290.31** | **291.70** |
| | HiSSE | -139.89 | 299.77 | 305.56 |
| | CID2 | -153.26 | 318.51 | 320.51 |
| | CID4 | -150.14 | 316.29 | 319.89 |
| Tree26 | Constant rate | -165.10 | 338.19 | 339.10 |
| | **BiSSE** | **-145.78** | **301.56** | **302.95** |
| | HiSSE | -145.78 | 311.56 | 317.35 |
| | CID2 | -159.30 | 330.59 | 332.59 |
| | CID4 | -153.70 | 323.40 | 327.00 |
| Tree27 | Constant rate | -164.45 | 336.90 | 337.81 |
| | **BiSSE** | **-145.64** | **301.28** | **302.68** |
| | HiSSE | -145.64 | 311.28 | 317.07 |
| | CID2 | -156.75 | 325.49 | 327.49 |
| | CID4 | -153.96 | 323.92 | 327.52 |
| Tree28 | Constant rate | -165.36 | 338.73 | 339.64 |
| | **BiSSE** | **-145.32** | **300.63** | **302.03** |
| | HiSSE | -145.32 | 310.63 | 316.42 |
| | CID2 | -156.77 | 325.55 | 327.55 |
| | CID4 | -152.13 | 320.25 | 323.85 |
| Tree29 | Constant rate | -165.51 | 339.03 | 339.94 |
| | **BiSSE** | **-149.10** | **308.21** | **309.60** |
| | HiSSE | -149.10 | 318.21 | 324.00 |
| | CID2 | -159.39 | 330.77 | 332.77 |
| | CID4 | -156.34 | 328.68 | 332.28 |
| Tree30 | Constant rate | -170.43 | 348.86 | 349.77 |
| | **BiSSE** | **-151.16** | **312.31** | **313.71** |
| | HiSSE | -149.82 | 319.64 | 325.43 |
| | CID2 | -162.39 | 336.78 | 338.78 |
| | CID4 | -158.44 | 332.89 | 336.49 |
| Tree31 | Constant rate | -173.75 | 355.51 | 356.42 |
| | **BiSSE** | **-151.31** | **312.61** | **314.01** |
| | HiSSE | -151.31 | 322.61 | 328.40 |
| | CID2 | -163.33 | 338.67 | 340.67 |
| | CID4 | -158.77 | 333.55 | 337.15 |
| Tree32 | Constant rate | -169.72 | 347.43 | 348.34 |
| | **BiSSE** | **-149.75** | **309.50** | **310.89** |
| | HiSSE | -149.75 | 319.50 | 325.29 |

| | | | | |
|---|---|---|---|---|
| | CID2 | -160.98 | 333.96 | 335.96 |
| | CID4 | -157.40 | 330.79 | 334.39 |
| Tree33 | Constant rate | -170.67 | 349.33 | 350.24 |
| | **BiSSE** | **-148.81** | **307.61** | **309.01** |
| | HiSSE | -148.38 | 316.77 | 322.56 |
| | CID2 | -160.23 | 332.46 | 334.46 |
| | CID4 | -154.64 | 325.29 | 328.89 |
| Tree34 | Constant rate | -161.36 | 330.71 | 331.62 |
| | **BiSSE** | **-141.22** | **292.44** | **293.84** |
| | HiSSE | -140.06 | 300.12 | 305.91 |
| | CID2 | -148.81 | 309.62 | 311.62 |
| | CID4 | -147.61 | 311.22 | 314.82 |
| Tree35 | Constant rate | -164.50 | 336.99 | 337.90 |
| | **BiSSE** | **-145.84** | **301.68** | **303.08** |
| | HiSSE | -144.50 | 309.00 | 314.79 |
| | CID2 | -154.73 | 321.46 | 323.46 |
| | CID4 | -151.99 | 319.98 | 323.58 |
| Tree36 | Constant rate | -164.18 | 336.36 | 337.27 |
| | **BiSSE** | **-145.02** | **300.03** | **301.43** |
| | HiSSE | -145.02 | 310.03 | 315.82 |
| | CID2 | -156.43 | 324.85 | 326.85 |
| | CID4 | -152.88 | 321.75 | 325.35 |
| Tree37 | Constant rate | -169.01 | 346.03 | 346.94 |
| | **BiSSE** | **-146.35** | **302.71** | **304.10** |
| | HiSSE | -144.32 | 308.64 | 314.43 |
| | CID2 | -160.65 | 333.30 | 335.30 |
| | CID4 | -155.25 | 326.50 | 330.10 |
| Tree38 | Constant rate | -166.00 | 339.99 | 340.90 |
| | **BiSSE** | **-147.94** | **305.89** | **307.28** |
| | HiSSE | -147.33 | 314.66 | 320.45 |
| | CID2 | -158.22 | 328.45 | 330.45 |
| | CID4 | -152.70 | 321.40 | 325.00 |
| Tree39 | Constant rate | -164.28 | 336.56 | 337.47 |
| | **BiSSE** | **-148.05** | **306.09** | **307.49** |
| | HiSSE | -146.48 | 312.97 | 318.76 |
| | CID2 | -157.76 | 327.52 | 329.52 |
| | CID4 | -153.16 | 322.32 | 325.92 |
| Tree40 | Constant rate | -164.47 | 336.95 | 337.86 |
| | **BiSSE** | **-143.96** | **297.92** | **299.32** |
| | HiSSE | -142.37 | 304.73 | 310.52 |
| | CID2 | -154.40 | 320.79 | 322.79 |

| | | | | |
|---|---|---|---|---|
| | CID4 | -150.38 | 316.76 | 320.36 |
| Tree41 | Constant rate | -165.03 | 338.06 | 338.97 |
| | **BiSSE** | **-145.38** | **300.76** | **302.16** |
| | HiSSE | -142.54 | 305.08 | 310.87 |
| | CID2 | -154.45 | 320.90 | 322.90 |
| | CID4 | -151.38 | 318.77 | 322.37 |
| Tree42 | Constant rate | -161.25 | 330.51 | 331.42 |
| | **BiSSE** | **-144.60** | **299.21** | **300.60** |
| | HiSSE | -142.10 | 304.19 | 309.98 |
| | CID2 | -151.27 | 314.54 | 316.54 |
| | CID4 | -151.44 | 318.88 | 322.48 |
| Tree43 | Constant rate | -168.10 | 344.20 | 345.11 |
| | **BiSSE** | **-147.73** | **305.46** | **306.86** |
| | HiSSE | -147.09 | 314.17 | 319.96 |
| | CID2 | -158.44 | 328.88 | 330.88 |
| | CID4 | -155.05 | 326.10 | 329.70 |
| Tree44 | Constant rate | -163.07 | 334.15 | 335.06 |
| | **BiSSE** | **-146.23** | **302.47** | **303.87** |
| | HiSSE | -144.99 | 309.97 | 315.76 |
| | CID2 | -155.16 | 322.32 | 324.32 |
| | CID4 | -152.06 | 320.12 | 323.72 |
| Tree45 | Constant rate | -163.14 | 334.27 | 335.18 |
| | **BiSSE** | **-150.55** | **311.11** | **312.50** |
| | HiSSE | -150.55 | 321.11 | 326.90 |
| | CID2 | -157.05 | 326.11 | 328.11 |
| | CID4 | -154.26 | 324.51 | 328.11 |
| Tree46 | Constant rate | -162.83 | 333.66 | 334.57 |
| | **BiSSE** | **-144.56** | **299.11** | **300.51** |
| | HiSSE | -144.34 | 308.68 | 314.47 |
| | CID2 | -153.13 | 318.26 | 320.26 |
| | CID4 | -150.36 | 316.71 | 320.31 |
| Tree47 | Constant rate | -165.96 | 339.91 | 340.82 |
| | **BiSSE** | **-145.25** | **300.50** | **301.89** |
| | HiSSE | -145.25 | 310.50 | 316.29 |
| | CID2 | -157.59 | 327.19 | 329.19 |
| | CID4 | -153.77 | 323.53 | 327.13 |
| Tree48 | Constant rate | -161.78 | 331.56 | 332.47 |
| | **BiSSE** | **-146.29** | **302.59** | **303.98** |
| | HiSSE | -143.84 | 307.69 | 313.48 |
| | CID2 | -153.92 | 319.85 | 321.85 |
| | CID4 | -151.56 | 319.12 | 322.72 |

| | | | | |
|---|---|---|---|---|
| Tree49 | Constant rate | -169.30 | 346.61 | 347.51 |
| | **BiSSE** | **-150.77** | **311.55** | **312.94** |
| | HiSSE | -150.77 | 321.55 | 327.34 |
| | CID2 | -160.89 | 333.77 | 335.77 |
| | CID4 | -157.09 | 330.18 | 333.78 |
| Tree50 | Constant rate | -164.49 | 336.99 | 337.90 |
| | **BiSSE** | **-146.04** | **302.07** | **303.47** |
| | HiSSE | -145.32 | 310.65 | 316.44 |
| | CID2 | -157.12 | 326.23 | 328.23 |
| | CID4 | -154.16 | 324.31 | 327.91 |
| Tree51 | Constant rate | -164.97 | 337.93 | 338.84 |
| | **BiSSE** | **-150.48** | **310.96** | **312.36** |
| | HiSSE | -149.73 | 319.46 | 325.25 |
| | CID2 | -157.35 | 326.71 | 328.71 |
| | CID4 | -154.21 | 324.42 | 328.02 |
| Tree52 | Constant rate | -160.07 | 328.14 | 329.05 |
| | **BiSSE** | **-147.73** | **305.46** | **306.86** |
| | HiSSE | -146.73 | 313.46 | 319.24 |
| | CID2 | -157.25 | 326.49 | 328.49 |
| | CID4 | -154.24 | 324.48 | 328.08 |
| Tree53 | Constant rate | -160.47 | 328.93 | 329.84 |
| | **BiSSE** | **-143.03** | **296.07** | **297.46** |
| | HiSSE | -140.45 | 300.90 | 306.69 |
| | CID2 | -151.96 | 315.92 | 317.92 |
| | CID4 | -147.89 | 311.77 | 315.37 |
| Tree54 | Constant rate | -159.58 | 327.17 | 328.08 |
| | **BiSSE** | **-143.02** | **296.04** | **297.43** |
| | HiSSE | -142.34 | 304.68 | 310.47 |
| | CID2 | -151.27 | 314.54 | 316.54 |
| | CID4 | -148.59 | 313.19 | 316.79 |
| Tree55 | Constant rate | -166.18 | 340.36 | 341.27 |
| | **BiSSE** | **-147.96** | **305.91** | **307.31** |
| | HiSSE | -147.96 | 315.91 | 321.70 |
| | CID2 | -158.03 | 328.06 | 330.06 |
| | CID4 | -154.63 | 325.27 | 328.87 |
| Tree56 | Constant rate | -167.44 | 342.89 | 343.80 |
| | **BiSSE** | **-144.02** | **298.03** | **299.43** |
| | HiSSE | -144.02 | 308.03 | 313.82 |
| | CID2 | -155.08 | 322.15 | 324.15 |
| | CID4 | -151.00 | 318.00 | 321.60 |
| | Constant rate | -160.31 | 328.62 | 329.52 |

| | | | | |
|---|---|---|---|---|
| Tree57 | **BiSSE** | **-143.09** | **296.18** | **297.58** |
| | HiSSE | -142.53 | 305.06 | 310.85 |
| | CID2 | -152.58 | 317.16 | 319.16 |
| | CID4 | -149.40 | 314.79 | 318.39 |
| Tree58 | Constant rate | -165.59 | 339.19 | 340.10 |
| | **BiSSE** | **-145.75** | **301.50** | **302.89** |
| | HiSSE | -144.73 | 309.46 | 315.25 |
| | CID2 | -155.59 | 323.19 | 325.19 |
| | CID4 | -151.87 | 319.74 | 323.34 |
| Tree59 | Constant rate | -166.66 | 341.32 | 342.23 |
| | **BiSSE** | **-142.20** | **294.40** | **295.80** |
| | HiSSE | -140.41 | 300.82 | 306.61 |
| | CID2 | -154.40 | 320.81 | 322.81 |
| | CID4 | -152.21 | 320.42 | 324.02 |
| Tree60 | Constant rate | -168.99 | 345.98 | 346.89 |
| | **BiSSE** | **-150.10** | **310.20** | **311.59** |
| | HiSSE | -149.56 | 319.12 | 324.91 |
| | CID2 | -163.27 | 338.54 | 340.54 |
| | CID4 | -158.04 | 332.08 | 335.68 |
| Tree61 | Constant rate | -162.70 | 333.40 | 334.30 |
| | **BiSSE** | **-145.17** | **300.35** | **301.74** |
| | HiSSE | -145.01 | 310.02 | 315.81 |
| | CID2 | -154.87 | 321.74 | 323.74 |
| | CID4 | -150.93 | 317.85 | 321.45 |
| Tree62 | Constant rate | -166.87 | 341.74 | 342.65 |
| | **BiSSE** | **-148.98** | **307.95** | **309.35** |
| | HiSSE | -148.98 | 317.95 | 323.74 |
| | CID2 | -160.16 | 332.33 | 334.33 |
| | CID4 | -156.36 | 328.72 | 332.32 |
| Tree63 | Constant rate | -159.65 | 327.31 | 328.22 |
| | **BiSSE** | **-141.75** | **293.50** | **294.89** |
| | HiSSE | -140.11 | 300.21 | 306.00 |
| | CID2 | -152.73 | 317.45 | 319.45 |
| | CID4 | -148.61 | 313.21 | 316.81 |
| Tree64 | Constant rate | -165.57 | 339.14 | 340.05 |
| | **BiSSE** | **-148.79** | **307.59** | **308.98** |
| | HiSSE | -146.62 | 313.24 | 319.03 |
| | CID2 | -158.26 | 328.53 | 330.53 |
| | CID4 | -155.28 | 326.56 | 330.16 |
| | Constant rate | -160.92 | 329.83 | 330.74 |
| | **BiSSE** | **-145.22** | **300.44** | **301.84** |

| | | | | |
|---|---|---|---|---|
| Tree65 | HiSSE | -144.40 | 308.79 | 314.58 |
| | CID2 | -156.87 | 325.74 | 327.74 |
| | CID4 | -151.53 | 319.07 | 322.67 |
| Tree66 | Constant rate | -161.63 | 331.26 | 332.17 |
| | **BiSSE** | **-145.37** | **300.75** | **302.14** |
| | HiSSE | -145.37 | 310.75 | 316.53 |
| | CID2 | -154.98 | 321.97 | 323.97 |
| | CID4 | -152.37 | 320.73 | 324.33 |
| Tree67 | Constant rate | -168.24 | 344.49 | 345.39 |
| | **BiSSE** | **-144.43** | **298.87** | **300.26** |
| | HiSSE | -143.85 | 307.70 | 313.49 |
| | CID2 | -161.45 | 334.91 | 336.91 |
| | CID4 | -152.64 | 321.28 | 324.88 |
| Tree68 | Constant rate | -159.12 | 326.23 | 327.14 |
| | **BiSSE** | **-140.82** | **291.64** | **293.03** |
| | HiSSE | -140.82 | 301.64 | 307.43 |
| | CID2 | -152.86 | 317.72 | 319.72 |
| | CID4 | -148.36 | 312.71 | 316.31 |
| Tree69 | Constant rate | -167.94 | 343.88 | 344.79 |
| | **BiSSE** | **-148.05** | **306.10** | **307.49** |
| | HiSSE | -145.85 | 311.71 | 317.50 |
| | CID2 | -158.53 | 329.06 | 331.06 |
| | CID4 | -154.70 | 325.40 | 329.00 |
| Tree70 | Constant rate | -161.83 | 331.67 | 332.57 |
| | **BiSSE** | **-142.37** | **294.75** | **296.14** |
| | HiSSE | -140.27 | 300.53 | 306.32 |
| | CID2 | -155.40 | 322.79 | 324.79 |
| | CID4 | -149.52 | 315.03 | 318.63 |
| Tree71 | Constant rate | -167.01 | 342.02 | 342.93 |
| | **BiSSE** | **-146.93** | **303.86** | **305.26** |
| | HiSSE | -146.41 | 312.81 | 318.60 |
| | CID2 | -156.57 | 325.13 | 327.13 |
| | CID4 | -153.40 | 322.81 | 326.41 |
| Tree72 | Constant rate | -158.70 | 325.40 | 326.31 |
| | **BiSSE** | **-139.91** | **289.83** | **291.22** |
| | HiSSE | -139.41 | 298.82 | 304.61 |
| | CID2 | -148.85 | 309.71 | 311.71 |
| | CID4 | -145.50 | 306.99 | 310.59 |
| Tree73 | Constant rate | -164.27 | 336.53 | 337.44 |
| | **BiSSE** | **-146.81** | **303.62** | **305.02** |
| | HiSSE | -146.22 | 312.44 | 318.23 |

| | | | | |
|---|---|---|---|---|
| | CID2 | -157.02 | 326.04 | 328.04 |
| | CID4 | -156.06 | 328.12 | 331.72 |
| Tree74 | Constant rate | -163.28 | 334.56 | 335.47 |
| | **BiSSE** | **-142.64** | **295.28** | **296.67** |
| | HiSSE | -141.79 | 303.58 | 309.37 |
| | CID2 | -155.50 | 322.99 | 324.99 |
| | CID4 | -149.08 | 314.15 | 317.75 |
| Tree75 | Constant rate | -164.19 | 336.39 | 337.30 |
| | **BiSSE** | **-147.50** | **305.00** | **306.39** |
| | HiSSE | -146.90 | 313.80 | 319.59 |
| | CID2 | -160.03 | 332.05 | 334.05 |
| | CID4 | -154.25 | 324.49 | 328.09 |
| Tree76 | Constant rate | -165.68 | 339.35 | 340.26 |
| | **BiSSE** | **-143.43** | **296.86** | **298.25** |
| | HiSSE | -141.41 | 302.82 | 308.61 |
| | CID2 | -151.96 | 315.93 | 317.93 |
| | CID4 | -148.26 | 312.51 | 316.11 |
| Tree77 | Constant rate | -153.49 | 314.99 | 315.90 |
| | **BiSSE** | **-136.99** | **283.97** | **285.37** |
| | HiSSE | -136.09 | 292.18 | 297.97 |
| | CID2 | -144.28 | 300.57 | 302.57 |
| | CID4 | -142.13 | 300.26 | 303.86 |
| Tree78 | Constant rate | -166.15 | 340.30 | 341.21 |
| | **BiSSE** | **-150.72** | **311.44** | **312.84** |
| | HiSSE | -149.78 | 319.56 | 325.35 |
| | CID2 | -160.38 | 332.75 | 334.75 |
| | CID4 | -156.47 | 328.94 | 332.54 |
| Tree79 | Constant rate | -166.74 | 341.47 | 342.38 |
| | **BiSSE** | **-141.77** | **293.54** | **294.93** |
| | HiSSE | -141.77 | 303.54 | 309.33 |
| | CID2 | -152.08 | 316.17 | 318.17 |
| | CID4 | -150.32 | 316.64 | 320.24 |
| Tree80 | Constant rate | -162.53 | 333.06 | 333.97 |
| | **BiSSE** | **-141.25** | **292.50** | **293.89** |
| | HiSSE | -141.25 | 302.50 | 308.29 |
| | CID2 | -154.04 | 320.08 | 322.08 |
| | CID4 | -150.23 | 316.45 | 320.05 |
| Tree81 | Constant rate | -163.49 | 334.98 | 335.89 |
| | **BiSSE** | **-145.08** | **300.16** | **301.55** |
| | HiSSE | -145.08 | 310.16 | 315.95 |
| | CID2 | -155.70 | 323.40 | 325.40 |

| | | | | |
|---|---|---|---|---|
| | CID4 | -153.21 | 322.42 | 326.02 |
| Tree82 | Constant rate | -166.58 | 341.16 | 342.07 |
| | **BiSSE** | **-143.76** | **297.52** | **298.91** |
| | HiSSE | -143.76 | 307.52 | 313.31 |
| | CID2 | -158.68 | 329.36 | 331.36 |
| | CID4 | -150.67 | 317.34 | 320.94 |
| Tree83 | Constant rate | -164.64 | 337.28 | 338.19 |
| | **BiSSE** | **-148.61** | **307.23** | **308.63** |
| | HiSSE | -147.78 | 315.56 | 321.35 |
| | CID2 | -158.64 | 329.28 | 331.28 |
| | CID4 | -155.79 | 327.58 | 331.18 |
| Tree84 | Constant rate | -158.32 | 324.65 | 325.56 |
| | **BiSSE** | **-136.86** | **283.72** | **285.11** |
| | HiSSE | -136.86 | 293.72 | 299.51 |
| | CID2 | -147.03 | 306.06 | 308.06 |
| | CID4 | -142.44 | 300.89 | 304.49 |
| Tree85 | Constant rate | -159.56 | 327.11 | 328.02 |
| | **BiSSE** | **-141.84** | **293.69** | **295.08** |
| | HiSSE | -141.84 | 303.69 | 309.47 |
| | CID2 | -151.13 | 314.25 | 316.25 |
| | CID4 | -148.71 | 313.42 | 317.02 |
| Tree86 | Constant rate | -157.83 | 323.65 | 324.56 |
| | **BiSSE** | **-144.11** | **298.22** | **299.62** |
| | HiSSE | -142.48 | 304.96 | 310.75 |
| | CID2 | -151.71 | 315.41 | 317.41 |
| | CID4 | -149.52 | 315.04 | 318.64 |
| Tree87 | Constant rate | -166.72 | 341.43 | 342.34 |
| | **BiSSE** | **-147.38** | **304.77** | **306.16** |
| | HiSSE | -146.83 | 313.66 | 319.45 |
| | CID2 | -159.20 | 330.41 | 332.41 |
| | CID4 | -155.68 | 327.36 | 330.96 |
| Tree88 | Constant rate | -154.75 | 317.49 | 318.40 |
| | **BiSSE** | **-139.94** | **289.88** | **291.28** |
| | HiSSE | -139.94 | 299.88 | 305.67 |
| | CID2 | -148.07 | 308.14 | 310.14 |
| | CID4 | -143.71 | 303.42 | 307.02 |
| Tree89 | Constant rate | -167.85 | 343.70 | 344.61 |
| | **BiSSE** | **-150.18** | **310.37** | **311.76** |
| | HiSSE | -150.18 | 320.37 | 326.16 |
| | CID2 | -160.32 | 332.64 | 334.64 |
| | CID4 | -156.76 | 329.52 | 333.12 |

| | | | | |
|---|---|---|---|---|
| Tree90 | Constant rate | -165.17 | 338.33 | 339.24 |
| | **BiSSE** | **-147.37** | **304.75** | **306.14** |
| | HiSSE | -145.90 | 311.80 | 317.58 |
| | CID2 | -157.67 | 327.35 | 329.35 |
| | CID4 | -156.52 | 329.04 | 332.64 |
| Tree91 | Constant rate | -160.02 | 328.05 | 328.96 |
| | **BiSSE** | **-143.68** | **297.36** | **298.75** |
| | HiSSE | -142.00 | 304.01 | 309.79 |
| | CID2 | -152.73 | 317.45 | 319.45 |
| | CID4 | -151.83 | 319.65 | 323.25 |
| Tree92 | Constant rate | -159.15 | 326.30 | 327.21 |
| | **BiSSE** | **-137.60** | **285.20** | **286.59** |
| | HiSSE | -136.62 | 293.24 | 299.03 |
| | CID2 | -149.90 | 311.81 | 313.81 |
| | CID4 | -144.63 | 305.26 | 308.86 |
| Tree93 | Constant rate | -157.79 | 323.59 | 324.50 |
| | **BiSSE** | **-141.39** | **292.78** | **294.17** |
| | HiSSE | -139.60 | 299.21 | 305.00 |
| | CID2 | -152.31 | 316.62 | 318.62 |
| | CID4 | -150.26 | 316.52 | 320.12 |
| Tree94 | Constant rate | -154.36 | 316.71 | 317.62 |
| | **BiSSE** | **-143.96** | **297.92** | **299.32** |
| | HiSSE | -140.93 | 301.85 | 307.64 |
| | CID2 | -148.37 | 308.74 | 310.74 |
| | CID4 | -147.02 | 310.03 | 313.63 |
| Tree95 | Constant rate | -165.92 | 339.85 | 340.76 |
| | **BiSSE** | **-145.98** | **301.96** | **303.35** |
| | HiSSE | -145.98 | 311.96 | 317.75 |
| | CID2 | -157.51 | 327.03 | 329.03 |
| | CID4 | -153.14 | 322.29 | 325.89 |
| Tree96 | Constant rate | -160.55 | 329.09 | 330.00 |
| | **BiSSE** | **-140.52** | **291.04** | **292.43** |
| | HiSSE | -140.52 | 301.04 | 306.83 |
| | CID2 | -151.88 | 315.75 | 317.75 |
| | CID4 | -149.74 | 315.48 | 319.08 |
| Tree97 | Constant rate | -166.10 | 340.19 | 341.10 |
| | **BiSSE** | **-144.96** | **299.92** | **301.31** |
| | HiSSE | -144.96 | 309.92 | 315.71 |
| | CID2 | -159.62 | 331.24 | 333.24 |
| | CID4 | -153.46 | 322.91 | 326.51 |
| | Constant rate | -157.42 | 322.85 | 323.76 |

| | | | | |
|---|---|---|---|---|
| Tree98 | **BiSSE** | **-136.37** | **282.75** | **284.14** |
| | HiSSE | -135.86 | 291.71 | 297.50 |
| | CID2 | -146.13 | 304.26 | 306.26 |
| | CID4 | -146.50 | 308.99 | 312.59 |
| Tree99 | Constant rate | -163.81 | 335.62 | 336.53 |
| | **BiSSE** | **-147.16** | **304.31** | **305.71** |
| | HiSSE | -145.85 | 311.71 | 317.50 |
| | CID2 | -157.64 | 327.28 | 329.28 |
| | CID4 | -154.53 | 325.07 | 328.67 |
| Tree100 | Constant rate | -161.98 | 331.96 | 332.87 |
| | BiSSE | **-143.54** | **297.07** | **298.47** |
| | HiSSE | -142.65 | 305.31 | 311.10 |
| | CID2 | -153.79 | 319.58 | 321.58 |
| | CID4 | -151.14 | 318.28 | 321.88 |
