## Supplementary material for "‘Dispersification’ of *Agalinis* (Orobanchaceae) into South America is associated with hummingbird pollination and perennial life history shifts": Table S7.

**Table S7.** Log-likelihood values (lnL), Akaike Information Criterion (AIC), and corrected AIC (AICc) for Multistate Hidden State Speciation and Extinction (MuHiSSE) models combining pollination syndrome and life history strategy with diversification in *Agalinis* for the congruification tree, r8s tree, BEAST MCC nuclear and plastid trees, and Cluster 1–10 trees. Categories are bee-pollinated annual species (00), bee-pollinated perennial species (01), hummingbird-pollinated annual species (10), and hummingbird-pollinated perennial species (11). The constant rate and MuCID2 models assume diversification is independent of these traits. Models NoHidden1, NoHidden2, and NoHidden3 do not include hidden states, while MuHiSSE1, MuHiSSE2, and MuHiSSE3 include hidden states A and B. Model NoHidden1 associates diversification with life history strategy, NoHidden2 associates diversification with pollination syndrome, and NoHidden3 assigns unique turnover rates to each category. The corresponding MuHiSSE models include hidden states for each of these trait associations.

| **Tree** | **Model** | **lnL** | **AIC** | **AICc** | **turnover 00A** | **turnover 01A** | **turnover 10A** | **turnover 11A** | **turnover 00B** | **turnover 01B** | **turnover 10B** | **turnover 11B** |
| --- | --- | --- | --- | --- | --- | --- | --- | --- | --- | --- | --- | --- |
| **Congruification** | Constant rate | -176.14 | 372.28 | 378.07 | 0.14 | 0.14 | 0.14 | 0.14 | NA | NA | NA | NA |
|  | NoHidden1 | -166.71 | 355.43 | 362.56 | 0.11 | 0.55 | 0.11 | 0.55 | NA | NA | NA | NA |
|  | NoHidden2 | -164.85 | 351.71 | 358.84 | 0.11 | 0.11 | 0.76 | 0.76 | NA | NA | NA | NA |
|  | NoHidden3 | -165.50 | 357.00 | 367.40 | 0.11 | 0.73 | 0.00 | 0.00 | NA | NA | NA | NA |
|  | **MuHiSSE1** | **-157.35** | **342.69** | **355.05** | **0.06** | **0.06** | **0.91** | **0.91** | **0.07** | **0.07** | **0.00** | **0.00** |
|  | MuHiSSE2 | -162.06 | 352.12 | 364.48 | 0.00 | 0.00 | 0.00 | 0.00 | 0.18 | 0.64 | 0.18 | 0.64 |
|  | MuHiSSE3 | -162.14 | 376.28 | 440.10 | 0.11 | 0.00 | 0.00 | 0.00 | 0.16 | 0.80 | 0.00 | 0.05 |
|  | MuCID2 | -170.42 | 364.85 | 373.51 | 0.44 | 0.44 | 0.44 | 0.44 | 0.11 | 0.11 | 0.11 | 0.11 |
| **r8s** | Constant rate | -183.29 | 386.57 | 392.36 | 0.12 | 0.12 | 0.12 | 0.12 | NA | NA | NA | NA |
|  | NoHidden1 | -176.83 | 375.66 | 382.80 | 0.09 | 0.32 | 0.09 | 0.32 | NA | NA | NA | NA |
|  | **NoHidden2** | **-174.53** | **371.05** | **378.19** | **0.09** | **0.09** | **0.47** | **0.47** | **NA** | **NA** | **NA** | **NA** |
|  | NoHidden3 | -176.14 | 378.28 | 388.68 | 0.09 | 0.47 | 0.00 | 0.00 | NA | NA | NA | NA |
|  | MuHiSSE1 | -176.18 | 380.36 | 392.72 | 0.16 | 0.16 | 0.32 | 0.32 | 0.00 | 0.00 | 0.00 | 0.00 |
|  | MuHiSSE2 | -171.30 | 370.60 | 382.95 | 0.15 | 0.00 | 0.15 | 0.00 | 0.00 | 0.42 | 0.00 | 0.42 |
|  | MuHiSSE3 | -169.27 | 390.54 | 454.36 | 0.00 | 0.00 | 0.00 | 0.00 | 0.15 | 0.72 | 0.10 | 0.00 |
|  | MuCID2 | -175.56 | 375.12 | 383.78 | 0.19 | 0.19 | 0.19 | 0.19 | 0.00 | 0.00 | 0.00 | 0.00 |
| **BEAST MCC nuclear** | Constant rate | -173.31 | 366.62 | 372.41 | 0.15 | 0.15 | 0.15 | 0.15 | NA | NA | NA | NA |
|  | NoHidden1 | -165.04 | 352.09 | 359.22 | 0.11 | 0.43 | 0.11 | 0.43 | NA | NA | NA | NA |
|  | NoHidden2 | -164.68 | 351.36 | 358.49 | 0.11 | 0.11 | 0.62 | 0.62 | NA | NA | NA | NA |
|  | **NoHidden3** | **-156.64** | **339.28** | **349.68** | **0.02** | **0.23** | **0.00** | **0.48** | **NA** | **NA** | **NA** | **NA** |
|  | MuHiSSE1 | -157.49 | 342.99 | 355.34 | 0.09 | 0.09 | 0.39 | 0.39 | 0.00 | 0.00 | 0.49 | 0.49 |
|  | MuHiSSE2 | -160.26 | 348.51 | 360.87 | 0.16 | 0.00 | 0.16 | 0.00 | 0.00 | 0.50 | 0.00 | 0.50 |
|  | MuHiSSE3 | -162.18 | 376.37 | 440.19 | 0.12 | 0.00 | 0.00 | 0.00 | 0.18 | 0.61 | 0.00 | 0.37 |
|  | MuCID2 | -169.12 | 362.23 | 370.90 | 0.39 | 0.39 | 0.39 | 0.39 | 0.11 | 0.11 | 0.11 | 0.11 |
| **BEAST MCC plastid** | Constant rate | -172.92 | 365.84 | 371.63 | 0.15 | 0.15 | 0.15 | 0.15 | NA | NA | NA | NA |
|  | NoHidden1 | -159.16 | 340.32 | 347.45 | 0.11 | 0.71 | 0.11 | 0.71 | NA | NA | NA | NA |
|  | **NoHidden2** | **-158.12** | **338.24** | **345.38** | **0.08** | **0.08** | **0.64** | **0.64** | **NA** | **NA** | **NA** | **NA** |
|  | NoHidden3 | -159.13 | 344.26 | 354.66 | 0.11 | 0.67 | 0.00 | 0.78 | NA | NA | NA | NA |
|  | MuHiSSE1 | -157.38 | 342.76 | 355.12 | 0.06 | 0.06 | 0.90 | 0.90 | 0.07 | 0.07 | 0.38 | 0.38 |
|  | MuHiSSE2 | -155.42 | 338.85 | 351.20 | 0.13 | 0.00 | 0.13 | 0.00 | 0.02 | 0.74 | 0.02 | 0.74 |
|  | MuHiSSE3 | -153.80 | 359.60 | 423.41 | 0.14 | 0.00 | 0.00 | 0.66 | 0.03 | 0.70 | 0.00 | 0.82 |
|  | MuCID2 | -161.43 | 346.86 | 355.53 | 0.62 | 0.62 | 0.62 | 0.62 | 0.11 | 0.11 | 0.11 | 0.11 |
| **BEAST cluster 1** | Constant rate | -180.53 | 381.06 | 386.85 | 0.13 | 0.13 | 0.13 | 0.13 | NA | NA | NA | NA |
|  | NoHidden1 | -170.51 | 363.03 | 370.16 | 0.10 | 0.44 | 0.10 | 0.44 | NA | NA | NA | NA |
|  | **NoHidden2** | **-162.26** | **346.52** | **353.65** | **0.04** | **0.04** | **0.47** | **0.47** | **NA** | **NA** | **NA** | **NA** |
|  | NoHidden3 | -161.90 | 349.81 | 360.21 | 0.03 | 0.43 | 0.00 | 0.15 | NA | NA | NA | NA |
|  | MuHiSSE1 | -173.66 | 375.33 | 387.68 | 0.23 | 0.23 | 0.35 | 0.35 | 0.00 | 0.00 | 0.00 | 0.00 |
|  | MuHiSSE2 | -165.80 | 359.61 | 371.96 | 0.14 | 0.00 | 0.14 | 0.00 | 0.00 | 0.48 | 0.00 | 0.48 |
|  | MuHiSSE3 | -158.94 | 369.87 | 433.69 | 0.05 | 0.00 | 2.22 | 0.00 | 0.00 | 0.20 | 0.00 | 0.64 |
|  | MuCID2 | -174.04 | 372.08 | 380.75 | 0.10 | 0.10 | 0.10 | 0.10 | 0.39 | 0.39 | 0.39 | 0.39 |
| **BEAST cluster 2** | Constant rate | -167.28 | 354.56 | 360.35 | 0.17 | 0.17 | 0.17 | 0.17 | NA | NA | NA | NA |
|  | NoHidden1 | -157.99 | 337.99 | 345.12 | 0.13 | 0.63 | 0.13 | 0.63 | NA | NA | NA | NA |
|  | **NoHidden2** | **-156.90** | **335.79** | **342.93** | **0.13** | **0.13** | **0.98** | **0.98** | **NA** | **NA** | **NA** | **NA** |
|  | NoHidden3 | -155.57 | 337.13 | 347.53 | 0.13 | 0.00 | 1.07 | 0.89 | NA | NA | NA | NA |
|  | MuHiSSE1 | -153.97 | 335.93 | 348.28 | 0.19 | 0.19 | 1.15 | 1.15 | 0.00 | 0.00 | 0.00 | 0.00 |
|  | MuHiSSE2 | -154.34 | 336.68 | 349.04 | 0.00 | 0.00 | 0.00 | 0.00 | 0.19 | 0.72 | 0.19 | 0.72 |
|  | MuHiSSE3 | -153.18 | 358.35 | 422.17 | 0.00 | 0.00 | 0.00 | 0.00 | 0.20 | 0.00 | 0.00 | 1.16 |
|  | MuCID2 | -162.88 | 349.76 | 358.42 | 0.12 | 0.12 | 0.12 | 0.12 | 0.44 | 0.44 | 0.44 | 0.44 |
| **BEAST cluster 3** | Constant rate | -175.31 | 370.62 | 376.41 | 0.14 | 0.14 | 0.14 | 0.14 | NA | NA | NA | NA |
|  | NoHidden1 | -164.13 | 350.26 | 357.40 | 0.10 | 0.55 | 0.10 | 0.55 | NA | NA | NA | NA |
|  | **NoHidden2** | **-156.04** | **334.07** | **341.21** | **0.07** | **0.07** | **0.89** | **0.89** | **NA** | **NA** | **NA** | **NA** |
|  | NoHidden3 | -156.24 | 338.48 | 348.88 | 0.04 | 0.15 | 0.54 | 0.60 | NA | NA | NA | NA |
|  | MuHiSSE1 | -164.16 | 356.32 | 368.68 | 6.21 | 6.21 | 0.67 | 0.67 | 0.09 | 0.09 | 0.00 | 0.00 |
|  | MuHiSSE2 | -159.38 | 346.76 | 359.12 | 0.15 | 0.00 | 0.15 | 0.00 | 0.00 | 0.62 | 0.00 | 0.62 |
|  | MuHiSSE3 | -156.20 | 364.40 | 428.22 | 0.07 | 0.27 | 0.00 | 0.60 | 0.00 | 0.00 | 0.00 | 0.00 |
|  | MuCID2 | -167.20 | 358.40 | 367.06 | 0.10 | 0.10 | 0.10 | 0.10 | 0.49 | 0.49 | 0.49 | 0.49 |
| **BEAST cluster 4** | Constant rate | -172.73 | 365.47 | 371.26 | 0.15 | 0.15 | 0.15 | 0.15 | NA | NA | NA | NA |
|  | NoHidden1 | -160.56 | 343.13 | 350.26 | 0.11 | 0.61 | 0.11 | 0.61 | NA | NA | NA | NA |
|  | **NoHidden2** | **-153.68** | **329.35** | **336.49** | **0.03** | **0.03** | **0.59** | **0.59** | **NA** | **NA** | **NA** | **NA** |
|  | NoHidden3 | -150.94 | 327.87 | 338.27 | 0.02 | 0.66 | 0.00 | 0.32 | NA | NA | NA | NA |
|  | MuHiSSE1 | -158.06 | 344.11 | 356.47 | 5.22 | 5.22 | 0.00 | 0.00 | 0.00 | 0.00 | 0.00 | 0.00 |
|  | MuHiSSE2 | -154.85 | 337.69 | 350.04 | 0.16 | 0.00 | 0.16 | 0.00 | 0.00 | 0.67 | 0.00 | 0.67 |
|  | MuHiSSE3 | -151.71 | 355.42 | 419.24 | 0.17 | 0.00 | 0.00 | 0.00 | 0.00 | 0.91 | 0.01 | 0.01 |
|  | MuCID2 | -163.13 | 350.26 | 358.92 | 0.55 | 0.55 | 0.55 | 0.55 | 0.11 | 0.11 | 0.11 | 0.11 |
| **BEAST cluster 5** | Constant rate | -179.21 | 378.42 | 384.21 | 0.13 | 0.13 | 0.13 | 0.13 | NA | NA | NA | NA |
|  | NoHidden1 | -169.91 | 361.82 | 368.95 | 0.10 | 0.50 | 0.10 | 0.50 | NA | NA | NA | NA |
|  | NoHidden2 | -166.57 | 355.14 | 362.27 | 0.10 | 0.10 | 0.74 | 0.74 | NA | NA | NA | NA |
|  | **NoHidden3** | **-165.30** | **356.59** | **366.99** | **0.11** | **0.00** | **3.50** | **0.75** | **NA** | **NA** | **NA** | **NA** |
|  | MuHiSSE1 | -163.75 | 355.51 | 367.86 | 0.15 | 0.15 | 0.81 | 0.81 | 0.00 | 0.00 | 0.00 | 0.00 |
|  | MuHiSSE2 | -165.82 | 359.64 | 371.99 | 0.00 | 0.00 | 0.00 | 0.00 | 0.15 | 0.58 | 0.15 | 0.58 |
|  | MuHiSSE3 | -163.91 | 379.82 | 443.63 | 0.16 | 0.00 | 0.00 | 0.88 | 0.00 | 0.00 | 0.00 | 0.00 |
|  | MuCID2 | -173.35 | 370.71 | 379.38 | 0.38 | 0.38 | 0.38 | 0.38 | 0.10 | 0.10 | 0.10 | 0.10 |
| **0.BEAST cluster 6** | Constant rate | -175.09 | 370.17 | 375.96 | 0.14 | 0.14 | 0.14 | 0.14 | NA | NA | NA | NA |
|  | NoHidden1 | -167.18 | 356.36 | 363.50 | 0.11 | 0.47 | 0.11 | 0.47 | NA | NA | NA | NA |
|  | **NoHidden2** | **-166.76** | **355.51** | **362.65** | **0.11** | **0.11** | **0.82** | **0.82** | **NA** | **NA** | **NA** | **NA** |
|  | NoHidden3 | -165.58 | 357.16 | 367.56 | 0.11 | 0.62 | 0.00 | 0.00 | NA | NA | NA | NA |
|  | MuHiSSE1 | -171.52 | 371.04 | 383.39 | 0.20 | 0.20 | 0.24 | 0.24 | 0.00 | 0.00 | 0.13 | 0.13 |
|  | MuHiSSE2 | -163.53 | 355.06 | 367.41 | 0.00 | 0.00 | 0.00 | 0.00 | 0.15 | 0.54 | 0.15 | 0.54 |
|  | MuHiSSE3 | -161.76 | 375.53 | 439.34 | 0.11 | 0.00 | 0.00 | 0.00 | 0.13 | 0.70 | 0.00 | 0.00 |
|  | MuCID2 | -171.27 | 366.54 | 375.20 | 0.36 | 0.36 | 0.36 | 0.36 | 0.11 | 0.11 | 0.11 | 0.11 |
| **BEAST cluster 7** | Constant rate | -177.01 | 374.01 | 379.80 | 0.14 | 0.14 | 0.14 | 0.14 | NA | NA | NA | NA |
|  | NoHidden1 | -164.26 | 350.51 | 357.65 | 0.10 | 0.58 | 0.10 | 0.58 | NA | NA | NA | NA |
|  | **NoHidden2** | -156.42 | 334.85 | 341.98 | 0.07 | 0.07 | 2.47 | 2.47 | **NA** | **NA** | **NA** | **NA** |
|  | NoHidden3 | -153.29 | 332.58 | 342.98 | 0.03 | 0.22 | 0.01 | 0.73 | NA | NA | NA | NA |
|  | MuHiSSE1 | -164.32 | 356.64 | 368.99 | 3.20 | 3.20 | 0.88 | 0.88 | 0.00 | 0.00 | 0.00 | -164.32 |
|  | MuHiSSE2 | -159.08 | 346.17 | 358.52 | 0.15 | 0.00 | 0.15 | 0.00 | 0.00 | 0.62 | 0.00 | -159.08 |
|  | MuHiSSE3 | -153.29 | 358.59 | 422.40 | 0.12 | 0.02 | 0.43 | 1.19 | 0.00 | 0.00 | 0.00 | 0.28 |
|  | MuCID2 | -168.07 | 360.13 | 368.80 | 0.52 | 0.52 | 0.52 | 0.52 | 0.10 | 0.10 | 0.10 | 0.10 |
| **BEAST cluster 8** | Constant rate | -171.44 | 362.88 | 368.67 | 0.16 | 0.16 | 0.16 | 0.16 | NA | NA | NA | NA |
|  | NoHidden1 | -160.31 | 342.62 | 349.75 | 0.12 | 0.56 | 0.12 | 0.56 | NA | NA | NA | NA |
|  | **NoHidden2** | **-151.73** | **325.45** | **332.59** | **0.07** | **0.07** | **0.63** | **0.63** | **NA** | **NA** | **NA** | **NA** |
|  | NoHidden3 | -158.76 | 343.51 | 353.91 | 0.12 | 0.86 | 0.00 | 0.00 | NA | NA | NA | NA |
|  | MuHiSSE1 | -150.45 | 328.90 | 341.25 | 0.13 | 0.13 | 0.00 | 0.00 | 0.00 | 0.66 | 0.00 | 0.66 |
|  | MuHiSSE2 | -155.19 | 338.39 | 350.74 | 0.19 | 0.00 | 0.19 | 0.00 | 0.00 | 0.00 | 1.06 | 1.06 |
|  | MuHiSSE3 | -150.33 | 352.65 | 416.47 | 0.20 | 0.00 | 0.00 | 0.00 | 0.00 | 1.98 | 2.28 | 0.00 |
|  | MuCID2 | -163.94 | 351.87 | 360.54 | 0.12 | 0.12 | 0.12 | 0.12 | 0.53 | 0.53 | 0.53 | 0.53 |
| **BEAST cluster 9** | Constant rate | -173.93 | 367.86 | 373.65 | 0.15 | 0.15 | 0.15 | 0.15 | NA | NA | NA | NA |
|  | NoHidden1 | -160.87 | 343.74 | 350.87 | 0.11 | 0.63 | 0.11 | 0.63 | NA | NA | NA | NA |
|  | **NoHidden2** | -151.93 | 325.85 | 332.99 | 0.06 | 0.06 | 1.07 | 1.07 | **NA** | **NA** | **NA** | **NA** |
|  | NoHidden3 | -149.27 | 324.54 | 334.94 | 0.05 | 0.00 | 0.13 | 0.77 | NA | NA | NA | NA |
|  | MuHiSSE1 | -151.86 | 331.72 | 344.07 | 0.09 | 0.09 | 0.75 | 0.75 | 0.00 | 0.00 | 0.00 | 0.00 |
|  | MuHiSSE2 | -154.74 | 337.47 | 349.82 | 0.15 | 0.00 | 0.15 | 0.00 | 0.00 | 0.70 | 0.00 | 0.70 |
|  | MuHiSSE3 | -147.91 | 347.82 | 411.64 | 0.04 | 0.31 | 0.00 | 2.07 | 0.00 | 0.00 | 0.00 | 0.94 |
|  | MuCID2 | -164.04 | 352.08 | 360.75 | 0.56 | 0.56 | 0.56 | 0.56 | 0.11 | 0.11 | 0.11 | 0.11 |
| **BEAST cluster 10** | Constant rate | -176.29 | 372.58 | 378.37 | 0.14 | 0.14 | 0.14 | 0.14 | NA | NA | NA | NA |
|  | NoHidden1 | -164.74 | 351.48 | 358.61 | 0.10 | 0.53 | 0.10 | 0.53 | NA | NA | NA | NA |
|  | **NoHidden2** | -155.95 | 333.90 | 341.04 | 0.04 | 0.04 | 0.58 | 0.58 | **NA** | **NA** | **NA** | **NA** |
|  | NoHidden3 | -153.03 | 332.07 | 342.47 | 0.03 | 0.21 | 0.02 | 0.66 | NA | NA | NA | NA |
|  | MuHiSSE1 | -159.38 | 346.76 | 359.11 | 0.00 | 0.00 | 0.00 | 0.00 | 0.17 | 0.17 | 0.99 | 0.99 |
|  | MuHiSSE2 | -159.74 | 347.49 | 359.84 | 0.15 | 0.00 | 0.15 | 0.00 | 0.00 | 0.57 | 0.00 | 0.57 |
|  | MuHiSSE3 | -156.35 | 364.69 | 428.51 | 0.18 | 0.00 | 0.00 | 1.02 | 0.00 | 0.00 | 0.00 | 0.37 |
|  | MuCID2 | -168.61 | 361.22 | 369.89 | 0.10 | 0.10 | 0.10 | 0.10 | 0.48 | 0.48 | 0.48 | 0.48 |
