## Supplementary material for "‘Dispersification’ of *Agalinis* (Orobanchaceae) into South America is associated with hummingbird pollination and perennial life history shifts": Table S8.

**Table S8.** Log-likelihood values (lnL), Akaike Information Criterion (AIC), and corrected AIC (AICc) for Multistate Hidden State Speciation and Extinction (MuHiSSE) models combining pollination syndrome and life history strategy with diversification in Agalinis for the 100 randomly sampled post–burn-in MCMC trees. The constant rate and MuCID2 models assume diversification is independent of these traits. Models NoHidden1, NoHidden2, and NoHidden3 do not include hidden states, while MuHiSSE1, MuHiSSE2, and MuHiSSE3 include hidden states A and B. Model NoHidden1 associates diversification with life history strategy, NoHidden2 associates diversification with pollination syndrome, and NoHidden3 assigns unique turnover rates to each category. The corresponding MuHiSSE models include hidden states for each of these trait associations.

| Tree | Model | lnL | AIC | AICc |
|---|---|---|---|---|
| Tree1 | Constant rate | -179.29 | 378.58 | 384.37 |
| | NoHidden1 | -167.07 | 356.14 | 363.27 |
| | **NoHidden2** | **-159.18** | **340.37** | **347.50** |
| | NoHidden3 | -166.00 | 357.99 | 368.39 |
| | MuHiSSE1 | -161.88 | 351.77 | 364.12 |
| | MuHiSSE2 | -162.33 | 352.65 | 365.01 |
| | MuHiSSE3 | -159.94 | 371.87 | 435.69 |
| | MuCID2 | -170.57 | 365.14 | 373.80 |
| Tree2 | Constant rate | -174.76 | 369.52 | 375.31 |
| | NoHidden1 | -164.97 | 351.95 | 359.08 |
| | **NoHidden2** | **-155.30** | **332.59** | **339.73** |
| | NoHidden3 | -163.33 | 352.65 | 363.05 |
| | MuHiSSE1 | -158.99 | 345.98 | 358.33 |
| | MuHiSSE2 | -158.46 | 344.91 | 357.26 |
| | MuHiSSE3 | -154.22 | 360.43 | 424.25 |
| | MuCID2 | -167.98 | 359.96 | 368.63 |
| Tree3 | Constant rate | -176.95 | 373.91 | 379.70 |
| | NoHidden1 | -163.99 | 349.97 | 357.11 |
| | **NoHidden2** | **-155.87** | **333.75** | **340.88** |
| | NoHidden3 | -163.59 | 353.18 | 363.58 |
| | MuHiSSE1 | -155.17 | 338.33 | 350.69 |
| | MuHiSSE2 | -158.62 | 345.24 | 357.59 |
| | MuHiSSE3 | -152.51 | 357.01 | 420.83 |
| | MuCID2 | -166.91 | 357.81 | 366.48 |
| Tree4 | Constant rate | -178.87 | 377.75 | 383.54 |
| | NoHidden1 | -167.22 | 356.44 | 363.58 |
| | **NoHidden2** | **-158.98** | **339.97** | **347.10** |
| | NoHidden3 | -156.22 | 338.45 | 348.85 |
| | MuHiSSE1 | -162.37 | 352.73 | 365.08 |
| | MuHiSSE2 | -162.16 | 352.32 | 364.68 |
| | MuHiSSE3 | -156.16 | 364.32 | 428.14 |
| | MuCID2 | -170.97 | 365.93 | 374.60 |
| Tree5 | Constant rate | -167.78 | 355.56 | 361.35 |
| | NoHidden1 | -155.24 | 332.49 | 339.62 |
| | **NoHidden2** | **-153.38** | **328.77** | **335.91** |
| | NoHidden3 | -153.88 | 333.75 | 344.15 |
| | MuHiSSE1 | -158.84 | 345.68 | 358.03 |
| | MuHiSSE2 | -151.92 | 331.85 | 344.20 |
| | MuHiSSE3 | -149.38 | 350.76 | 414.58 |
| | MuCID2 | -158.28 | 340.55 | 349.22 |
| Tree6 | Constant rate | -184.06 | 388.12 | 393.91 |
| | NoHidden1 | -174.07 | 370.15 | 377.28 |
| | **NoHidden2** | **-165.19** | **352.39** | **359.52** |
| | NoHidden3 | -163.18 | 352.36 | 362.76 |

| | | | | |
|---|---|---|---|---|
| Tree6 | MuHiSSE1 | -164.60 | 357.21 | 369.56 |
| | MuHiSSE2 | -167.92 | 363.84 | 376.20 |
| | MuHiSSE3 | -160.95 | 373.90 | 437.72 |
| | MuCID2 | -174.72 | 373.44 | 382.11 |
| Tree7 | Constant rate | -177.90 | 375.79 | 381.58 |
| | NoHidden1 | -164.92 | 351.83 | 358.97 |
| | **NoHidden2** | **-153.35** | **328.70** | **335.83** |
| | NoHidden3 | -150.20 | 326.40 | 336.80 |
| | MuHiSSE1 | -153.19 | 334.38 | 346.73 |
| | MuHiSSE2 | -158.69 | 345.37 | 357.73 |
| | MuHiSSE3 | -149.56 | 351.11 | 414.93 |
| | MuCID2 | -167.84 | 359.68 | 368.35 |
| Tree8 | Constant rate | -182.76 | 385.51 | 391.30 |
| | NoHidden1 | -171.27 | 364.54 | 371.67 |
| | **NoHidden2** | **-162.85** | **347.69** | **354.83** |
| | NoHidden3 | -162.79 | 351.57 | 361.97 |
| | MuHiSSE1 | -167.63 | 363.26 | 375.62 |
| | MuHiSSE2 | -166.73 | 361.45 | 373.80 |
| | MuHiSSE3 | -162.20 | 376.39 | 440.21 |
| | MuCID2 | -174.93 | 373.87 | 382.53 |
| Tree9 | Constant rate | -172.64 | 365.27 | 371.06 |
| | NoHidden1 | -160.81 | 343.63 | 350.76 |
| | **NoHidden2** | **-156.36** | **334.73** | **341.86** |
| | NoHidden3 | -160.28 | 346.55 | 356.95 |
| | MuHiSSE1 | -155.96 | 339.92 | 352.28 |
| | MuHiSSE2 | -156.01 | 340.01 | 352.37 |
| | MuHiSSE3 | -154.58 | 361.16 | 424.98 |
| | MuCID2 | -164.03 | 352.07 | 360.74 |
| Tree10 | Constant rate | -180.85 | 381.70 | 387.49 |
| | NoHidden1 | -168.60 | 359.21 | 366.34 |
| | NoHidden2 | -160.54 | 343.07 | 350.21 |
| | **NoHidden3** | **-156.48** | **338.95** | **349.35** |
| | MuHiSSE1 | -160.40 | 348.81 | 361.16 |
| | MuHiSSE2 | -163.53 | 355.05 | 367.41 |
| | MuHiSSE3 | -157.09 | 366.19 | 430.01 |
| | MuCID2 | -171.96 | 367.92 | 376.59 |
| Tree11 | Constant rate | -178.21 | 376.42 | 382.21 |
| | NoHidden1 | -165.79 | 353.58 | 360.72 |
| | **NoHidden2** | **-156.76** | **335.52** | **342.66** |
| | NoHidden3 | -165.23 | 356.45 | 366.85 |
| | MuHiSSE1 | -156.90 | 341.80 | 354.15 |
| | MuHiSSE2 | -160.38 | 348.77 | 361.12 |
| | MuHiSSE3 | -157.49 | 366.98 | 430.80 |
| | MuCID2 | -168.61 | 361.23 | 369.89 |
| Tree12 | Constant rate | -179.92 | 379.83 | 385.62 |
| | NoHidden1 | -167.27 | 356.55 | 363.68 |
| | NoHidden2 | -159.12 | 340.23 | 347.37 |
| | **NoHidden3** | **-155.03** | **336.06** | **346.46** |
| | MuHiSSE1 | -172.00 | 372.01 | 384.36 |
| | MuHiSSE2 | -162.40 | 352.81 | 365.16 |
| | MuHiSSE3 | -157.71 | 367.42 | 431.24 |
| | MuCID2 | -170.97 | 365.94 | 374.61 |
| | Constant rate | -178.90 | 377.79 | 383.58 |

| | | | | |
|---|---|---|---|---|
| Tree13 | NoHidden1 | -168.06 | 358.12 | 365.25 |
| | **NoHidden2** | **-159.84** | **341.68** | **348.82** |
| | NoHidden3 | -168.43 | 362.85 | 373.25 |
| | MuHiSSE1 | -171.16 | 370.31 | 382.67 |
| | MuHiSSE2 | -163.13 | 354.26 | 366.61 |
| | MuHiSSE3 | -156.29 | 364.58 | 428.40 |
| | MuCID2 | -171.72 | 367.45 | 376.11 |
| Tree14 | Constant rate | -176.88 | 373.76 | 379.55 |
| | NoHidden1 | -165.72 | 353.43 | 360.57 |
| | **NoHidden2** | **-159.29** | **340.57** | **347.71** |
| | NoHidden3 | -158.27 | 342.53 | 352.93 |
| | MuHiSSE1 | -166.60 | 361.21 | 373.56 |
| | MuHiSSE2 | -160.23 | 348.46 | 360.82 |
| | MuHiSSE3 | -155.51 | 363.03 | 426.84 |
| | MuCID2 | -168.91 | 361.81 | 370.48 |
| Tree15 | Constant rate | -178.03 | 376.07 | 381.86 |
| | NoHidden1 | -165.77 | 353.53 | 360.67 |
| | **NoHidden2** | **-153.36** | **328.71** | **335.85** |
| | NoHidden3 | -150.85 | 327.70 | 338.10 |
| | MuHiSSE1 | -163.18 | 354.36 | 366.71 |
| | MuHiSSE2 | -159.17 | 346.35 | 358.70 |
| | MuHiSSE3 | -151.84 | 355.69 | 419.51 |
| | MuCID2 | -169.33 | 362.66 | 371.33 |
| Tree16 | Constant rate | -178.93 | 377.86 | 383.65 |
| | NoHidden1 | -168.04 | 358.07 | 365.21 |
| | **NoHidden2** | **-160.21** | **342.41** | **349.55** |
| | NoHidden3 | -166.34 | 358.67 | 369.07 |
| | MuHiSSE1 | -171.99 | 371.98 | 384.33 |
| | MuHiSSE2 | -163.29 | 354.57 | 366.92 |
| | MuHiSSE3 | -160.33 | 372.66 | 436.48 |
| | MuCID2 | -171.33 | 366.66 | 375.33 |
| Tree17 | Constant rate | -174.52 | 369.03 | 374.82 |
| | NoHidden1 | -164.10 | 350.20 | 357.33 |
| | **NoHidden2** | **-155.38** | **332.77** | **339.90** |
| | NoHidden3 | -153.66 | 333.31 | 343.71 |
| | MuHiSSE1 | -166.65 | 361.30 | 373.65 |
| | MuHiSSE2 | -159.17 | 346.35 | 358.70 |
| | MuHiSSE3 | -154.10 | 360.20 | 424.02 |
| | MuCID2 | -165.85 | 355.70 | 364.37 |
| Tree18 | Constant rate | -170.88 | 361.77 | 367.56 |
| | NoHidden1 | -160.32 | 342.63 | 349.77 |
| | NoHidden2 | -153.57 | 329.13 | 336.27 |
| | **NoHidden3** | **-148.67** | **323.33** | **333.73** |
| | MuHiSSE1 | -156.41 | 340.81 | 353.17 |
| | MuHiSSE2 | -155.28 | 338.55 | 350.90 |
| | MuHiSSE3 | -148.15 | 348.30 | 412.12 |
| | MuCID2 | -163.68 | 351.35 | 360.02 |
| Tree19 | Constant rate | -178.24 | 376.48 | 382.27 |
| | NoHidden1 | -166.09 | 354.18 | 361.32 |
| | **NoHidden2** | **-163.57** | **349.13** | **356.27** |
| | NoHidden3 | -164.79 | 355.59 | 365.99 |
| | MuHiSSE1 | -169.79 | 367.58 | 379.93 |
| | MuHiSSE2 | -160.61 | 349.23 | 361.58 |

| | | | | |
|---|---|---|---|---|
| | MuHiSSE3 | -152.95 | 357.90 | 421.72 |
| | MuCID2 | -166.38 | 356.77 | 365.44 |
| Tree20 | Constant rate | -176.87 | 373.75 | 379.54 |
| | NoHidden1 | -166.46 | 354.92 | 362.05 |
| | **NoHidden2** | **-158.54** | **339.08** | **346.22** |
| | NoHidden3 | -159.65 | 345.29 | 355.69 |
| | MuHiSSE1 | -159.40 | 346.81 | 359.16 |
| | MuHiSSE2 | -162.48 | 352.96 | 365.32 |
| | MuHiSSE3 | -157.85 | 367.70 | 431.52 |
| | MuCID2 | -170.15 | 364.29 | 372.96 |
| Tree21 | Constant rate | -175.07 | 370.14 | 375.93 |
| | NoHidden1 | -166.59 | 355.18 | 362.32 |
| | **NoHidden2** | **-166.04** | **354.08** | **361.22** |
| | NoHidden3 | -166.31 | 358.62 | 369.02 |
| | MuHiSSE1 | -160.71 | 349.42 | 361.78 |
| | MuHiSSE2 | -161.17 | 350.33 | 362.68 |
| | MuHiSSE3 | -162.62 | 377.24 | 441.05 |
| | MuCID2 | -170.34 | 364.68 | 373.35 |
| Tree22 | Constant rate | -179.63 | 379.26 | 385.05 |
| | NoHidden1 | -165.26 | 352.52 | 359.65 |
| | **NoHidden2** | **-157.38** | **336.75** | **343.89** |
| | NoHidden3 | -165.15 | 356.29 | 366.69 |
| | MuHiSSE1 | -160.27 | 348.53 | 360.89 |
| | MuHiSSE2 | -158.29 | 344.57 | 356.93 |
| | MuHiSSE3 | -155.21 | 362.42 | 426.24 |
| | MuCID2 | -168.15 | 360.30 | 368.97 |
| Tree23 | Constant rate | -177.62 | 375.25 | 381.04 |
| | NoHidden1 | -165.75 | 353.50 | 360.64 |
| | **NoHidden2** | **-157.41** | **336.81** | **343.95** |
| | NoHidden3 | -164.87 | 355.74 | 366.14 |
| | MuHiSSE1 | -166.97 | 361.95 | 374.30 |
| | MuHiSSE2 | -161.47 | 350.94 | 363.30 |
| | MuHiSSE3 | -155.40 | 362.80 | 426.62 |
| | MuCID2 | -169.18 | 362.36 | 371.02 |
| Tree24 | Constant rate | -177.02 | 374.04 | 379.83 |
| | NoHidden1 | -168.16 | 358.31 | 365.45 |
| | **NoHidden2** | **-166.65** | **355.31** | **362.44** |
| | NoHidden3 | -167.97 | 361.93 | 372.33 |
| | MuHiSSE1 | -171.56 | 371.13 | 383.48 |
| | MuHiSSE2 | -164.27 | 356.54 | 368.90 |
| | MuHiSSE3 | -161.46 | 374.92 | 438.73 |
| | MuCID2 | -171.97 | 367.94 | 376.61 |
| Tree25 | Constant rate | -175.15 | 370.30 | 376.09 |
| | NoHidden1 | -160.60 | 343.20 | 350.33 |
| | **NoHidden2** | **-151.94** | **325.89** | **333.02** |
| | NoHidden3 | -148.88 | 323.76 | 334.16 |
| | MuHiSSE1 | -164.85 | 357.70 | 370.05 |
| | MuHiSSE2 | -155.72 | 339.44 | 351.80 |
| | MuHiSSE3 | -151.45 | 354.91 | 418.73 |
| | MuCID2 | -163.57 | 351.13 | 359.80 |
| | Constant rate | -175.62 | 371.25 | 377.03 |
| | NoHidden1 | -165.04 | 352.09 | 359.22 |
| | **NoHidden2** | **-162.32** | **346.63** | **353.77** |

| | | | | |
|---|---|---|---|---|
| Tree26 | NoHidden3 | -164.69 | 355.38 | 365.78 |
| | MuHiSSE1 | -158.39 | 344.79 | 357.14 |
| | MuHiSSE2 | -160.29 | 348.58 | 360.93 |
| | MuHiSSE3 | -159.61 | 371.21 | 435.03 |
| | MuCID2 | -166.22 | 356.44 | 365.11 |
| Tree27 | Constant rate | -176.29 | 372.57 | 378.36 |
| | NoHidden1 | -165.54 | 353.09 | 360.22 |
| | NoHidden2 | -157.95 | 337.89 | 345.03 |
| | **NoHidden3** | **-153.97** | **333.94** | **344.34** |
| | MuHiSSE1 | -158.10 | 344.20 | 356.55 |
| | MuHiSSE2 | -159.93 | 347.85 | 360.20 |
| | MuHiSSE3 | -153.49 | 358.98 | 422.79 |
| | MuCID2 | -169.34 | 362.69 | 371.36 |
| Tree28 | Constant rate | -175.59 | 371.17 | 376.96 |
| | NoHidden1 | -158.27 | 338.54 | 345.68 |
| | **NoHidden2** | **-156.81** | **335.62** | **342.75** |
| | NoHidden3 | -163.01 | 352.01 | 362.41 |
| | MuHiSSE1 | -168.12 | 364.24 | 376.60 |
| | MuHiSSE2 | -158.46 | 344.91 | 357.27 |
| | MuHiSSE3 | -156.66 | 365.33 | 429.15 |
| | MuCID2 | -167.83 | 359.66 | 368.33 |
| Tree29 | Constant rate | -176.89 | 373.79 | 379.58 |
| | NoHidden1 | -169.47 | 360.93 | 368.07 |
| | **NoHidden2** | **-162.23** | **346.46** | **353.60** |
| | NoHidden3 | -169.09 | 364.18 | 374.58 |
| | MuHiSSE1 | -164.37 | 356.74 | 369.10 |
| | MuHiSSE2 | -165.26 | 358.53 | 370.88 |
| | MuHiSSE3 | -165.74 | 383.48 | 447.30 |
| | MuCID2 | -172.97 | 369.93 | 378.60 |
| Tree30 | Constant rate | -181.38 | 382.75 | 388.54 |
| | NoHidden1 | -172.68 | 367.36 | 374.49 |
| | **NoHidden2** | **-171.23** | **364.45** | **371.59** |
| | NoHidden3 | -172.69 | 371.37 | 381.77 |
| | MuHiSSE1 | -162.78 | 353.56 | 365.91 |
| | MuHiSSE2 | -167.13 | 362.25 | 374.61 |
| | MuHiSSE3 | -169.08 | 390.15 | 453.97 |
| | MuCID2 | -176.77 | 377.55 | 386.21 |
| Tree31 | Constant rate | -186.37 | 392.74 | 398.53 |
| | NoHidden1 | -175.16 | 372.32 | 379.46 |
| | **NoHidden2** | **-163.71** | **349.42** | **356.55** |
| | NoHidden3 | -174.55 | 375.10 | 385.50 |
| | MuHiSSE1 | -163.56 | 355.11 | 367.47 |
| | MuHiSSE2 | -165.94 | 359.88 | 372.23 |
| | MuHiSSE3 | -166.43 | 384.86 | 448.68 |
| | MuCID2 | -178.57 | 381.14 | 389.81 |
| Tree32 | Constant rate | -180.81 | 381.63 | 387.41 |
| | NoHidden1 | -170.05 | 362.11 | 369.24 |
| | NoHidden2 | -167.85 | 357.71 | 364.84 |
| | **NoHidden3** | **-162.61** | **351.23** | **361.63** |
| | MuHiSSE1 | -164.07 | 356.13 | 368.48 |
| | MuHiSSE2 | -164.80 | 357.60 | 369.95 |
| | MuHiSSE3 | -158.86 | 369.71 | 433.53 |
| | MuCID2 | -173.93 | 371.85 | 380.52 |

| | | | | |
|---|---|---|---|---|
| Tree33 | Constant rate | -181.12 | 382.23 | 388.02 |
| | **NoHidden1** | **-170.46** | **362.93** | **370.06** |
| | NoHidden2 | -170.80 | 363.59 | 370.73 |
| | NoHidden3 | -170.46 | 366.93 | 377.33 |
| | MuHiSSE1 | -159.83 | 347.66 | 360.01 |
| | MuHiSSE2 | -164.00 | 355.99 | 368.35 |
| | MuHiSSE3 | -163.45 | 378.90 | 442.72 |
| | MuCID2 | -173.54 | 371.08 | 379.75 |
| Tree34 | Constant rate | -173.91 | 367.82 | 373.61 |
| | NoHidden1 | -158.46 | 338.93 | 346.06 |
| | NoHidden2 | -153.20 | 328.41 | 335.54 |
| | **NoHidden3** | **-146.49** | **318.98** | **329.38** |
| | MuHiSSE1 | -151.38 | 330.75 | 343.11 |
| | MuHiSSE2 | -152.53 | 333.06 | 345.42 |
| | MuHiSSE3 | -145.62 | 343.25 | 407.07 |
| | MuCID2 | -162.41 | 348.81 | 357.48 |
| Tree35 | Constant rate | -177.65 | 375.31 | 381.10 |
| | NoHidden1 | -163.33 | 348.66 | 355.79 |
| | **NoHidden2** | **-157.72** | **337.43** | **344.57** |
| | NoHidden3 | -163.09 | 352.18 | 362.58 |
| | MuHiSSE1 | -157.15 | 342.29 | 354.65 |
| | MuHiSSE2 | -158.48 | 344.96 | 357.31 |
| | MuHiSSE3 | -154.98 | 361.95 | 425.77 |
| | MuCID2 | -167.18 | 358.37 | 367.04 |
| Tree36 | Constant rate | -176.32 | 372.64 | 378.43 |
| | NoHidden1 | -164.25 | 350.51 | 357.64 |
| | **NoHidden2** | **-157.22** | **336.45** | **343.58** |
| | NoHidden3 | -156.88 | 339.75 | 350.15 |
| | MuHiSSE1 | -156.92 | 341.84 | 354.19 |
| | MuHiSSE2 | -159.43 | 346.86 | 359.21 |
| | MuHiSSE3 | -156.26 | 364.52 | 428.34 |
| | MuCID2 | -167.68 | 359.37 | 368.03 |
| Tree37 | Constant rate | -179.83 | 379.66 | 385.45 |
| | NoHidden1 | -168.14 | 358.27 | 365.41 |
| | **NoHidden2** | **-158.18** | **338.37** | **345.50** |
| | NoHidden3 | -157.13 | 340.25 | 350.65 |
| | MuHiSSE1 | -155.85 | 339.70 | 352.06 |
| | MuHiSSE2 | -162.07 | 352.14 | 364.49 |
| | MuHiSSE3 | -158.91 | 369.82 | 433.63 |
| | MuCID2 | -172.03 | 368.06 | 376.73 |
| Tree38 | Constant rate | -176.51 | 373.01 | 378.80 |
| | NoHidden1 | -165.83 | 353.66 | 360.80 |
| | **NoHidden2** | **-158.97** | **339.93** | **347.07** |
| | NoHidden3 | -165.11 | 356.22 | 366.62 |
| | MuHiSSE1 | -160.90 | 349.80 | 362.15 |
| | MuHiSSE2 | -159.81 | 347.61 | 359.97 |
| | MuHiSSE3 | -158.48 | 368.95 | 432.77 |
| | MuCID2 | -167.28 | 358.56 | 367.22 |
| Tree39 | Constant rate | -177.08 | 374.17 | 379.96 |
| | NoHidden1 | -165.93 | 353.85 | 360.99 |
| | **NoHidden2** | **-160.14** | **342.29** | **349.42** |
| | NoHidden3 | -165.92 | 357.84 | 368.24 |
| | MuHiSSE1 | -159.86 | 347.71 | 360.07 |

| | | | | |
|---|---|---|---|---|
| | MuHiSSE2 | -161.33 | 350.66 | 363.01 |
| | MuHiSSE3 | -159.17 | 370.34 | 434.16 |
| | MuCID2 | -169.40 | 362.79 | 371.46 |
| Tree40 | Constant rate | -177.32 | 374.64 | 380.43 |
| | NoHidden1 | -164.24 | 350.48 | 357.62 |
| | **NoHidden2** | **-156.53** | **335.05** | **342.19** |
| | NoHidden3 | -155.27 | 336.54 | 346.94 |
| | MuHiSSE1 | -155.44 | 338.88 | 351.23 |
| | MuHiSSE2 | -158.95 | 345.90 | 358.25 |
| | MuHiSSE3 | -156.08 | 364.16 | 427.98 |
| | MuCID2 | -168.37 | 360.73 | 369.40 |
| Tree41 | Constant rate | -175.87 | 371.75 | 377.54 |
| | NoHidden1 | -163.69 | 349.38 | 356.52 |
| | **NoHidden2** | **-157.33** | **336.65** | **343.79** |
| | NoHidden3 | -162.11 | 350.23 | 360.63 |
| | MuHiSSE1 | -158.06 | 344.12 | 356.47 |
| | MuHiSSE2 | -157.51 | 343.02 | 355.37 |
| | MuHiSSE3 | -152.30 | 356.61 | 420.43 |
| | MuCID2 | -167.13 | 358.26 | 366.93 |
| Tree42 | Constant rate | -172.50 | 365.00 | 370.79 |
| | NoHidden1 | -159.84 | 341.69 | 348.82 |
| | **NoHidden2** | **-157.27** | **336.54** | **343.67** |
| | NoHidden3 | -157.02 | 340.03 | 350.43 |
| | MuHiSSE1 | -154.04 | 336.08 | 348.43 |
| | MuHiSSE2 | -156.17 | 340.34 | 352.69 |
| | MuHiSSE3 | -153.62 | 359.24 | 423.05 |
| | MuCID2 | -162.05 | 348.10 | 356.76 |
| Tree43 | Constant rate | -179.48 | 378.95 | 384.74 |
| | NoHidden1 | -165.98 | 353.96 | 361.09 |
| | **NoHidden2** | **-163.85** | **349.70** | **356.84** |
| | NoHidden3 | -165.00 | 356.01 | 366.41 |
| | MuHiSSE1 | -159.22 | 346.44 | 358.79 |
| | MuHiSSE2 | -161.35 | 350.70 | 363.05 |
| | MuHiSSE3 | -159.29 | 370.57 | 434.39 |
| | MuCID2 | -169.31 | 362.62 | 371.28 |
| Tree44 | Constant rate | -173.25 | 366.49 | 372.28 |
| | NoHidden1 | -163.58 | 349.17 | 356.31 |
| | **NoHidden2** | **-163.22** | **348.44** | **355.58** |
| | NoHidden3 | -162.73 | 351.46 | 361.86 |
| | MuHiSSE1 | -158.70 | 345.40 | 357.75 |
| | MuHiSSE2 | -161.32 | 350.64 | 362.99 |
| | MuHiSSE3 | -158.09 | 368.19 | 432.01 |
| | MuCID2 | -166.78 | 357.56 | 366.22 |
| Tree45 | Constant rate | -174.72 | 369.44 | 375.23 |
| | NoHidden1 | -166.90 | 355.81 | 362.94 |
| | **NoHidden2** | **-166.01** | **354.03** | **361.16** |
| | NoHidden3 | -166.42 | 358.84 | 369.24 |
| | MuHiSSE1 | -162.55 | 353.11 | 365.46 |
| | MuHiSSE2 | -162.66 | 353.33 | 365.68 |
| | MuHiSSE3 | -162.83 | 377.66 | 441.48 |
| | MuCID2 | -168.12 | 360.25 | 368.92 |
| | Constant rate | -174.33 | 368.67 | 374.46 |
| | NoHidden1 | -162.45 | 346.91 | 354.04 |

| | | | | |
|---|---|---|---|---|
| Tree46 | **NoHidden2** | **-156.57** | **335.14** | **342.27** |
| | NoHidden3 | -156.99 | 339.97 | 350.37 |
| | MuHiSSE1 | -156.23 | 340.46 | 352.81 |
| | MuHiSSE2 | -157.49 | 342.98 | 355.33 |
| | MuHiSSE3 | -156.02 | 364.03 | 427.85 |
| | MuCID2 | -164.54 | 353.08 | 361.75 |
| Tree47 | Constant rate | -176.43 | 372.85 | 378.64 |
| | NoHidden1 | -164.67 | 351.35 | 358.48 |
| | **NoHidden2** | **-157.41** | **336.81** | **343.95** |
| | NoHidden3 | -165.43 | 356.86 | 367.26 |
| | MuHiSSE1 | -160.19 | 348.39 | 360.74 |
| | MuHiSSE2 | -159.83 | 347.65 | 360.00 |
| | MuHiSSE3 | -159.23 | 370.46 | 434.28 |
| | MuCID2 | -167.41 | 358.82 | 367.48 |
| Tree48 | Constant rate | -174.40 | 368.80 | 374.59 |
| | NoHidden1 | -162.99 | 347.98 | 355.12 |
| | **NoHidden2** | **-157.30** | **336.59** | **343.73** |
| | NoHidden3 | -163.50 | 353.00 | 363.40 |
| | MuHiSSE1 | -167.59 | 363.18 | 375.53 |
| | MuHiSSE2 | -158.16 | 344.32 | 356.67 |
| | MuHiSSE3 | -158.34 | 368.69 | 432.50 |
| | MuCID2 | -167.08 | 358.17 | 366.83 |
| Tree49 | Constant rate | -181.64 | 383.28 | 389.07 |
| | NoHidden1 | -171.46 | 364.92 | 372.05 |
| | NoHidden2 | -163.58 | 349.16 | 356.30 |
| | **NoHidden3** | **-159.21** | **344.41** | **354.81** |
| | MuHiSSE1 | -162.98 | 353.96 | 366.31 |
| | MuHiSSE2 | -166.19 | 360.39 | 372.74 |
| | MuHiSSE3 | -160.42 | 372.84 | 436.65 |
| | MuCID2 | -174.28 | 372.56 | 381.23 |
| Tree50 | Constant rate | -175.73 | 371.46 | 377.25 |
| | NoHidden1 | -164.08 | 350.15 | 357.29 |
| | **NoHidden2** | **-156.73** | **335.46** | **342.60** |
| | NoHidden3 | -155.21 | 336.43 | 346.83 |
| | MuHiSSE1 | -156.64 | 341.27 | 353.63 |
| | MuHiSSE2 | -160.01 | 348.02 | 360.37 |
| | MuHiSSE3 | -156.65 | 365.29 | 429.11 |
| | MuCID2 | -167.59 | 359.18 | 367.85 |
| Tree51 | Constant rate | -178.33 | 376.66 | 382.45 |
| | NoHidden1 | -167.65 | 357.29 | 364.43 |
| | **NoHidden2** | **-167.57** | **357.13** | **364.27** |
| | NoHidden3 | -167.52 | 361.04 | 371.44 |
| | MuHiSSE1 | -161.98 | 351.96 | 364.31 |
| | MuHiSSE2 | -161.99 | 351.97 | 364.33 |
| | MuHiSSE3 | -159.59 | 371.18 | 435.00 |
| | MuCID2 | -171.10 | 366.19 | 374.86 |
| Tree52 | Constant rate | -170.61 | 361.22 | 367.01 |
| | NoHidden1 | -160.59 | 343.18 | 350.31 |
| | **NoHidden2** | **-158.94** | **339.88** | **347.01** |
| | NoHidden3 | -158.20 | 342.40 | 352.80 |
| | MuHiSSE1 | -157.48 | 342.97 | 355.32 |
| | MuHiSSE2 | -157.77 | 343.54 | 355.90 |
| | MuHiSSE3 | -156.13 | 364.25 | 428.07 |

| | | | | |
|---|---|---|---|---|
| | MuCID2 | -164.73 | 353.46 | 362.12 |
| Tree53 | Constant rate | -171.49 | 362.99 | 368.78 |
| | NoHidden1 | -158.49 | 338.98 | 346.11 |
| | **NoHidden2** | **-155.43** | **332.86** | **340.00** |
| | NoHidden3 | -155.51 | 337.01 | 347.41 |
| | MuHiSSE1 | -164.83 | 357.66 | 370.02 |
| | MuHiSSE2 | -153.57 | 335.15 | 347.50 |
| | MuHiSSE3 | -150.96 | 353.91 | 417.73 |
| | MuCID2 | -161.74 | 347.47 | 356.14 |
| Tree54 | Constant rate | -172.36 | 364.73 | 370.52 |
| | NoHidden1 | -159.70 | 341.41 | 348.54 |
| | **NoHidden2** | **-155.73** | **333.47** | **340.60** |
| | NoHidden3 | -154.00 | 333.99 | 344.39 |
| | MuHiSSE1 | -153.72 | 335.45 | 347.80 |
| | MuHiSSE2 | -155.37 | 338.74 | 351.09 |
| | MuHiSSE3 | -151.16 | 354.32 | 418.14 |
| | MuCID2 | -162.62 | 349.25 | 357.92 |
| Tree55 | Constant rate | -177.82 | 375.63 | 381.42 |
| | NoHidden1 | -166.35 | 354.70 | 361.83 |
| | **NoHidden2** | **-165.45** | **352.91** | **360.04** |
| | NoHidden3 | -166.35 | 358.70 | 369.10 |
| | MuHiSSE1 | -159.29 | 346.58 | 358.93 |
| | MuHiSSE2 | -160.62 | 349.25 | 361.60 |
| | MuHiSSE3 | -161.78 | 375.56 | 439.37 |
| | MuCID2 | -169.90 | 363.81 | 372.47 |
| Tree56 | Constant rate | -179.25 | 378.50 | 384.29 |
| | NoHidden1 | -166.03 | 354.05 | 361.19 |
| | **NoHidden2** | **-156.01** | **334.03** | **341.16** |
| | NoHidden3 | -155.14 | 336.29 | 346.69 |
| | MuHiSSE1 | -165.23 | 358.46 | 370.82 |
| | MuHiSSE2 | -159.57 | 347.14 | 359.50 |
| | MuHiSSE3 | -158.17 | 368.34 | 432.16 |
| | MuCID2 | -169.87 | 363.74 | 372.40 |
| Tree57 | Constant rate | -172.03 | 364.07 | 369.86 |
| | NoHidden1 | -159.96 | 341.92 | 349.05 |
| | **NoHidden2** | **-154.74** | **331.49** | **338.62** |
| | NoHidden3 | -152.58 | 331.17 | 341.57 |
| | MuHiSSE1 | -165.16 | 358.32 | 370.67 |
| | MuHiSSE2 | -155.94 | 339.88 | 352.23 |
| | MuHiSSE3 | -152.76 | 357.52 | 421.34 |
| | MuCID2 | -163.48 | 350.95 | 359.62 |
| Tree58 | Constant rate | -178.53 | 377.05 | 382.84 |
| | NoHidden1 | -165.28 | 352.56 | 359.70 |
| | **NoHidden2** | **-157.66** | **337.33** | **344.46** |
| | NoHidden3 | -165.12 | 356.24 | 366.64 |
| | MuHiSSE1 | -158.23 | 344.45 | 356.81 |
| | MuHiSSE2 | -160.17 | 348.35 | 360.70 |
| | MuHiSSE3 | -158.90 | 369.81 | 433.62 |
| | MuCID2 | -169.54 | 363.08 | 371.75 |
| Tree59 | Constant rate | -179.10 | 378.20 | 383.99 |
| | NoHidden1 | -158.08 | 338.17 | 345.30 |
| | **NoHidden2** | **-154.35** | **330.70** | **337.83** |
| | NoHidden3 | -151.95 | 329.90 | 340.30 |

| | | | | |
|---|---|---|---|---|
| [illegible] | MuHiSSE1 | -153.55 | 335.09 | 347.44 |
| | MuHiSSE2 | -159.25 | 346.49 | 358.85 |
| | MuHiSSE3 | -150.45 | 352.90 | 416.71 |
| | MuCID2 | -170.37 | 364.74 | 373.41 |
| Tree60 | Constant rate | -181.40 | 382.81 | 388.60 |
| | NoHidden1 | -170.56 | 363.12 | 370.25 |
| | **NoHidden2** | **-162.88** | **347.75** | **354.89** |
| | NoHidden3 | -162.13 | 350.26 | 360.66 |
| | MuHiSSE1 | -174.08 | 376.17 | 388.52 |
| | MuHiSSE2 | -165.87 | 359.73 | 372.09 |
| | MuHiSSE3 | -162.16 | 376.33 | 440.15 |
| | MuCID2 | -174.25 | 372.50 | 381.16 |
| Tree61 | Constant rate | -175.15 | 370.29 | 376.08 |
| | NoHidden1 | -164.03 | 350.07 | 357.20 |
| | NoHidden2 | -157.52 | 337.05 | 344.18 |
| | **NoHidden3** | **-152.52** | **331.04** | **341.44** |
| | MuHiSSE1 | -157.52 | 343.03 | 355.39 |
| | MuHiSSE2 | -159.82 | 347.64 | 359.99 |
| | MuHiSSE3 | -157.58 | 367.15 | 430.97 |
| | MuCID2 | -168.08 | 360.16 | 368.83 |
| Tree62 | Constant rate | -178.07 | 376.15 | 381.94 |
| | NoHidden1 | -167.14 | 356.29 | 363.42 |
| | NoHidden2 | -166.25 | 354.50 | 361.64 |
| | **NoHidden3** | **-157.91** | **341.81** | **352.21** |
| | MuHiSSE1 | -162.94 | 353.89 | 366.24 |
| | MuHiSSE2 | -162.83 | 353.66 | 366.02 |
| | MuHiSSE3 | -157.28 | 366.56 | 430.38 |
| | MuCID2 | -170.62 | 365.25 | 373.92 |
| Tree63 | Constant rate | -170.56 | 361.12 | 366.91 |
| | NoHidden1 | -158.54 | 339.08 | 346.22 |
| | **NoHidden2** | **-153.25** | **328.49** | **335.63** |
| | NoHidden3 | -153.54 | 333.08 | 343.48 |
| | MuHiSSE1 | -160.81 | 349.62 | 361.98 |
| | MuHiSSE2 | -154.56 | 337.11 | 349.46 |
| | MuHiSSE3 | -151.98 | 355.97 | 419.78 |
| | MuCID2 | -161.19 | 346.39 | 355.05 |
| Tree64 | Constant rate | -177.18 | 374.35 | 380.14 |
| | NoHidden1 | -166.69 | 355.38 | 362.51 |
| | **NoHidden2** | **-161.41** | **344.83** | **351.96** |
| | NoHidden3 | -166.42 | 358.83 | 369.23 |
| | MuHiSSE1 | -160.06 | 348.11 | 360.46 |
| | MuHiSSE2 | -162.41 | 352.82 | 365.17 |
| | MuHiSSE3 | -156.82 | 365.63 | 429.45 |
| | MuCID2 | -169.22 | 362.45 | 371.11 |
| Tree65 | Constant rate | -173.37 | 366.74 | 372.53 |
| | NoHidden1 | -162.90 | 347.79 | 354.93 |
| | **NoHidden2** | **-158.53** | **339.07** | **346.20** |
| | NoHidden3 | -160.41 | 346.83 | 357.23 |
| | MuHiSSE1 | -157.48 | 342.96 | 355.31 |
| | MuHiSSE2 | -158.56 | 345.12 | 357.47 |
| | MuHiSSE3 | -152.57 | 357.14 | 420.95 |
| | MuCID2 | -166.40 | 356.80 | 365.47 |
| | Constant rate | -172.86 | 365.72 | 371.51 |

| | | | | |
|---|---|---|---|---|
| Tree66 | NoHidden1 | -160.77 | 343.55 | 350.68 |
| | NoHidden2 | -156.71 | 335.42 | 342.55 |
| | **NoHidden3** | **-152.19** | **330.37** | **340.77** |
| | MuHiSSE1 | -156.08 | 340.16 | 352.52 |
| | MuHiSSE2 | -157.43 | 342.86 | 355.21 |
| | MuHiSSE3 | -155.10 | 362.20 | 426.02 |
| | MuCID2 | -164.11 | 352.23 | 360.90 |
| Tree67 | Constant rate | -179.76 | 379.52 | 385.31 |
| | NoHidden1 | -167.80 | 357.60 | 364.73 |
| | **NoHidden2** | **-165.63** | **353.25** | **360.39** |
| | NoHidden3 | -167.16 | 360.33 | 370.73 |
| | MuHiSSE1 | -162.84 | 353.69 | 366.04 |
| | MuHiSSE2 | -161.84 | 351.68 | 364.03 |
| | MuHiSSE3 | -159.03 | 370.06 | 433.88 |
| | MuCID2 | -171.67 | 367.35 | 376.02 |
| Tree68 | Constant rate | -170.71 | 361.42 | 367.21 |
| | NoHidden1 | -158.89 | 339.79 | 346.92 |
| | **NoHidden2** | **-152.62** | **327.23** | **334.37** |
| | NoHidden3 | -149.54 | 325.09 | 335.49 |
| | MuHiSSE1 | -163.87 | 355.74 | 368.09 |
| | MuHiSSE2 | -155.25 | 338.50 | 350.85 |
| | MuHiSSE3 | -148.68 | 349.35 | 413.17 |
| | MuCID2 | -162.86 | 349.72 | 358.38 |
| Tree69 | Constant rate | -180.69 | 381.37 | 387.16 |
| | NoHidden1 | -167.22 | 356.45 | 363.58 |
| | **NoHidden2** | **-160.31** | **342.61** | **349.75** |
| | NoHidden3 | -159.44 | 344.88 | 355.28 |
| | MuHiSSE1 | -160.34 | 348.67 | 361.03 |
| | MuHiSSE2 | -161.85 | 351.70 | 364.05 |
| | MuHiSSE3 | -160.83 | 373.66 | 437.48 |
| | MuCID2 | -170.72 | 365.45 | 374.11 |
| Tree70 | Constant rate | -173.88 | 367.77 | 373.56 |
| | NoHidden1 | -162.58 | 347.16 | 354.30 |
| | **NoHidden2** | **-155.29** | **332.59** | **339.73** |
| | NoHidden3 | -155.58 | 337.15 | 347.55 |
| | MuHiSSE1 | -155.27 | 338.54 | 350.89 |
| | MuHiSSE2 | -157.43 | 342.85 | 355.20 |
| | MuHiSSE3 | -150.22 | 352.44 | 416.26 |
| | MuCID2 | -166.20 | 356.40 | 365.07 |
| Tree71 | Constant rate | -178.81 | 377.62 | 383.41 |
| | NoHidden1 | -165.07 | 352.14 | 359.28 |
| | **NoHidden2** | **-163.37** | **348.75** | **355.88** |
| | NoHidden3 | -163.34 | 352.68 | 363.08 |
| | MuHiSSE1 | -166.23 | 360.46 | 372.81 |
| | MuHiSSE2 | -160.27 | 348.54 | 360.89 |
| | MuHiSSE3 | -157.32 | 366.64 | 430.46 |
| | MuCID2 | -167.97 | 359.95 | 368.61 |
| Tree72 | Constant rate | -170.48 | 360.96 | 366.75 |
| | NoHidden1 | -158.76 | 339.52 | 346.65 |
| | NoHidden2 | -157.54 | 337.07 | 344.21 |
| | **NoHidden3** | **-149.96** | **325.92** | **336.32** |
| | MuHiSSE1 | -154.70 | 337.40 | 349.75 |
| | MuHiSSE2 | -153.14 | 334.29 | 346.64 |

| | | | | |
|---|---|---|---|---|
| | MuHiSSE3 | -154.29 | 360.59 | 424.41 |
| | MuCID2 | -162.83 | 349.66 | 358.33 |
| Tree73 | Constant rate | -175.53 | 371.05 | 376.84 |
| | NoHidden1 | -163.89 | 349.78 | 356.92 |
| | **NoHidden2** | **-158.73** | **339.45** | **346.59** |
| | NoHidden3 | -162.25 | 350.50 | 360.90 |
| | MuHiSSE1 | -158.31 | 344.63 | 356.98 |
| | MuHiSSE2 | -160.13 | 348.26 | 360.61 |
| | MuHiSSE3 | -158.20 | 368.39 | 432.21 |
| | MuCID2 | -166.75 | 357.50 | 366.16 |
| Tree74 | Constant rate | -174.73 | 369.45 | 375.24 |
| | NoHidden1 | -162.43 | 346.86 | 354.00 |
| | **NoHidden2** | **-154.33** | **330.65** | **337.79** |
| | NoHidden3 | -162.17 | 350.34 | 360.74 |
| | MuHiSSE1 | -156.84 | 341.68 | 354.03 |
| | MuHiSSE2 | -156.86 | 341.71 | 354.06 |
| | MuHiSSE3 | -150.04 | 352.08 | 415.90 |
| | MuCID2 | -165.59 | 355.17 | 363.84 |
| Tree75 | Constant rate | -176.74 | 373.47 | 379.26 |
| | NoHidden1 | -167.32 | 356.64 | 363.77 |
| | **NoHidden2** | **-159.69** | **341.38** | **348.52** |
| | NoHidden3 | -166.96 | 359.91 | 370.31 |
| | MuHiSSE1 | -159.98 | 347.96 | 360.31 |
| | MuHiSSE2 | -162.39 | 352.79 | 365.14 |
| | MuHiSSE3 | -160.55 | 373.10 | 436.92 |
| | MuCID2 | -170.61 | 365.22 | 373.88 |
| Tree76 | Constant rate | -179.28 | 378.57 | 384.35 |
| | NoHidden1 | -163.79 | 349.57 | 356.71 |
| | **NoHidden2** | **-155.10** | **332.20** | **339.34** |
| | NoHidden3 | -163.78 | 353.56 | 363.96 |
| | MuHiSSE1 | -153.47 | 334.93 | 347.29 |
| | MuHiSSE2 | -157.33 | 342.67 | 355.02 |
| | MuHiSSE3 | -150.34 | 352.68 | 416.50 |
| | MuCID2 | -167.37 | 358.74 | 367.40 |
| Tree77 | Constant rate | -165.66 | 351.31 | 357.10 |
| | NoHidden1 | -152.77 | 327.54 | 334.67 |
| | **NoHidden2** | **-149.34** | **320.68** | **327.82** |
| | NoHidden3 | -149.98 | 325.96 | 336.36 |
| | MuHiSSE1 | -147.45 | 322.90 | 335.25 |
| | MuHiSSE2 | -148.43 | 324.86 | 337.22 |
| | MuHiSSE3 | -146.56 | 345.12 | 408.93 |
| | MuCID2 | -156.05 | 336.10 | 344.76 |
| Tree78 | Constant rate | -176.00 | 372.00 | 377.79 |
| | NoHidden1 | -166.61 | 355.21 | 362.35 |
| | **NoHidden2** | **-164.60** | **351.21** | **358.34** |
| | NoHidden3 | -166.15 | 358.30 | 368.70 |
| | MuHiSSE1 | -161.15 | 350.30 | 362.65 |
| | MuHiSSE2 | -161.96 | 351.92 | 364.27 |
| | MuHiSSE3 | -161.08 | 374.17 | 437.99 |
| | MuCID2 | -169.69 | 363.39 | 372.05 |
| | Constant rate | -178.82 | 377.64 | 383.43 |
| | NoHidden1 | -168.30 | 358.60 | 365.73 |
| | **NoHidden2** | **-154.56** | **331.12** | **338.26** |

| | | | | |
|---|---|---|---|---|
| Tree79 | NoHidden3 | -168.01 | 362.01 | 372.41 |
| | MuHiSSE1 | -155.58 | 339.16 | 351.52 |
| | MuHiSSE2 | -160.61 | 349.21 | 361.57 |
| | MuHiSSE3 | -159.34 | 370.68 | 434.50 |
| | MuCID2 | -170.73 | 365.47 | 374.13 |
| Tree80 | Constant rate | -172.57 | 365.14 | 370.93 |
| | NoHidden1 | -162.72 | 347.44 | 354.58 |
| | **NoHidden2** | **-160.23** | **342.46** | **349.59** |
| | NoHidden3 | -162.04 | 350.09 | 360.49 |
| | MuHiSSE1 | -162.60 | 353.21 | 365.56 |
| | MuHiSSE2 | -157.49 | 342.99 | 355.34 |
| | MuHiSSE3 | -156.01 | 364.03 | 427.84 |
| | MuCID2 | -166.59 | 357.17 | 365.84 |
| Tree81 | Constant rate | -175.24 | 370.48 | 376.27 |
| | NoHidden1 | -163.20 | 348.39 | 355.53 |
| | **NoHidden2** | **-157.10** | **336.19** | **343.33** |
| | NoHidden3 | -162.19 | 350.38 | 360.78 |
| | MuHiSSE1 | -155.72 | 339.44 | 351.79 |
| | MuHiSSE2 | -158.37 | 344.75 | 357.10 |
| | MuHiSSE3 | -155.00 | 362.00 | 425.81 |
| | MuCID2 | -166.91 | 357.83 | 366.49 |
| Tree82 | Constant rate | -177.83 | 375.65 | 381.44 |
| | NoHidden1 | -164.18 | 350.35 | 357.49 |
| | NoHidden2 | -163.00 | 348.00 | 355.13 |
| | **NoHidden3** | **-152.95** | **331.90** | **342.30** |
| | MuHiSSE1 | -164.25 | 356.51 | 368.86 |
| | MuHiSSE2 | -158.76 | 345.51 | 357.86 |
| | MuHiSSE3 | -153.97 | 359.95 | 423.77 |
| | MuCID2 | -168.86 | 361.72 | 370.39 |
| Tree83 | Constant rate | -175.36 | 370.71 | 376.50 |
| | NoHidden1 | -164.99 | 351.98 | 359.12 |
| | **NoHidden2** | **-160.50** | **343.01** | **350.14** |
| | NoHidden3 | -160.11 | 346.22 | 356.62 |
| | MuHiSSE1 | -159.79 | 347.59 | 359.94 |
| | MuHiSSE2 | -160.79 | 349.58 | 361.93 |
| | MuHiSSE3 | -159.63 | 371.26 | 435.08 |
| | MuCID2 | -168.25 | 360.50 | 369.16 |
| Tree84 | Constant rate | -171.52 | 363.04 | 368.83 |
| | NoHidden1 | -156.21 | 334.42 | 341.55 |
| | NoHidden2 | -149.31 | 320.63 | 327.76 |
| | **NoHidden3** | **-144.02** | **314.04** | **324.44** |
| | MuHiSSE1 | -148.26 | 324.52 | 336.88 |
| | MuHiSSE2 | -151.20 | 330.40 | 342.75 |
| | MuHiSSE3 | -145.66 | 343.33 | 407.15 |
| | MuCID2 | -158.88 | 341.76 | 350.43 |
| Tree85 | Constant rate | -171.60 | 363.20 | 368.99 |
| | NoHidden1 | -158.71 | 339.41 | 346.55 |
| | **NoHidden2** | **-153.75** | **329.50** | **336.64** |
| | NoHidden3 | -153.45 | 332.91 | 343.31 |
| | MuHiSSE1 | -153.59 | 335.17 | 347.52 |
| | MuHiSSE2 | -153.52 | 335.03 | 347.38 |
| | MuHiSSE3 | -145.79 | 343.58 | 407.40 |
| | MuCID2 | -162.64 | 349.27 | 357.94 |

| | | | | |
|---|---|---|---|---|
| Tree86 | Constant rate | -168.76 | 357.51 | 363.30 |
| | NoHidden1 | -158.87 | 339.75 | 346.88 |
| | **NoHidden2** | **-156.12** | **334.23** | **341.37** |
| | NoHidden3 | -155.83 | 337.66 | 348.06 |
| | MuHiSSE1 | -154.96 | 337.91 | 350.26 |
| | MuHiSSE2 | -154.61 | 337.21 | 349.57 |
| | MuHiSSE3 | -152.21 | 356.41 | 420.23 |
| | MuCID2 | -162.46 | 348.92 | 357.59 |
| Tree87 | Constant rate | -178.62 | 377.25 | 383.03 |
| | NoHidden1 | -168.11 | 358.23 | 365.36 |
| | **NoHidden2** | **-159.93** | **341.86** | **348.99** |
| | NoHidden3 | -159.79 | 345.59 | 355.99 |
| | MuHiSSE1 | -163.40 | 354.80 | 367.16 |
| | MuHiSSE2 | -163.31 | 354.63 | 366.98 |
| | MuHiSSE3 | -157.96 | 367.93 | 431.75 |
| | MuCID2 | -170.87 | 365.74 | 374.41 |
| Tree88 | Constant rate | -166.29 | 352.58 | 358.37 |
| | NoHidden1 | -154.77 | 331.54 | 338.68 |
| | **NoHidden2** | **-152.94** | **327.89** | **335.02** |
| | NoHidden3 | -154.77 | 335.53 | 345.93 |
| | MuHiSSE1 | -148.96 | 325.92 | 338.28 |
| | MuHiSSE2 | -152.24 | 332.49 | 344.84 |
| | MuHiSSE3 | -149.59 | 351.19 | 415.01 |
| | MuCID2 | -158.47 | 340.94 | 349.61 |
| Tree89 | Constant rate | -180.77 | 381.54 | 387.33 |
| | NoHidden1 | -170.04 | 362.07 | 369.21 |
| | **NoHidden2** | **-163.52** | **349.05** | **356.18** |
| | NoHidden3 | -169.80 | 365.60 | 376.00 |
| | MuHiSSE1 | -164.18 | 356.36 | 368.71 |
| | MuHiSSE2 | -165.14 | 358.28 | 370.63 |
| | MuHiSSE3 | -157.84 | 367.67 | 431.49 |
| | MuCID2 | -172.43 | 368.87 | 377.54 |
| Tree90 | Constant rate | -176.48 | 372.97 | 378.76 |
| | NoHidden1 | -164.32 | 350.63 | 357.77 |
| | **NoHidden2** | **-159.55** | **341.09** | **348.23** |
| | NoHidden3 | -158.17 | 342.34 | 352.74 |
| | MuHiSSE1 | -167.53 | 363.07 | 375.42 |
| | MuHiSSE2 | -160.01 | 348.02 | 360.37 |
| | MuHiSSE3 | -157.13 | 366.25 | 430.07 |
| | MuCID2 | -167.77 | 359.53 | 368.20 |
| Tree91 | Constant rate | -171.28 | 362.56 | 368.35 |
| | NoHidden1 | -160.10 | 342.20 | 349.34 |
| | NoHidden2 | -159.08 | 340.17 | 347.30 |
| | **NoHidden3** | **-151.17** | **328.34** | **338.74** |
| | MuHiSSE1 | -153.91 | 335.82 | 348.17 |
| | MuHiSSE2 | -155.04 | 338.07 | 350.42 |
| | MuHiSSE3 | -152.61 | 357.22 | 421.04 |
| | MuCID2 | -162.61 | 349.22 | 357.89 |
| Tree92 | Constant rate | -171.43 | 362.87 | 368.66 |
| | NoHidden1 | -161.35 | 344.70 | 351.83 |
| | **NoHidden2** | **-149.87** | **321.74** | **328.88** |
| | NoHidden3 | -149.14 | 324.27 | 334.67 |
| | MuHiSSE1 | -150.53 | 329.06 | 341.41 |

| | | | | |
|---|---|---|---|---|
| | MuHiSSE2 | -155.18 | 338.37 | 350.72 |
| | MuHiSSE3 | -147.83 | 347.65 | 411.47 |
| | MuCID2 | -165.19 | 354.38 | 363.04 |
| Tree93 | Constant rate | -169.61 | 359.22 | 365.01 |
| | NoHidden1 | -158.54 | 339.08 | 346.21 |
| | **NoHidden2** | **-153.41** | **328.81** | **335.95** |
| | NoHidden3 | -152.34 | 330.67 | 341.07 |
| | MuHiSSE1 | -152.45 | 332.90 | 345.25 |
| | MuHiSSE2 | -154.30 | 336.60 | 348.96 |
| | MuHiSSE3 | -152.38 | 356.77 | 420.59 |
| | MuCID2 | -162.80 | 349.61 | 358.27 |
| Tree94 | Constant rate | -163.67 | 347.34 | 353.13 |
| | NoHidden1 | -154.76 | 331.52 | 338.66 |
| | **NoHidden2** | **-154.60** | **331.20** | **338.34** |
| | NoHidden3 | -154.15 | 334.31 | 344.71 |
| | MuHiSSE1 | -151.08 | 330.16 | 342.52 |
| | MuHiSSE2 | -152.79 | 333.57 | 345.93 |
| | MuHiSSE3 | -150.71 | 353.42 | 417.24 |
| | MuCID2 | -158.83 | 341.66 | 350.32 |
| Tree95 | Constant rate | -177.81 | 375.63 | 381.41 |
| | NoHidden1 | -167.29 | 356.59 | 363.72 |
| | **NoHidden2** | **-158.63** | **339.26** | **346.39** |
| | NoHidden3 | -158.03 | 342.05 | 352.45 |
| | MuHiSSE1 | -158.32 | 344.64 | 356.99 |
| | MuHiSSE2 | -160.80 | 349.59 | 361.94 |
| | MuHiSSE3 | -157.26 | 366.52 | 430.34 |
| | MuCID2 | -171.27 | 366.54 | 375.20 |
| Tree96 | Constant rate | -171.24 | 362.48 | 368.27 |
| | NoHidden1 | -161.39 | 344.78 | 351.91 |
| | **NoHidden2** | **-152.87** | **327.74** | **334.88** |
| | NoHidden3 | -161.26 | 348.51 | 358.91 |
| | MuHiSSE1 | -154.60 | 337.19 | 349.55 |
| | MuHiSSE2 | -156.37 | 340.73 | 353.08 |
| | MuHiSSE3 | -151.79 | 355.58 | 419.40 |
| | MuCID2 | -163.77 | 351.55 | 360.21 |
| Tree97 | Constant rate | -177.98 | 375.97 | 381.76 |
| | NoHidden1 | -165.65 | 353.29 | 360.43 |
| | **NoHidden2** | **-156.30** | **334.59** | **341.73** |
| | NoHidden3 | -165.53 | 357.06 | 367.46 |
| | MuHiSSE1 | -156.40 | 340.80 | 353.16 |
| | MuHiSSE2 | -160.98 | 349.95 | 362.31 |
| | MuHiSSE3 | -159.41 | 370.82 | 434.63 |
| | MuCID2 | -169.31 | 362.63 | 371.30 |
| Tree98 | Constant rate | -169.92 | 359.84 | 365.63 |
| | NoHidden1 | -155.84 | 333.68 | 340.82 |
| | **NoHidden2** | **-149.62** | **321.25** | **328.38** |
| | NoHidden3 | -155.08 | 336.15 | 346.55 |
| | MuHiSSE1 | -147.38 | 322.77 | 335.12 |
| | MuHiSSE2 | -149.89 | 327.77 | 340.13 |
| | MuHiSSE3 | -146.50 | 344.99 | 408.81 |
| | MuCID2 | -158.53 | 341.05 | 349.72 |
| | Constant rate | -175.40 | 370.80 | 376.59 |
| | NoHidden1 | -165.62 | 353.24 | 360.37 |

| | | | | |
|---|---|---|---|---|
| Tree99 | NoHidden2 | -164.17 | 350.33 | 357.47 |
| | **NoHidden3** | **-157.86** | **341.73** | **352.13** |
| | MuHiSSE1 | -161.06 | 350.12 | 362.47 |
| | MuHiSSE2 | -160.77 | 349.54 | 361.89 |
| | MuHiSSE3 | -157.90 | 367.79 | 431.61 |
| | MuCID2 | -168.18 | 360.36 | 369.02 |
| Tree100 | Constant rate | -173.37 | 366.74 | 372.53 |
| | NoHidden1 | -160.62 | 343.25 | 350.38 |
| | **NoHidden2** | **-156.19** | **334.39** | **341.52** |
| | NoHidden3 | -153.43 | 332.86 | 343.26 |
| | MuHiSSE1 | -154.08 | 336.16 | 348.51 |
| | MuHiSSE2 | -157.13 | 342.25 | 354.61 |
| | MuHiSSE3 | -151.95 | 355.89 | 419.71 |
| | MuCID2 | -164.20 | 352.39 | 361.06 |
